# Molecular dynamics and in silico mutagenesis on the reversible inhibitor-bound SARS-CoV-2 Main Protease complexes reveal the role of a lateral pocket in enhancing the ligand affinity

**DOI:** 10.1101/2020.10.31.363309

**Authors:** Ying Li Weng, Shiv Rakesh Naik, Nadia Dingelstad, Subha Kalyaanamoorthy, Aravindhan Ganesan

**Affiliations:** ArGan’s Lab, School of Pharmacy, Faculty of Science, University of Waterloo, Ontario, Canada; Department of Chemistry, Faculty of Science, University of Waterloo, Ontario, Canada

**Keywords:** SARS-CoV-2, main protease, novel coronavirus, molecular dynamics, in silico mutagenesis

## Abstract

The 2019 novel coronavirus pandemic caused by SARS-CoV-2 remains a serious health threat to humans and a number of countries are already in the middle of the second wave of infection. There is an urgent need to develop therapeutics against this deadly virus. Recent scientific evidences have suggested that the main protease (M^pro^) enzyme in SARS-CoV-2 can be an ideal drug target due to its crucial role in the viral replication and transcription processes. Therefore, there are ongoing research efforts to identify drug candidates against SARS-CoV-2 M^pro^ that resulted in hundreds of X-ray crystal structures of ligand bound M^pro^ complexes in the protein data bank (PDB) that describe structural details of different chemotypes of fragments binding within different sites in M^pro^. In this work, we perform rigorous molecular dynamics (MD) simulation of 62 reversible ligand-M^pro^ complexes in the PDB to gain mechanistic insights about their interactions at atomic level. Using a total of ~2.25 μs long MD trajectories, we identified and characterized different pockets and their conformational dynamics in the apo M^pro^ structure. Later, using the published PDB structures, we analyzed the dynamic interactions and binding affinity of small ligands within those pockets. Our results identified the key residues that stabilize the ligands in the catalytic sites and other pockets in M^pro^. Our analyses unraveled the role of a lateral pocket in the catalytic site in M^pro^ that is critical for enhancing the ligand binding to the enzyme. We also highlighted the important contribution from HIS163 in this lateral pocket towards ligand binding and affinity against M^pro^ through computational mutation analyses. Further, we revealed the effects of explicit water molecules and M^pro^ dimerization in the ligand association with the target. Thus, comprehensive molecular level insights gained from this work can be useful to identify or design potent small molecule inhibitors against SARS-CoV-2 M^pro^.

## Introduction

The novel coronavirus, severe acute respiratory syndrome coronavirus 2 (or SARS-CoV-2), was declared a global pandemic on March 11, 2020^1^ and as of October 27, this outbreak has already caused over 43 million cases and 1.16 million deaths worldwide. While efforts for developing a potent vaccine against this virus are occurring at an accelerated pace, there are currently no clinically approved drug or vaccines to provide a cure or immunity from this deadly virus. Furthermore, the rapid adaptive nature of the virus and its ability to mutate^2–5^ present new challenges in developing effective novel antiviral therapy. Thus, understanding the specific drug targets of SARS-CoV-2 ^6,7^, clarifying its mechanism of virulence^8,9^, searching for a repurposable drug and development of a small molecule drug^10–14^ or a vaccine^15–17^ against SARS-CoV-2 have since been the main focus of scientific research.

SARS-CoV-2 is an enveloped, positive-sense, single-stranded beta-coronavirus that exhibits significant sequence and structure homology to that of the previous coronaviruses strains such SARS-CoV-1 (generally known as SARS-CoV) and Middle Eastern Respiratory Syndrome coronavirus (MERS-CoV)^1^. Amino acid sequence analyses revealed that SARS-CoV-2 shared ~79% sequence identity with SARS-CoV-1 and ~50% similarity to MERS^18^. Much like the previously identified coronaviruses, SARS-Cov-2 is composed of structural proteins that are important in producing a complete viral particle, non-structural proteins that act as enzymes or transcription/replication factors in the viral life cycle, and numerous accessory proteins^19,20^. The structural proteins include the spike, envelope, nucleocapsid, and membrane proteins, which have been identified as targets in recent publications concerning SARS-CoV-2 drug discovery^19,20^. There are 16 non-structural proteins (nsps) including the RNA-dependent RNA polymerase, helicase protein, papin-like protease (PL^pro^), and 3-chymotrypsin-like protease (also known as the main protease or M^pro^)[3]. The two cysteine proteases, PL^pro^ and M^pro^, process most of the other nsps. PL^pro^ cleaves nsp 1 to 3 and M^pro^ cleaves nsp 4-16 of the polyprotein at 11 cleavage cites ^14,21,22^.Amongst the other targets, the SARS-CoV-2 M^pro^ (or it will be interchangeably used with M^pro^ in the text hereafter) remains as an attractive therapeutic target as its role of cleaving the coronavirus polyproteins into functional components is vital for viral replication and survival^19,22,23^. Inhibition of the M^pro^ can block proteolytic enzyme activity on polyproteins, thereby, leaving a long peptide of polyprotein that is incapable of performing viral replication by itself^21,24^. In addition, the M^pro^ is highly conserved among coronaviruses. For example, the amino acid sequence of SARS-CoV-2 M^pro^ shares >95% similarity with that of SARS-CoV M^pro^ (sequence alignment provided in Fig. 1a). Therefore, there have been significant attraction towards targeting M^pro^ to control SARS-CoV-2 and there are currently hundreds of crystal structures of M^pro^ (either apo or in complex with ligands) reported in the protein data bank (PDB).

**Fig. 1:**
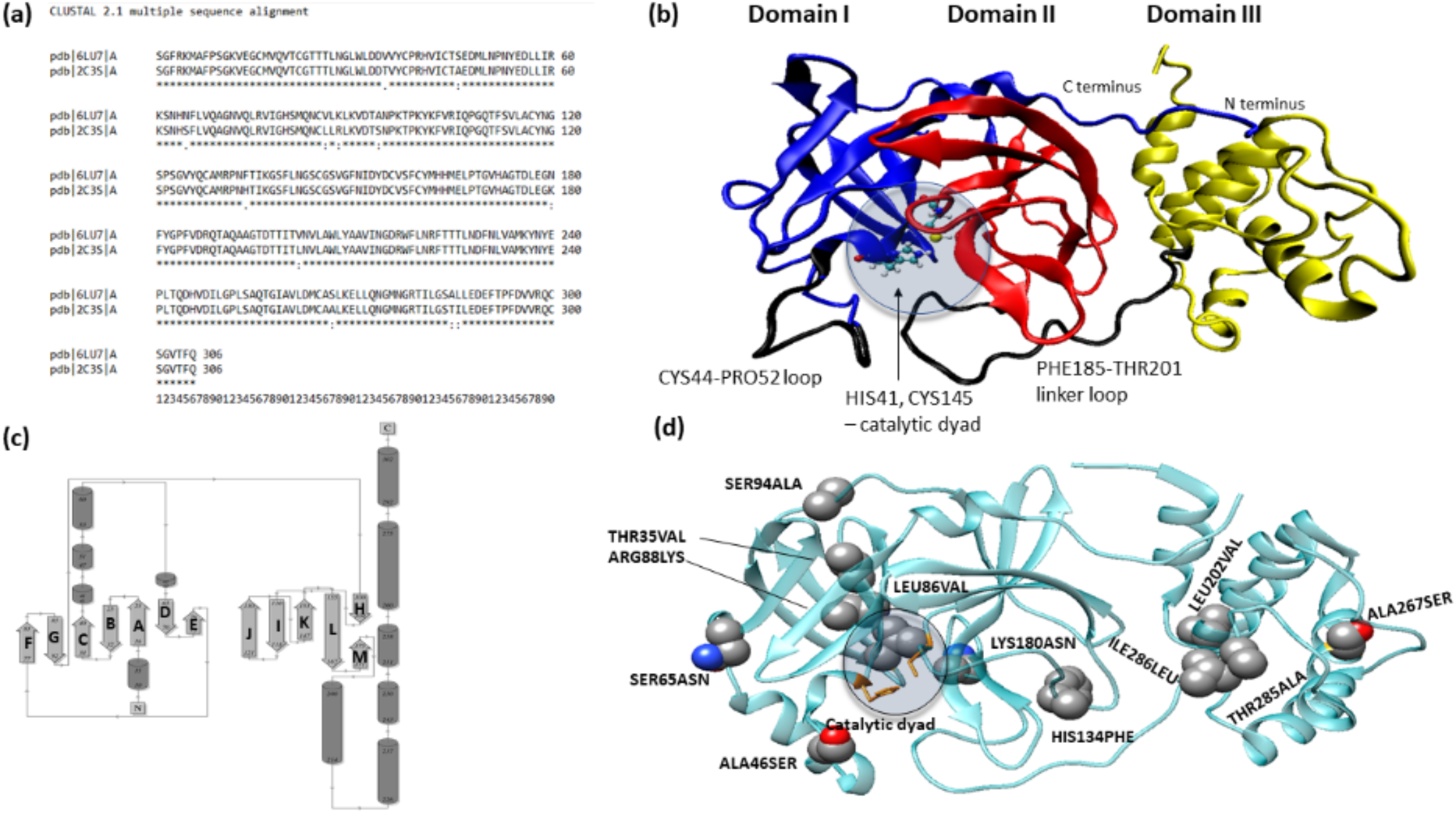
Sequence (a), structure (b), topology (c) of SARS-CoV-2 M^pro^ along with the comparison of its structure against SARS-CoV-1 M^pro^(d). **(a)** SARS-CoV-2 M^pro^ share 96% protein sequence similarity with SARS-CoV-1 M^pro^, as indicated by Clustal x2 (2.1) multiple sequence alignment. (b) A 3D structure of SARS-CoV-2 M^pro^ is shown with Domains I, II and III colored in blue, red and yellow respective. Two important loops close to the catalytic dyad (HIS41 and CYS145) is shown in black: CYS44-PRO52 loop flanks the catalytic dyad and PHE185-THR201 connects Domain II with Domain III. (c) A topology diagram showing the secondary structural elements in SARS-CoV-2 M^pro^ with β-sheets marked as A-M. (d) Structural alignment of SARS-CoV-2 M^pro^ and Sars-CoV-1 show only 12 mutations of amino acids that are shown as sphere representation.

Based on the known three-dimensional (3D) structure (in Fig. 1b), M^pro^ is composed of three domains including the N-terminal domain I (residues 1-101) domain II (residues 102-184) and the C-terminal domain III (residues 201-301)^25^. As seen in the topology diagram of M^pro^ in Fig. 1c, M^pro^ contains 13 β-sheets and 9 α-helices. domains I and II are mainly composed of antiparallel β-sheets, with domain I having seven antiparallel β-sheets (labelled A through G) and domain II consisting of the other six antiparallel β-sheets (labelled H through M in Fig. 1c). Unlike these domains, domain III does not have any β-sheet and is composed of a cluster of five α-helices. domains II and III are linked by a 17-residue long linker loop corresponding to the region PHE185-THR201 (Fig. 1b). The binding site cleft is formed at the interface of domains I and II with HIS41(from domain I) and CYS145 (from domain II) form a catalytic dyad^26^, which is unlike a catalytic triad in other cysteine and serine proteases^27,28^. It is proposed that a water molecule at the binding site plays the role of the third residue (in the catalytic triad) and is important in the catalytic mechanism of SARS-COV-2 M^pro 25,29^. A loop formed by CYS44-PRO52 in domain I and the domains II-III linker loop encase the active site as shown in Fig. 1b. Under physiological conditions, M^pro^ exists as a homodimer form, in which two monomers are placed in a perpendicular orientation to each other. In this orientation, the dimer interface is formed by the domain II of the first monomer particularly through residue GLU166 and the N-terminal residues of the second monomer (generally called the “N-finger”)^23^ enclosing a contact surface area of ~1390 Å^2 21^ The N finger from monomer 2 is critical as it closes the catalytic site in monomer A. Further, it is proposed that domain II is crucial in modulating the formation of dimer interface in M^pro30^. Comparison of the structure of SARS-CoV-2 Mpro against the SARS-CoV-1 Mpro (in Fig. 1d), show only amino acid variations in only 12 positions, which include THR35VAL, ALA46SER, SER65ASN, LEU86VAL, ARG88LYS, SER94ALA, HIS134PHE, LYS180ASN, LEU202VAL, ALA267SER, THR285ALA, ILE286LEU. With the exception of ALA46SER that is part of the CYS44-PRO52 loop in domain I, all the other mutations are away from the catalytic site and mostly on the surface regions on the three domains (Fig. 1b). However, as acknowledged by Bzówka et al., despite the high degree of sequence identify between the M^pro^ in SARS-CoV-1 and SARS-CoV-2, it is important to be mindful of the differences in the shape and sizes of their active site and the potential impact of these 12 mutations^25^. The availability of high-resolution crystal structures of Mpro has facilitated a number of *in silico-based* screening efforts to find suitable molecules from known drugs^18,31–33^ (for drug repurposing) or from the natural product libraries^29,34–37^. Apart from computational studies, the screening against M^pro^ was also carried out through experimental approaches^21,38–40^. For example, a screening performed using the fluorescence resonance energy transfer(FRET)-based enzymatic assay^39^ identified a number of inhibitors such as boceprevir, calpain inhibitors II and XIII, and GC-376 that inhibited M^pro^ with low micromolar IC50 values under the experimental conditions. Subsequently, the X-ray crystal structure of M^pro^ in complex with GC-376 was also reported in the PDB (PDB: 6WTT). Another structure-based design effort^38^ focussed on designing novel inhibitors from the chemical class of peptidomimetic α-ketoamides and identified a number of compounds that exhibited near-equipotency against the alphacoronaviruses, betacoronaviruses and enteroviruses when tested against the recombinant proteases, viral replicons and virus-infected cell cultures. Modifying a specific amide bond in an inhibitor identified from this work^38^ into a pyridone ring resulted in a novel α-ketoamide with improved pharmacokinetic properties (e.g., half-life of the inhibitor in plasma)^21^. The crystal structure of this new α-ketoamide inhibitor in complex with M^pro^ was also reported in the PDB (PDB ID: 6Y2F). One of the other significant researches against M^pro^ was reported by Douangamath et al^14^, in which rigorous fragment screening against SARS-CoV-2 M^pro^ resulted in several crystal structures of M^pro^ in complex with several covalent and non-covalent fragments. Most of the fragments screened were found to be bound within the catalytic site of M^pro^; however, a number of non-covalent fragments were bound at different other sites including the dimer interface site. These fragment-bound M^pro^ crystal structures^14^ provide a wealth of information about the ligand-M^pro^ interactions and also reveal several other cavities on the surface of M^pro^.

While a large number of experimentally resolved structures of ligand M^pro^ complexes are emerging rapidly, it might be important and useful to understand the dynamic interactions of these complexes and to garner any novel molecular level insights from the dynamic process. It is known that molecular dynamics (MD) simulation is a powerful tool to study the dynamic molecular recognition processes in biological systems such as ligand-protein and protein-protein complexes^41–45^. Earlier, MD has been successful in the discovery of a novel cryptic binding trench in HIV integrase enzyme^46^ that eventually led to the development of novel anti-HIV inhibitors such as raltegravir. Therefore, in this work, we use MD to characterize the dynamic interactions of almost all the crystal structures released as of June 10^th^, 2020 in PDB (the list is provided in the supplementary Table, ST1) and evaluate their binding free energies using the end-point molecular mechanics generalized Born surface area (MM-GBSA) approach. This includes a total of 62 ligand-bound complexes of M^pro^, whose 3D superposed structure is shown in Fig. 2 and the chemical structures of all the ligands are shown in supplementary Table, ST1. Due to the effective use of computational resources, we initially studied all the complexes with a monomer of M^pro^. It should be acknowledged that it has been suggested earlier that M^pro^ could be inactive in the monomer state^23,30^. However, since the activation process of an enzyme generally takes a much longer time scale than the practical limits of classical MD approach, the simulation of ligand bound to a monomer M^pro^ should be sufficient to capture the overall structural stability of the complexes and evaluate their binding affinity through end-point methods. Nevertheless, we selectively resimulated the complexes with a dimer form of M^pro^ when the ligand is either initially bound at (or at the proximity to) the dimer interface or when the ligand bound in the catalytic site unbound from the site during MD simulation. Each complex was subjected to a 30 ns long MD simulation which resulted in a total of ~2.25μs long MD trajectories. Our results highlight the significance of a lateral pocket in the ligand-Mpro affinity and, in particular, the key role of HIS163 in this pocket through computational mutagenesis. Further, we also calculated the binding free energy of the complexes (with ligand bound at the catalytic site) in the presence of explicit water molecules so as to unravel the effects of water molecules on the ligand-receptor affinity. Thus, our work should be useful to further our knowledge about the interactions of reversible ligand-SARS-CoV-2 M^pro^ complexes at molecular level.

**Fig. 2.**
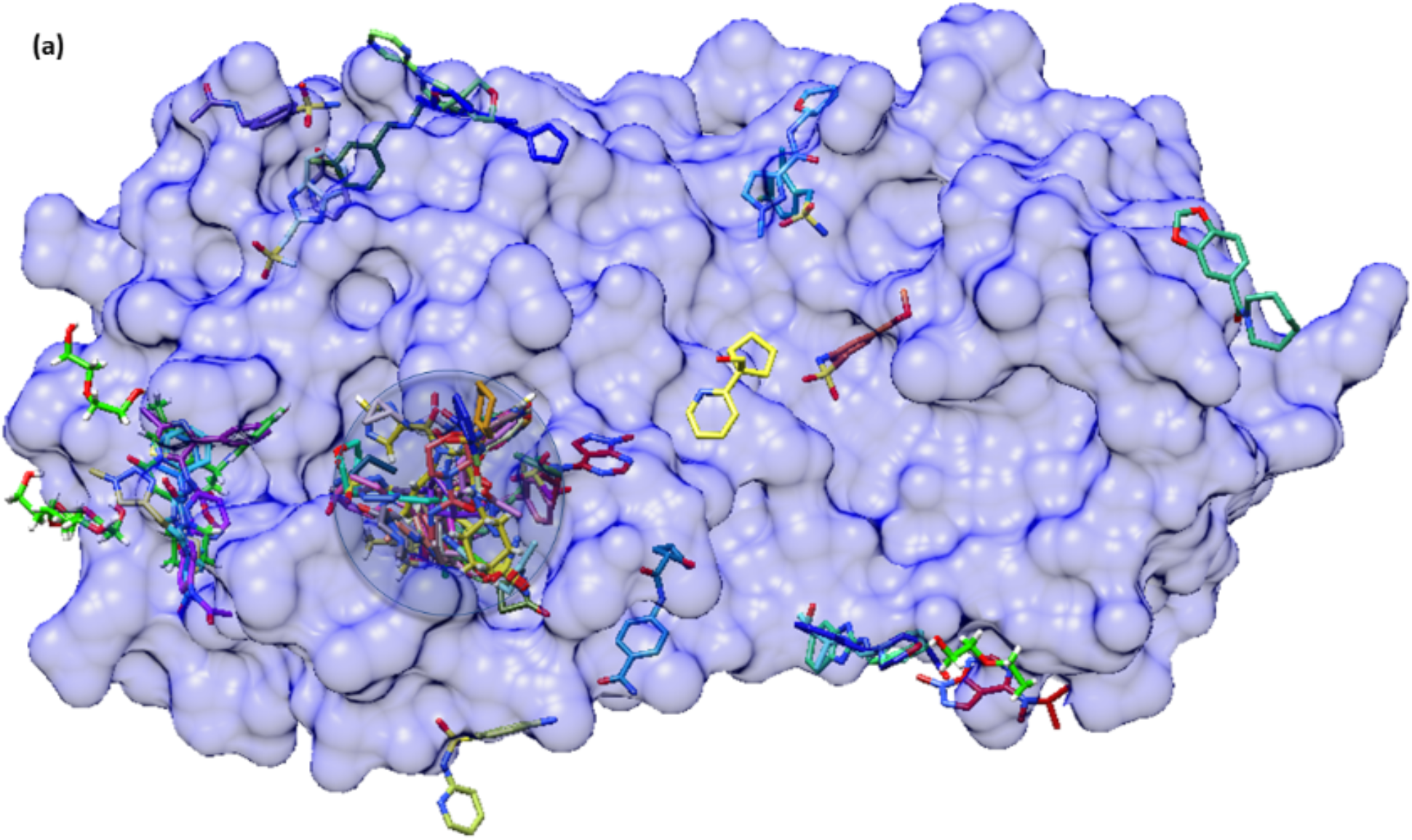
The binding of 62 reversible ligands (shown as stick representation) on different sites within SARS-CoV-2 M^pro^ (shown in Surf representation). Majority of the small molecules bound within the catalytic dyad site (shown in circle) and the rest bind at different surface sites on M^pro^.

## 2. Computational methods

### 2.1. Preparation of ligand-M^pro^ complexes for MD

A total of 62 experimentally resolved crystal structures of reversible small molecule-SARS-CoV-2 M^pro^ complexes (the list of PDB codes are provide in supplementary table ST1), along with the crystal structure of an apo Mpro (PDB: 6M2Q), were retrieved from PDB. Initially, all the DMS compounds, water molecules and ions were removed from the downloaded structures. For each complex, the AMBERff14SB force field^47^ and the general AMBER force field (GAFF2)^48^ were used for the protein and ligand, respectively. The partial charges of the ligand were assigned using the semi-empirical AM1-BCC^49^ procedures in antechamber^50^. The missing ligand parameters were obtained using the parmchk2 tools available within the AmberTools^51^. Subsequently, the ligand-M^pro^ complexes were prepared in using tleap program in AMBER18 package^52^ by solvating the complex in a cubic box of TIP3P water molecules with a minimum of 12 Å distance between the box boundaries and the closest solute atoms. The solvated systems where subsequently charge neutralized with 150 mM concentration of NaCl counter ions. Thus, the systems were prepared for subsequent MD simulation.

### 2.2. MD simulation of ligand-M^pro^ complexes

All MD simulations were performed using the AMBER18 molecular dynamics program^52^ with pmemd.cuda engine. MD simulation of each ligand-M^pro^ complex was performed in five consecutive stages: (1) energy minimization, (2) NVT heating, (3) NPT equilibration, (4) NPT pre-production simulation, and (5) production simulation. The initial stage of gradient minimization of the system was performed in six rounds. In the first round of minimization, a strong harmonic constraint of 100 kcal mol^−1^Å^−2^ was applied on the solute atoms and 10000 steps of minimization (1000 steps of steepest descent minimization + 9000 steps of conjugate gradient minimization) was performed. The next four rounds of minimization was performed with almost same parameters as the first round except for the harmonic constraints that was reduced as 50>10>5 kcal mol^−1^Å^−2^ in each round. Finally, 20000 steps of energy minimization of the system without any constraints was performed. Following the minimization, each complex was gradually heated to 310K over a duration of 100 ps and, subsequently, equilibrated in four rounds, each being 400 ps long. In the first two rounds of equilibration, positional restraints were applied on the nonhydrogen atoms of the protein residues using restraint force constants of 5 kcal mol^−1^Å^−2^ (first round) and 0.1 kcal mol^−1^Å^−2^ (second round). Following this, the third round of equilibration was performed with a weak restraint of 0.01 kcal mol^−1^Å^−2^ applied to only the backbone atoms of protein residues. Finally, a 400 ps long restraint-free equilibration of the system were carried before proceeding to production simulation. The equilibrated system underwent a 2 ns long preproduction simulation under isothermal-isobaric (NPT) conditions. Following the pre-production run, each complex was subjected to 3 subsequent 10 ns long MD simulation (making a total of 30 ns long MD simulation) using an integration timestep of 2fs. Throughout the simulation, temperature was controlled using the Langevin thermostat^53^ and the pressure was kept at 1 bar using the Berendsen barostat^54^. During the production runs, coordinates were stored every 2 ps thereby resulting in a total of 15000 snapshots from a 30ns long MD trajectory. VMD 1.9.3^55^ and USCF Chimera^56^ were used to visualize the molecular dynamics trajectories. All simulations were performed using the Graham and Cedar GPU clusters available within the ComputeCanada infrastructure. The resulting MD trajectories of the ligand-M^pro^ complexes were processed through the CPPTRAJ tool ^57^ to generate the root mean square deviation (RMSD) plots and the per-residue root mean square fluctuation or RMSF (for the apo trajectories only).

### 2.3. Relative binding free energies of ligand-M^pro^ complexes

Following the MD simulation, for each complex, the last 10ns of the MD trajectory was used to compute the binding free energy scores between the ligand and the M^pro^ bound in a complex. For this purpose, we employed the MM-GBSA method^58^ with the implicit solvent model of GB-Neck2 (or igb=8)^59^. The snapshots sampled at a regular interval of 10 ps and thus a total of 1000 frames were used to calculate the MM-GBSA energies. The binding free energy (ΔG_bind_) using the MM-PB(GB)SA can be estimated as,

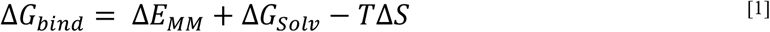

Here, ΔE_MM_ is the summation of non-bonded and bonded interaction energies^58^. The solvation energy, Δ*G_solv_*, is the sum of the polar and non-polar contributions of solvation, where the polar solvation terms are calculated using a Generalized-Born model or a Poisson-Boltzmann solver and the non-polar terms are computed based on the size of the solvent accessible surface area in the ligand and M^pro [58]^. However, the last term corresponding to conformational entropy (TΔS) is neglected as it is computationally expensive. It should also be noted that previous studies^60,61^ have shown that incorporating the entropic contribution in the ΔG_bind_ calculations do not assure accurate binding free energy. Therefore, in this work, we calculate only the relative binding free energy, which is often considered as useful in relative ranking of compounds. All MM-GBSA calculations in this study were carried out using the MMPBSA.py script^62^ included in the AmberTools^52^. The pairwise decomposition analyses (idecomp=4) using the MD trajectories were also carried out so as to identify the key ligand-residue energetic contributors to the binding free energy of the complexes^63^.

### 2.4. Relative binding free energies of ligand-M^pro^ complexes in the presence of explicit water molecules

The binding free energies of ligand-M^pro^ complexes were also calculated in the presence of 0-6 explicit water molecules. These calculations were performed using a MM-GBSA variant called the NWAT-MMGBSA. Maffucci et al^64^ demonstrated that this NWAT-MM-GBSA method is able to improve the correlation between the calculated and measured binding free energies of biological systems. All parameters in the NWAT-MM-GBSA calculations were same as those explained for the routine MM-GBSA method (section 2.3), except the explicit water molecules that were captured from the MD trajectories using the CPPTRAJ tool^57^ of Amber 18 program^52^. The cpptraj module was employed to select the desired number of snapshots and to generate the related topology files for the free energy calculations; whereas, the interface residues were found using the “InterfaceResidues” Pymol script. The NWAT calculation for each system was repeated by increasing the number of explicit water molecules from 0 to 6. All NWAT calculations utilized the shell scripts from Maffucci et al.^64,65^.

### 2.5. Preparation of mutated systems

Each of the selected ligand-Mpro complexes were modified by a single point mutation of HIS163 to ALA163 using the ‘swapaa’ command in UCSF Chimera^56^. The mutated systems were then prepared and subjected to 30 ns long MD simulation as discussed in the above sections for the wild type complexes (in sections 2.1-2.2). However, for the mutated systems, the relative binding free energies were calculated only using the MM-GBSA approach (section 2.3) and the free energies in the presence of explicit water molecules were not computed.

### 2.6. Preparation of ligand-bound dimer models of Mpro

The models of ligand-bound Mpro dimer complexes were constructed using the experimentally resolved 3D structure of a Mpro dimer in the PDB (PDB: 6WTT). This was performed by aligning the ligand-bound Mpro monomer on the structure of Mpro dimer and the duplicate chain after alignment was deleted to obtain the ligand-bound dimer complex. For example, we modeled the dimer structure using the PDB structure of 5RFA, which represents a ligand-bound Mpro monomer complex. This 5RFA structure was aligned against the chain A of the 6WTT structure and it is expected that chains will be 100% aligned as they are essentially the structures from the same sequence. After alignment, the bound-ligand from 5RFA was extracted into the dimer structure (6WTT) and the protein chain of 5RFA was deleted. This resulted in a starting configuration of a ligand-bound Mpro dimer complex. The same process was carried out to build all the 13 dimer complexes in this study. All produces in building the dimer models were performed using the UCSF Chimera. Subsequently, all the dimer complexes were subjected to 30 ns long MD simulation and followed by binding free energy estimation using MM-GBSA method as described above.

## 3. Results and Discussion

### 3.1. MD simulation and analyses of apo SARS-CoV-2 M^pro^

In order to understand the dynamics of a SARS-CoV-2 M^pro^ in the absence of a bound ligand, we performed 30 ns long MD simulation of an apo structure of M^pro^ downloaded from PDB (PDB ID 6M2Q). Initially, the stability of the structure was assessed by plotting the fluctuation of backbone RMSD (Fig. 3a) during the course of MD simulation. It was noted that the backbone RMSD of M^pro^, as a whole, exhibited a fluctuation in the range of 1-3 Å, most often only varying within 0.5 – 1 Å. As can be seen in Fig. 3a, the RMSD plot showed a slightly increased fluctuation, reaching ~3 Å, around 14-15 ns timescale that was caused due to the movements of flexible C- and N-terminal loops. We analyzed the backbone RMSDs of the individual domains such as domain I-III in M^pro^(see in Supplementary Figure, S.Fig. 1) so as to deduce the dynamic segments. We found that domains I and II remained quite stable throughout the simulation with backbone RMSD changes within ~0.7 and 1.2 Å. Nevertheless, Domain III that features a globular cluster of five helices exhibited fairly higher fluctuations that reached >1.5 Å during the mid-course of simulation and reaching a plateau during the last 10 ns. We performed RMSF analyses (Fig. 3b) to reveal the averaged per-residue fluctuations during MD simulations. The most stable segments of M^pro^ include the β-sheets in domains I and II, the loop connecting these domains (ASP92-PHE112), and the helices of domain III. Unsurprisingly, the most flexible regions occurred at the N and C terminus of the protein ^45^. In addition to this segment, the other regions that displayed elasticity during MD were the loops connecting different secondary structures in domain III (ASN277-THR292), domain II (ASP153-ASP155 and LEU167-VAL171), and domain I (THR45-ASN65 and ALA70-VAL73). The loops enclosing the catalytic site in M^pro^ such as CYS44-PRO52 loop and the PHE185-THR201 linker loop (domain II-III linker) only exhibited small variations indicating their role in stabilising the active site. The catalytic residues, HIS41(from domain I) and CYS145 (from domain II) remained highly stable throughout the simulation. Despite the small flexibility within the different loop regions, the apo protein structure proved to be quite stable and only reached a maximum RMSD of 3 Å and a maximum RMSF of 2 Å. We also assessed different electrostatic interactions in M^pro^ that contributed to the stability of the structure during MD (Supplementary figure S.Fig. 2). We found strong hydrogen bonds (H-bonds) between the carboxylate side chain in THR111 and the hydroxyl moiety in ASP295 in domain II of M^pro^ (plot shown in supplementary figure, S.Fig. 3a). In addition, we found stable salt bridge interactions formed by two ASP-ARG residue pairs: one pair of salt bridge was formed between ARG131 in domain II and ASP289 in domain III (supplementary figure, S.Fig.3b); and another pair of salt bridge was established between ARG40 in domain I and ASP187 located in the DII-DIII linker loop (supplementary figure, S.Fig.3c). These observations are consistent with the previous studies^66,67^. In addition, we also identified a critical water-mediated interaction near the active site of Mpro, where a central water-molecule formed a 3-way H-bond network with residues HIS41 (the catalytic residue), ASP187 and HIS164. We noticed that whenever the water molecule at this tri-junction moved out of the binding site during MD, another water molecule entered the site and maintained this H-bond network. This suggests the critical role of water molecules in the stability of the binding site during MD simulation. Moreover, as indicated earlier, it is hypothesized that this water molecule in the binding site could play a role of the third residue in regulating catalysis process in M^pro[25,29]^.

**Fig. 3.**
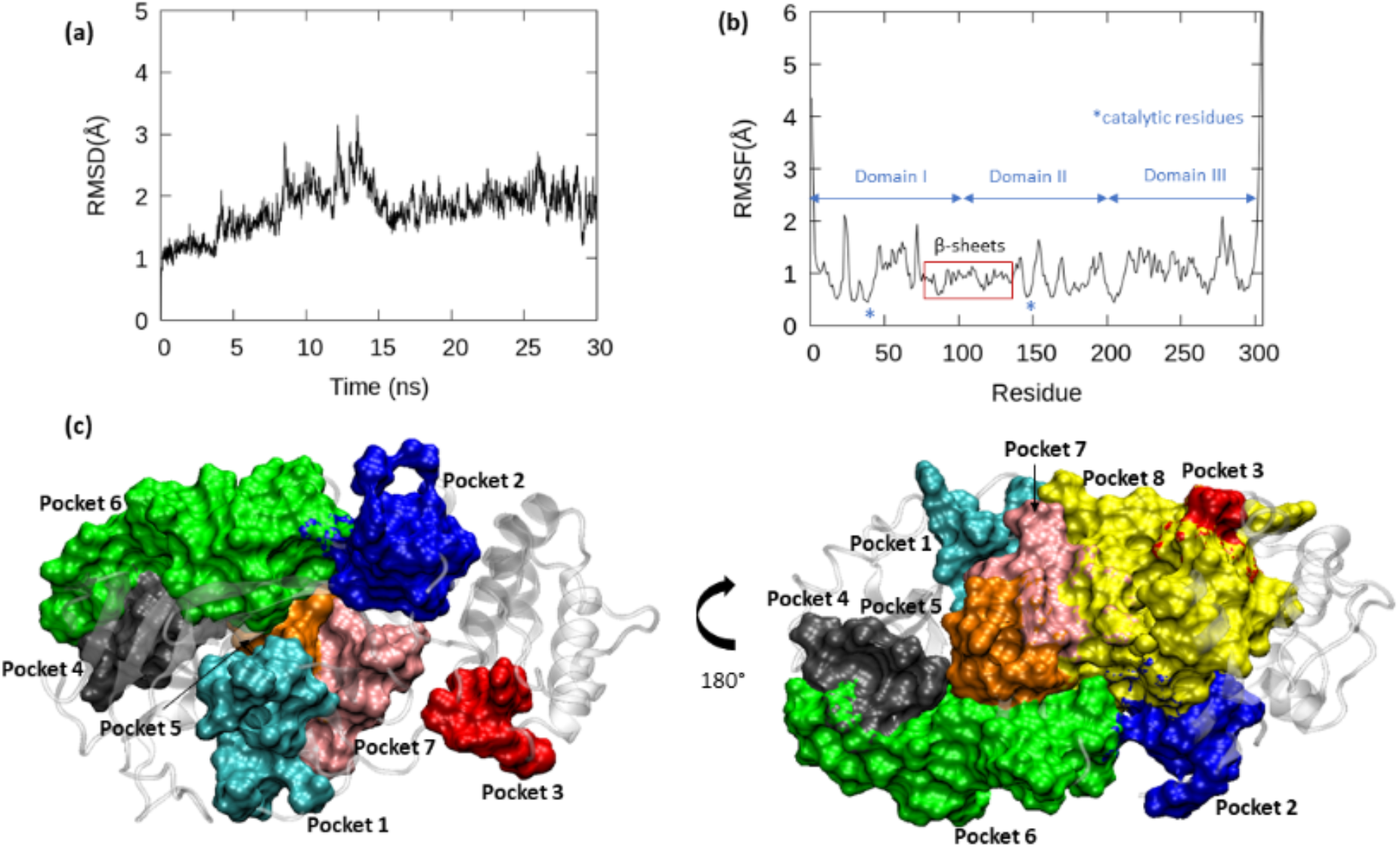
The backbone RMSD (a) and RMSF (b) fluctuations of apo Mpro (PDB: 6M2Q) and the different pockets on the M^pro^ (c and d) identified through MD simulation. RMSD of the apo structure changed between 1 – 3 Å during MD and the per-residue fluctuation during MD is shown as RMSF and the different domains in the enzyme are marked. Using the “MDpocket” tool, various transient pockets were identified and characterized within the apo structure during the MD simulation.

Since, the 62 crystal structure complexes studied in this work show ligand binding at different sites other than the actual orthosteric catalytic site in M^pro^, we explored the dynamics of different pockets in M^pro^. A protein pocket detection tool, MDpocket^68^, was employed for this purpose. Using the 30 ns MD trajectory of the apo Mpro, we sampled snapshots at a regular interval of 10 ps from the last 25 ns, thereby, using 2500 snapshots of the enzyme for the pocket analysis. Each snapshot was saved as an individual PDB file and was used to determine favourable binding pockets within the protein. In addition to the main binding site, many small and transient pockets were identified using the putative channels and small cavities search options within MDpocket^68^. Through this analysis, we identified a total of 8 pockets in Mpro monomer (Fig. 3c-d) that we name as Pocket1-Pocket8 in the text hereafter. It should be noted that ligands binding in all these pockets have been described in the PDB structures studied here. A table comparing different characteristics of each of these pockets such as residues, pocket volume, and PDB codes of the structures with ligand bound in the respective pockets are presented in the supplementary table ST. 2. The parameters used for MDpocket analyses are also provided in the footnote of this supplementary table. Pocket 1 is obviously the catalytic pocket that is encircled by the loop composed of PHE140-CYS145, a β-sheet (‘L’ in Fig. 1c) made of HIS163-GLU166, and the residues from the domain II-III linker loop including HIS172, VAL186, ASP187, ARG188, GLN189, and GLN192. The volume of this pocket was ~800 Å^3^ in our MD snapshots and, as expected, most compounds in this study are bound in this pocket. Among the other pockets, three pockets (including pockets 2, 3, 7) are found within the domains II and III, one pocket (pocket 4) is found in domain I, and two pockets (pocket 5, 6) are found between domains I and II (Fig. 3cd). Pocket 2 is a small cavity (mostly volume < 320 Å^3^) that is found between domains II and III and formed by the N-terminal (PHE3-PHE8) and C-terminal(PHE291-THR304) residues, along with residues THR111 and GLN127 from domain II. In two of the crystal structures studied here, 5RFA (1-methyl-N-{[(2S)-oxolan-2-yl]methyl}-1H-pyrazole-3-carboxamide) and 5RGQ (1-(4-fluoro-2-methylphenyl)methanesulfonamide), ligands were found to bind at this pocket. Pocket 3 is another small and transient surface pocket formed by THR198 (from domain II-III linker) and the selected residues from domain III (MET235 ASN238 TYR239 GLU240 PRO24). Fragments such as 2-{[(1H-benzimidazol-2-yl)amino]methyl}phenol (PDB: 5REC), [(2~{R})-4-(phenylmethyl)morpholin-2-yl]methanol (PDBL: 5RGS), and (2R,3R)-1-benzyl-2-methylpiperidin-3-ol (PDB: 5REE) were found to bind in this pocket. Similar to pocket 2, pocket 7 is another transient pocket that is mainly formed by residues from domain II (ARG105-PRO108, MET130, PHE134, PHE181-PRO184) but is located near the interface between domains II and III. It may be due to the transient nature of this pocket that only one carboxamide containing ligand (PDB: 5REG) was found to bind in this pocket. Pocket 4, a fairly large pocket of 600 Å^3^, was found to be formed majorly by the β-sheets composed of residues ASP34-TYR37, GLY79, HIS80, SER81, LYS88-LYS90. This pocket remained stable throughout out MD simulation, which suggests that this could be a viable pocket for ligand binding. In consistent with this, we found that ligands (of different size and class) from at least 7 known crystal structures (5RFC, 5RH4, 5RE6, 5RE5, 5RGG, 5RFB, and 6YVF) were found to be bound in this pocket. Pocket 5 is slightly smaller than Pocket 4 but is also quite stable throughout the MD trajectory. This pocket is positioned near the catalytic site (or Pocket 1) between domains I and II, but on the opposite face of the protein. Four ligands were found to interact with this pocket in the crystal structures (PDBs: 5REI, 5RED, 5RF5, 5RGR). Nevertheless, there is another large pocket of 800 Å^3^, Pocket 6, that is also found between domains I and II and five ligands from the PDB structures of 5RF4, 5RFD, 5RE8, 5RF9, and 5RFJ were found to be interacting at this site. This suggests the stable nature of this pocket under experimental conditions, much like the behaviour observed during our MD simulation. In addition to the well-defined pockets discussed above, we found a huge surface area (dubbed here as pocket 8) of 2,250 Å^3^ that was found largely on the surface of domains 2 and 3 (Fig.3c) and connecting many of the smaller cavities. There is only one crystal structure (PDB: 5RF0) that describes the binding of a ligand within this surface. Mapping these residues onto a dimer structure of Mpro (6WTT), we found that this surface is mostly buried at the dimer interface (refer to the figure provided in the foot note of supplementary table, ST. 2), which explains the existence of this dynamic surface area during MD analyses. However, since the dimerization in SARS-CoV-2 M^pro^ is crucial for the activity of the enzyme and thus the replication of the virus, this large surface area of pocket 8 can be an interesting site for drug-binding, as targeting dimerization in M^pro23^ is considered as a potential therapeutic strategy to inhibit the virus replication.

### 3.2. Stability of ligand-SARS-CoV-2 M^pro^ complexes during MD simulation

Following the analyses of pockets and their dynamics in the apo structure of M^pro^, we performed MD simulation and analyses of the selected crystal structures (supplementary table, ST. 1) in this study. As described in the methods section, we performed 30 ns long MD simulation of each of the 62 reversible ligand-M^pro^ complexes that resulted in a total of 1.86 μs long molecular trajectories for analyses. We analyzed the stability of the protein and ligand structures using the RMSD analyses. In order to simplify the data presentation, we calculated the average backbone RMSD fluctuations of the proteins and the average RMSD fluctuation of ligand in each complex during the course of MD simulation, which are provided in the supplementary table, ST. 3. As can be seen in Fig. 4a, the average backbone RMSD variation of proteins in all the complexes were mostly in the range of 1.5-2.3 Å, when compared to that of 1.81 Å for the apo (or ligand-free) Mpro structure. This suggests that the binding of ligand had only limited impact on the structural dynamics of the enzyme. Two PDB structures, 5RE5 and 5REG, were the only exception to this trend and displayed slightly higher average backbone RMSD value of 2.84 Å and 2.87 Å, respectively, which are still within the fluctuations that are usually seen for proteins of this size, as also previously shown by a MD study^66^ of SARS-CoV-2 Mpro structure. Analyses of their trajectories showed that the PHE185-THR201 linker loop was slightly more flexible in these complexes during the course of simulation. The flexible nature of this long loop connecting domains II and III is already known^66,69^.

**Fig. 4.**
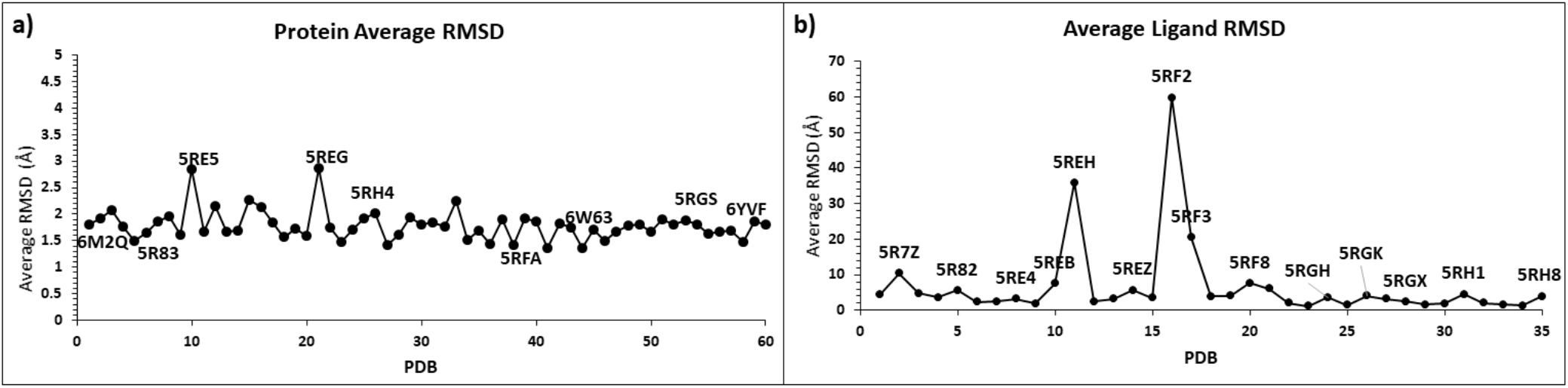
Average RMSD for the proteins in all the 62 systems studied here with reference to that of the apo structure of SARS-CoV-2 M^pro^ (a), and average unaligned RMSD of ligands bound with the catalytic site in M^pro^ (b). (a) Protein backbone RMSD remain stable throughout MD with exception of 2 peaks for PDB ID 5RE5 and 5REG. (b) Except for a few unstable ligands, most of the other ligands remained stable in the binding site during the course of MD.

Nevertheless, unlike the stable proteins, ligands that were co-crystalized in the structures studied here displayed a spectrum of behaviour that described their nature of stability and interactions with the target. We initially analyzed the stability of the ligands bound within the HIS41-CYS145 catalytic binding site (or pocket 1), refer to Fig. 4b. It was noted that most of the ligands bound within the catalytic binding site exhibited an average RMSD in the range of 1-6 Å. Lower the RMSD value (i.e., < 3 Å) means the ligand was more stable in the initial pose (e.g., ligands in PDB structures: 5RG1, 5RGW, 6W63, 5RH3); whereas, an average RMSD in the range of 3-6 Å indicate that the ligand, despite bound stable in the catalytic site, underwent some conformational changes to stabilize the association with M^pro^ (e.g., ligands in the PDB structures 5R7Y, 5R82, 5REZ, 5RH8, and 5RHD). For instance, in the 5RHD structure (in supplementary figure, SFig.4), the 1-[4-(methylsulfonyl)phenyl]piperazine ligand was initially bound in a pose in which the methylsulfonyl group was interacting with the “L” β-sheet residues such as MET165-LEU167 and the piperazine moiety is stabilized by the H-bond interactions with CYS44-PRO52 loop that flanks the catalytic site. Nevertheless, during the course of MD simulation (in the first 10 ns of simulation), the methylsulfonyl group changed its orientation and interacted with the loop containing GLY143, SER144 and CYS145. Thus, this change of binding pose of the ligand during MD simulation resulted in a slightly higher RMSD range of over 4.5 Å. However, there were a few ligands that bound within the catalytic site and had weak association with M^pro^ that resulted in their unbinding. For instance, the smallest ligands in this study such as 1-azanylpropylideneazanium (in 5RF2) and pyrimidin-5-amine (in 5RF3) that were originally bound within the binding site displayed weak interactions with M^pro^ and egressed during the course of MD simulation. This explains the large average RMSDs for these ligands (in Fig. 4b) during our MD simulation. Another ligand (~{N}[2-(5-fluoranyl-1~{H}-indol-3-yl)ethyl]ethanamide) that was co-crystallized in PDB 5R7Z was initially bound in a pose in which the fluroindole group was buried into the catalytic site and the ethanamide group was projecting towards the surface of M^pro^. However, during the course of 30ns long MD simulation, this ligand explored different orientations within the catalytic site and eventually unbound from the pocket. Similar behaviour was also seen in 1-cyclohexyl-3-(2-pyridin-4-ylethyl)urea bound with M^pro^ in 5REH structure, where the pyridine ring was bound inside the pocket and the cyclohexane moiety stuck outside on the surface that resulted in weak interactions with M^pro^ and egression during MD. It should be noted that the current simulation of ligand-bound complexes are based on the monomer state of Mpro. However, it is known that Mpro exists as a dimer, where the N-terminus of the second chain will close the binding site of the first monomer chain. Therefore, it is possible that 5R7Z could be a more stable ligand under a dimer conformation. In order to address this concern, we analysed the dynamic interactions of the select ligands with the dimer state of M^pro^. These include the ligands that were originally bound within the catalytic site of Mpro and unbound during MD. The results of these dimer simulation are discussed in the later section.

Next, we analyzed the stability of complexes in which the ligands were bound in the other pockets (pockets 2-8) found in the apo structure. The average RMSD of the ligands bound in these pockets are presented in the supplementary figure, SFig.5. Two ligands, 1-(4-fluoro-2-methylphenyl)methanesulfonamide (5RGQ) and 1-methyl-N-{[(2S)-oxolan-2-yl]methyl}-1H-pyrazole-3-carboxamide (5RFA) were bound in pocket 2 composed of the N-terminal and C-terminal residues of Mpro monomer, which forms the dimer interaction surface. While the former ligand remained stable throughout 30 ns long MD (average RMSD of 2.29 Å), the latter disassociated during the course of simulation and displayed higher RMSD changes. Despite pocket 3 between domains II and III is a small cavity, the ligands 2-{[(1H-benzimidazol-2-yl)amino]methyl}phenol (5REC) and [(2~{R})-4-(phenylmethyl)morpholin-2-yl]methanol (5RGS) remained highly stable highlighting their complementing properties to this pocket through H-bond interactions with TYR239 and GLU240 in this pocket (more discussion are presented in the following section). Nevertheless, the piperidinol based ligand (in 5REE) was not able to make such key H-bonds therefore unable to maintain stable interactions with pocket 3, which is reflected by its average RMSD of 12.47 Å during MD. From the RMSD analyses, it is clear that the pocket 4 favours an acid group or its derivatives when compared to an amide or amine-based ligands. For example, the ligands based on benzoinc acid (6YVF), propanoic acid (5RH4), and carbamate (5RFC) exhibited stable interactions with this pocket. Nevertheless, the carboxamide-based (5RGG and 5RE5), acetamide-based (5Re6) and ethanamine (5RFB) ligands remained loosely bound to this pocket. It was interesting to note that all the ligands that bound to pockets 5 and 6 that are located between domains I and II displayed weak affinity towards this site, as all of them unbound during the MD simulations. This suggests that this site is either not druggable or the ligands explored in this screening do not have physicochemical features to complement this site. Thus, our stability analyses highlight the stability and dynamic interactions of the experimentally reported ligand-M^pro^ complexes.

### 3.3. Binding free energy analyses of ligand-M^pro^ complexes

In the earlier section, we discussed the stability of reversible ligands in different pockets in SARS-CoV-2 M^pro^ structure. Building on these analyses, we calculated the binding free energy of the ligand-Mpro complexes using the end-point MM-GBSA method based on the snapshots of the complexes sampled from their respective MD trajectories. For each complex, we sampled the snapshots at a regular interval of 10 ps from the last 10 ns trajectory, which resulted in 1000 frames for the MM-GBSA calculations. The binding free energies of the complexes are provided in the supplementary table, ST. 3 We also decomposed the ligand-residue pairwise energetic contribution to the binding free energy of the complex. Our intention is to evaluate the binding affinity of the ligand-M^pro^ complexes that were relaxed through MD simulations and identify the key residues contributing to their affinity. As expected, the complexes in which the bound-ligand disassociated in the course of MD (as discussed in the earlier section) displayed a low affinity of > −10 kcal/mol that describes weak and non-specific interactions with the surface of M^pro^ during the simulation (supplementary figure, SFig. 6). This is particularly true for the ligands bound in pockets 5 and 6 located at the junction of domain I and II, as these complexes did not show any stable interactions. In the similar nature, those ligands that egressed from the catalytic pocket also had weak affinity against Mpro and these include the structural complexes of 5REH, 5RF2, 5RF3, 5RF8 and 5R7Z as discussed in the stability analyses. Fig. 5 compares the binding free energies of ligands bound to pocket 3 (Fig. 5a) and pocket 4 (Fig. 5b). In pocket 3, the ligands, 2-{[(1H-benzimidazol-2-yl)amino]methyl}phenol (5REC) and [(2~{R})-4-(phenylmethyl)morpholin-2-yl]methanol (5RGS), displayed a binding affinity scores of −17.56 kcal/mol and −14.79 kcal/mol, respectively. Analysing their binding poses (in Fig. 5a), the alcohol (-OH) functional group in these compounds bound inside the cavity and was stabilized by the hydrogen bond interactions with TYR239, GLU240 and MET235. Nevertheless, in the binding pose of the ligand ((2R,3R)-1-benzyl-2-methylpiperidin-3-ol) that was bound to pocket 3 in Mpro (5REE complex), the phenyl ring was occupying the cavity and the OH group in the piperidin-3-ol moiety was not able to make H-bond interactions with the key residues in this pocket. Consequently, the ligand moved out of the pocket 3 during MD, thereby resulting in a poor affinity of ~-2 kcal/mol (in Fig. 5a). Earlier we noted that pocket 4 favoured the binding of an acid group as opposed to the amide or amine-based fragments. In consistent with this interpretation, the binding affinity scores of propanoic acid ligand (5RH4) and benzoinc acid ligand (6YVF) were −17.50 kcal/mol, −15.35 kcal/mol, respectively (Fig. 5b). Analyses of their binding poses show that acid groups in these ligands bury themselves in the cavity. In this orientation, they are able to make electrostatic interactions (H-bonds) with the LYS residues in the pocket (LYS88 and LYS90). Energy decomposition analyses revealed that the pairwise interactions of the ligand with key restudies such as HIS80, SER81, LYS88 and LYS90 make significant contributions to the binding affinity of the complexes (Fig. 5b). Nevertheless, the carbamate-based ligand in the 5RFC structure (Fig. 5b), a hydrophobic ring deep inside the pocket but has a carboxy group that was placed on the edge of the pocket thereby allowing it to make some electrostatic contacts with SER81 and LYS90. This pose was not favoured as a result, this ligand displayed weaker affinity (−10.63 kcal/mol) when compared to the other acid-based ligands. Nevertheless, the amide or amine compounds including carboxamide-based (5RGG and 5RE5), acetamide-based (5Re6) and ethanamine (5RFB) ligands were not able to accommodate the cavity in pocket 4, as a result they were not stable in the pocket and made several transient interactions. The binding free energy scores in Fig.5b does not represent the ligand affinity to one site rather a cumulative effect of interacting with different sites on the surface of M^pro^ during the course of MD. Fig. 6a compares the binding free energies of the complexes in which the ligands were bound within the catalytic site of Mpro throughout the simulation. From our analyses of MD trajectories of these complexes, it was clear that the ligand bound in the active site (or pocket 1) was mainly stabilized by their interactions with four segments, apart from the catalytic dyad residues, HIS41 (from domain I) and CYS145 (from domain II). These include (1) a loop (dubbed as ‘L1’) connecting the J and K β-sheets leading to CYS145, shown as yellow in Fig. 6b, (2) domain II-III linker (shown as green in Fig. 6b) composed of residues VAL186 through ALA193 and flanking on the other side of the active site (dubbed as ‘L2), (iii) The ‘L’ β-sheet (see in Fig. 1c) that is lining the active site and composed of residues HIS163-LEU167 (shown as blue in Fig. 6b), and (iv) key hydrophobic residues surrounding the catalytic HIS41 such as LEU27 and MET49. Depending upon how strongly the ligands engaged with these segments, their binding affinity scores varied. For example, the ligand, 5-fluoro-1-[(5-methyl-1,3,4-thiadiazol-2-yl)methyl]-1,2,3,6-tetrahydropyridine, was bound to the catalytic site of M^pro^ (PDB: 5RGH) and had a weak binding affinity of −12.10 kcal/mol. Analyzing their trajectory, we found that the fluorine-attached tetrahydropyridine ring was anchored to the ‘L1’ loop through the interaction of fluorine atom with that of GLN189 and GLN192 residues (supplementary Fig, SFig. 7a). However, the other methyl-attached thiadiazol moiety of the ligand remained flexible and underwent significant conformational changes during MD (supplementary Fig, SFig. 7b). As a result of this fluctuation in the ligand pose, the affinity of this complex remained low. In a similar nature, the ligands cocrystallized with M^pro^ in complexes 5R82 (6-(ethylamino)pyridine-3-carbonitrile), 5R80 (methyl 4-sulfamoylbenzoate), 5RHD (1-[4-(methylsulfonyl)phenyl]piperazine), 5RGK (2-fluoro-N-[2-(pyridin-4-yl)ethyl]benzamide), and 5REB (1-[(thiophen-3-yl)methyl]piperidin-4-ol) were also anchored to one of the four segments (in Fig. 6b) and remained dynamic within the catalytic binding site. Therefore, these complexes had a binding affinity > −15 kcal/mol as seen in Fig. 6a. For example, the conformational dynamics of the ligand, 6-(ethylamino)pyridine-3-carbonitrile, in the 5R82 structure is clearly described in the supplementary figure, SFig. 8.

**Fig. 5.**
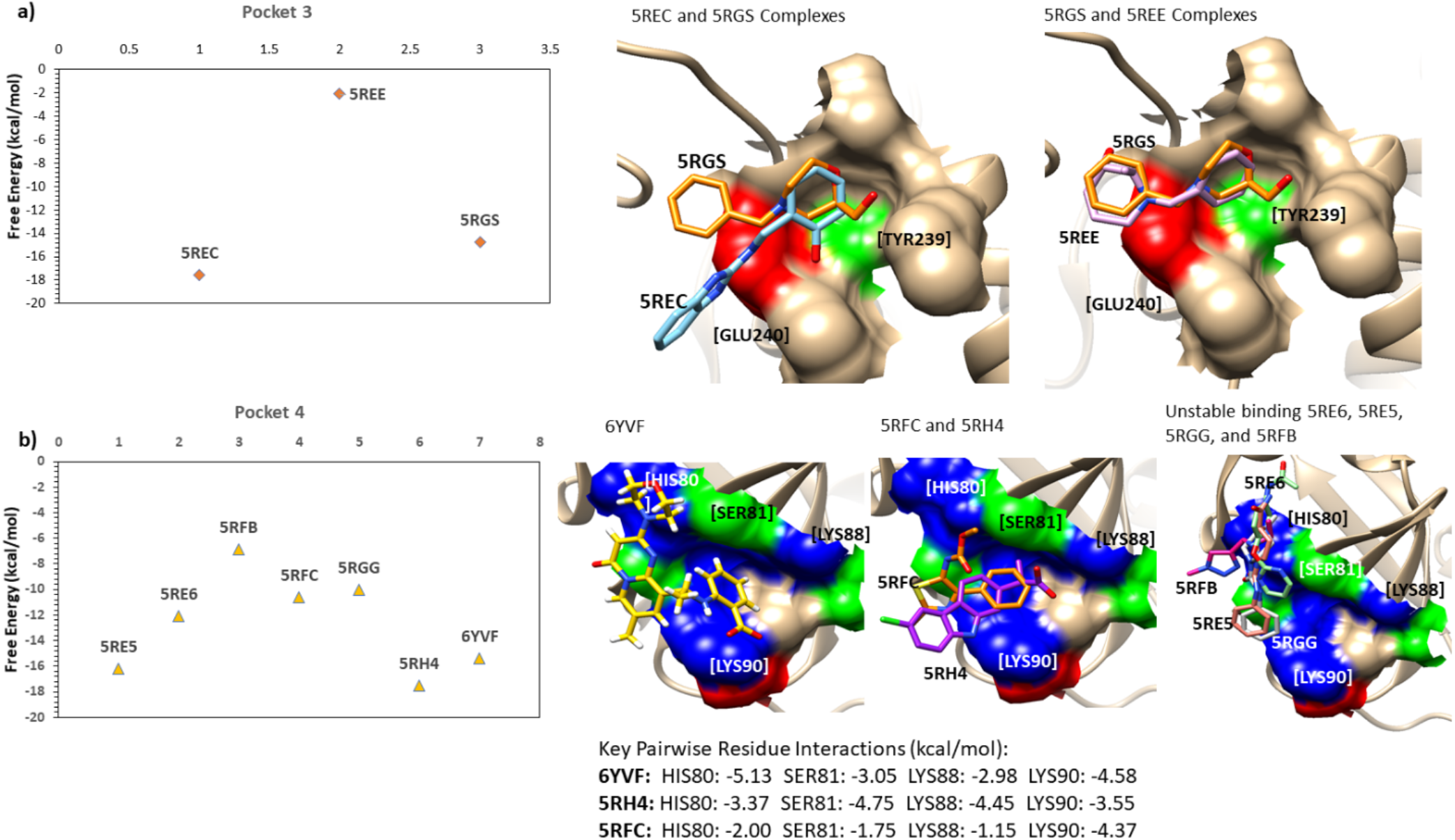
The binding free energy of the ligand-M^pro^ complexes in which the ligands were bound in pocket 3 (a) and in pocket 4 (b) along with their respective binding poses shown. Both in pocket 3 and pocket 4, a few ligands remained stable due to their ability make key electrostatic interactions with the residues in the pocket while the other ligands failed to make such interactions that resulted in their weak affinity against M^pro^.

**Fig. 6.**
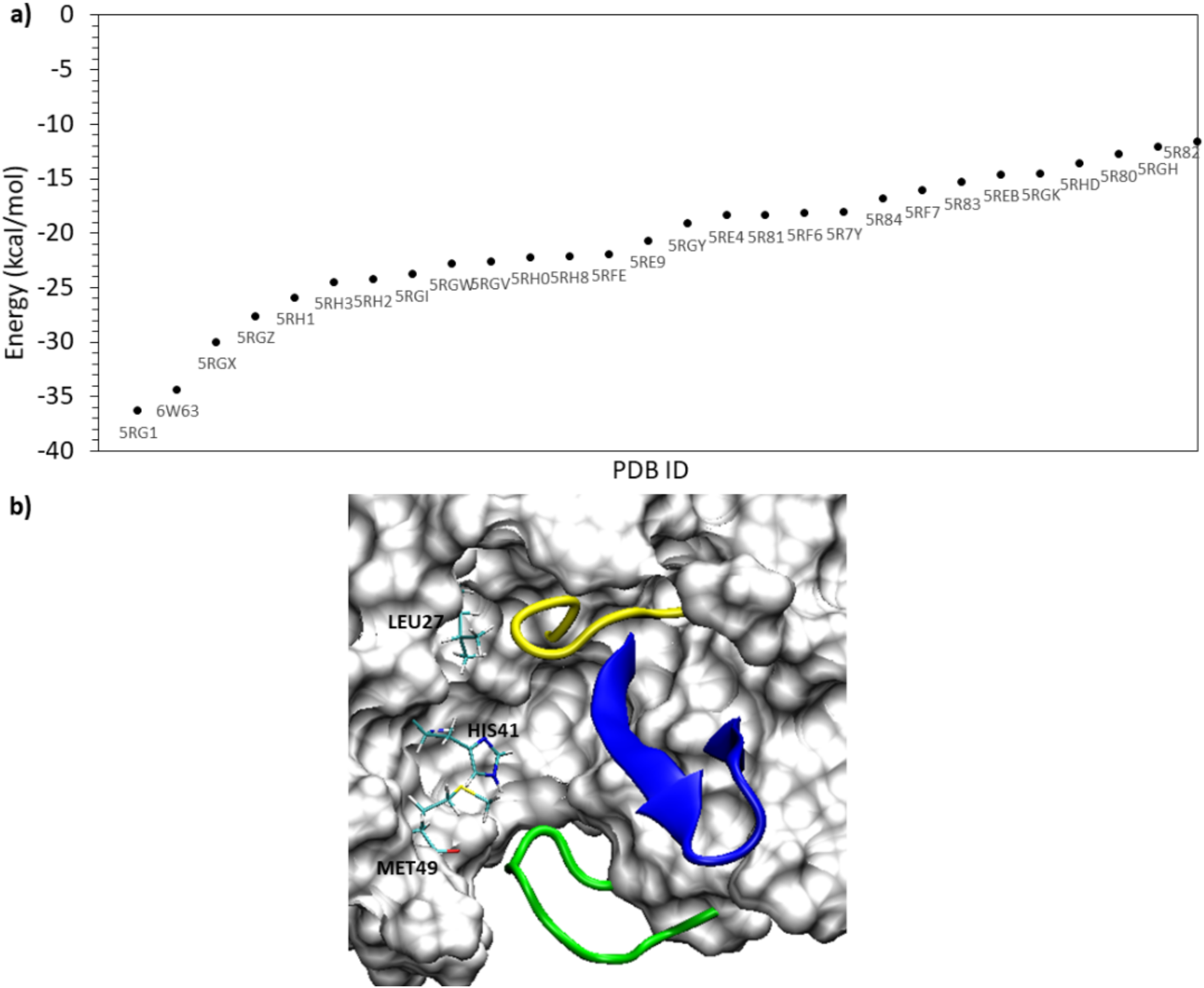
The scatter plot comparing the binding free energies of the ligand-M^pro^ complexes in which the ligands were bound in catalytic site (a) and the identification of four key components that are critical for ligand binding in this site (b). In (b), two loops flank the active site pocket: the yellow loop made up of residues in the 140s range (called the L1 loop); and the green loop that is part of the domain II-III linker Loop (called the L2 loop). In addition, the ‘L’ β-sheet is shown in blue, and the residues such as HIS41, LEU27, and MET49 located deep inside the pocket are shown as stick representation.

We noted that the ligands bound to the catalytic site and stabilized by the interactions with two of the four segments described above exhibited a better binding affinity score that is < −15 kcal/mol (in Fig. 6b). For example, the bulky compound in the series studied here is N-(4-tert-butylphenyl)-N-[(1R)-2-(cyclohexylamino)-2-oxo-1-(pyridin-3-yl)ethyl]-1H-imidazole-4-carboxamide that was co-crystallized with M^pro^ in the pdb structure 6W63. This compound was bound against M^pro^ in a pose (in Fig. 7), in which the pyridine ring occupied a lateral cavity formed by the ‘L1’ loop (yellow) and the ‘L’ β-sheet and formed a stable H-bond with HIS163 residue (Fig. 7c) in this cavity. In addition, the OH groups in this ligand made strong H-bonds with GLU166 and GLY143 (refer to the 2D interaction diagram shown in Fig. 7d). In addition, the imidazole ring interacted with HIS 41, and the methyl group attached to cyclohexamine moiety made hydrophobic contacts with the ‘L2’ loop (i.e., the domain II-III linker loop). Pairwise decomposition of interaction energies identified the key residues stabilizing this complex, which include HIS41, ASN142, GLY143, MET165, and GLU166. The interaction of ligands with these key residues contributed at least −3 kcal/mol energy to the binding free energy. Therefore, given the complementary properties between the ligand and the protein in the 6W63 structure, this complex had a high binding affinity score of −34.4 kcal/mol. Unlike the compound in 6w63 complex, the Nalpha-acetyl-N-(3-bromoprop-2-yn-1-yl)-L-tyrosinamide ligand in the PDB structure of 5RG1 is a much linear compound; yet, it displayed strong binding affinity of −36.30 kcal/mol towards M^pro^. However, similar to the former, the phenol group in this ligand also filled the lateral pocket (Fig. 8a) through H-bonds with the imidazole side-chain of HIS163 residue (Fig. 8c). In addition, the ligand also formed two strong H-bonds with GLU166 (in Fig. 8c), and an addition H-bond with GLN189. Decomposition analyses identified HIS163, MET165, GLU166, ARG188, GLN189 as key energetic contributions to the binding free energy (Fig. 8b). Similar to these compounds, several other pyridine-containing compounds exhibited similar interactions and possesssed a binding affinity value better than −22 kcal/mol (Fig. 6a). These include (2R)-2-(3-chlorophenyl)-N-(4-methylpyridin-3-yl)propenamide (5RH3), 2-(3-cyanophenyl)-N-(4-methylpyridin-3-yl)acetamide (5RGX), 2-(3-cyanophenyl)-N-(pyridin-3-yl)acetamide (5RGZ), 2-(5-chlorothiophen-2-yl)-N-(pyridin-3-yl)acetamide (5RH1), (2R)-2-(3-chlorophenyl)-N-(4-methylpyridin-3-yl)propenamide (5RH3), 2-(3-chlorophenyl)-N-(4-methylpyridin-3-yl)acetamide (5RH2), 2-(5-cyanopyridin-3-yl)-N-(pyridin-3-yl)acetamide (5RGW), and N-(5-methylthiophen-2-yl)-N’-pyridin-3-ylurea (5RH0). In all these compounds, the pyridine ring occupied the lateral pocket and interacted with HIS163, whereas the linker group in their structures made H-bond interactions with ASN142, GLY143 and GLN189. For example, the binding mode of ligand-enzyme complex in the PDB structure 5RH3, the corresponding H-bond evolution plots and energy decomposition graph are all provided in the supplementary figure SFig. 9. Unlike the pyridine-based compounds, the ligand co-crystallized in the PDB structure 5RGV, 2-(isoquinolin-4-yl)-N-phenylacetamide, also had a high affinity of −22.65 kcal/mol; nevertheless, in this structure, it was an isoquinoline group that formed a H-bond with HIS163 and occupied the lateral pocket. This shows that the lateral pocket is able to engage different moieties such as pyridine, phenol and isoquinoline group.

**Fig. 7.**
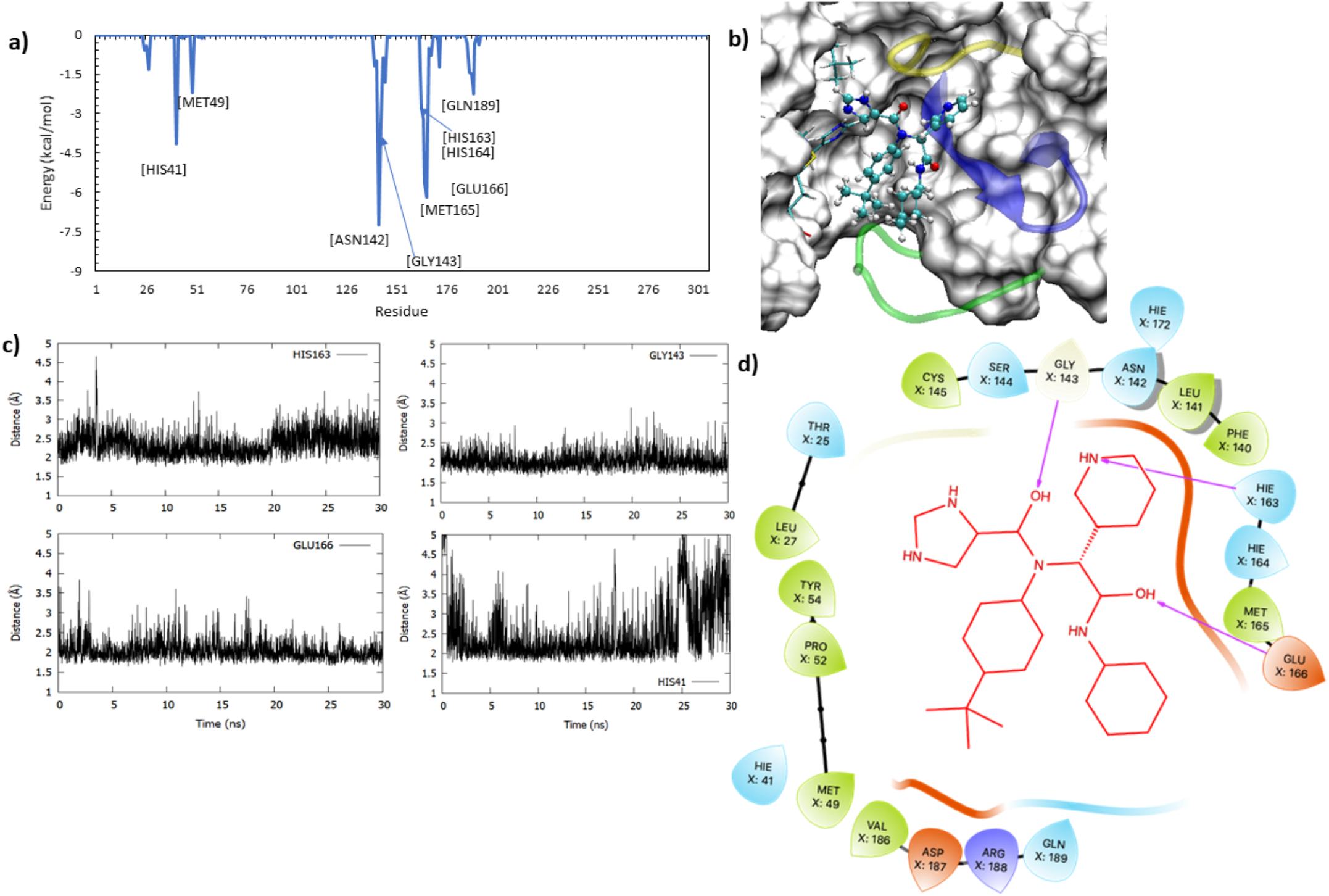
The pair-wise decomposition analyses (a) for the binding mode of ligand in 6W63 complex (3) and its hydrogen bond interactions with the key residues in the pocket (3), along with the 2D interaction diagram for the ligand-M^pro^ binding mode are shown. The decomposition plot (a) identified the key residues such as HIS41, GLY143, GLU166 and HIS163 that contribute to the ligand-protein complex through stable H-bond interactions with the ligand as shown in the H-bond evolution plots (c) and the 3D (c) and 2D (d) interaction diagrams.

**Fig. 8.**
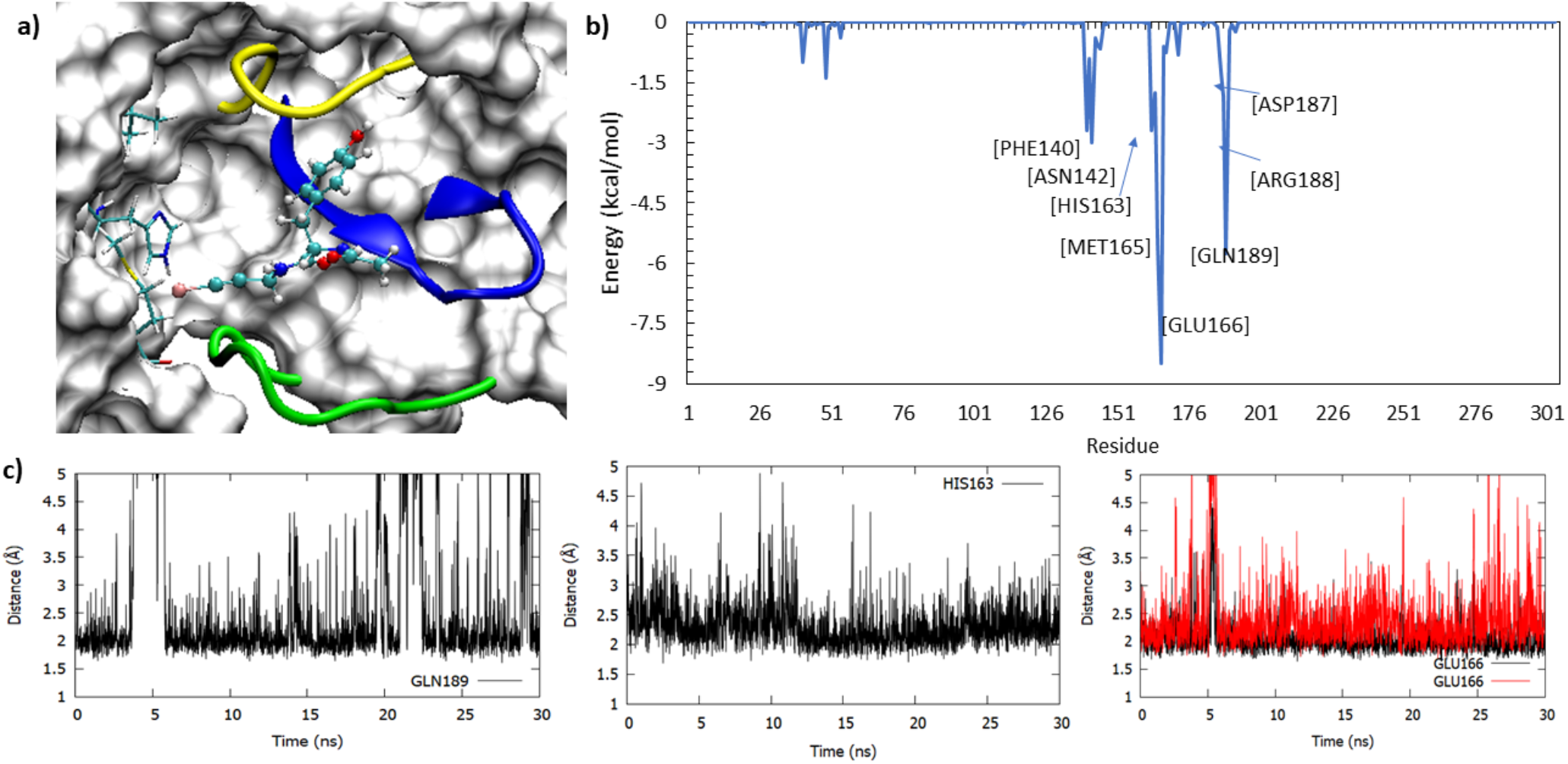
The binding mode of ligand in the 5RG1 complex within the catalytic site of M^pro^ (a), the decomposition of key residues contributing to this ligand-receptor complex (b), along with the time evolution of H-bond interactions of the bound ligand with residues such as GLN189, HIS163 and GLU166 (c). The ligand occupied the lateral pocket in the catalytic site of Mpro, where it formed stable H-bonds with HIS163 residue. This binding mode was supported by other key residues such as GLN189 and 166.

The other molecules in Fig. 6 in the binding affinity range of −15 and −22 kcal/mol include ligands that formed stable interactions with the ‘L2’ loop, ‘L’ β-sheet and also with ‘L1’ loop. However, did not occupy the lateral cavity. For example, the 1-methyl-3,4-dihydro-2~{H}-quinoline-7-sulfonamide ligand (in 5R81), interacted with the ‘L2’ loop through hydrogen bonds with GLN190 and GLY192, and with the ‘L’ β-sheet through via hydrophobic interactions with MET165 and GLU166, in addition to the interactions with HIS41 (supplementary figure, SFig. 10). However, this ligand did not engage with the lateral pocket and consequently had a weaker binding affinity of −18.3 kcal/mol. In the similar nature, ligands including 5-(1,4-oxazepan-4-yl)pyridine-2-carbonitrile (5RF6), N-[(4-cyanophenyl)methyl]morpholine-4-carboxamide (5REE), 2-(4-methylphenoxy)-1-(4-methylpiperazin-4-ium-1-yl)ethenone (5RE9), N-(4-methoxypyridin-2-yl)-2-(naphthalen-2-yl)acetamide (5RGY), and N-(2-phenylethyl)methanesulfonamide (5R7Y) interacted with different segments within the binding site without occupying the lateral cavity and all of them displayed weaker binding affinity toward M^pro^. While our MM-GBSA analyses suggest that ligand engagement with the lateral pocket in Mpro improves the binding affinity; however, the ligand complex of 5RF7 was an exception. The ligand, 1-(4-methylpiperazin-1-yl)-2-(1H-pyrrolo[2,3-b]pyridin-3-yl)ethan-1-one, in this complex contains a pyrrolopyridine moiety that indeed bound into the lateral pocket in the M^pro^ binding site and formed H-bonds with HIS163. However, the methylpiperazine group on the other hand remained flexible and formed only transient H-bonds with the key residues such as GLY143. ASN142 and GLU166 (supplementary figure, SFig. 11). However, it can be noted that although this ligand did not make stable interactions with the key residues in the catalytic pocket such as MET49, GLY143, SER144, MET164, and GLU166 (Fig. 9b), it still had an affinity of ~-16.1 kcal/mol that was mainly due to the stable engagement of the ligand with the lateral pocket in M^pro^. Again, this highlights the important role of the lateral pocket in the ligand M^pro^ interactions.

**Fig. 9.**
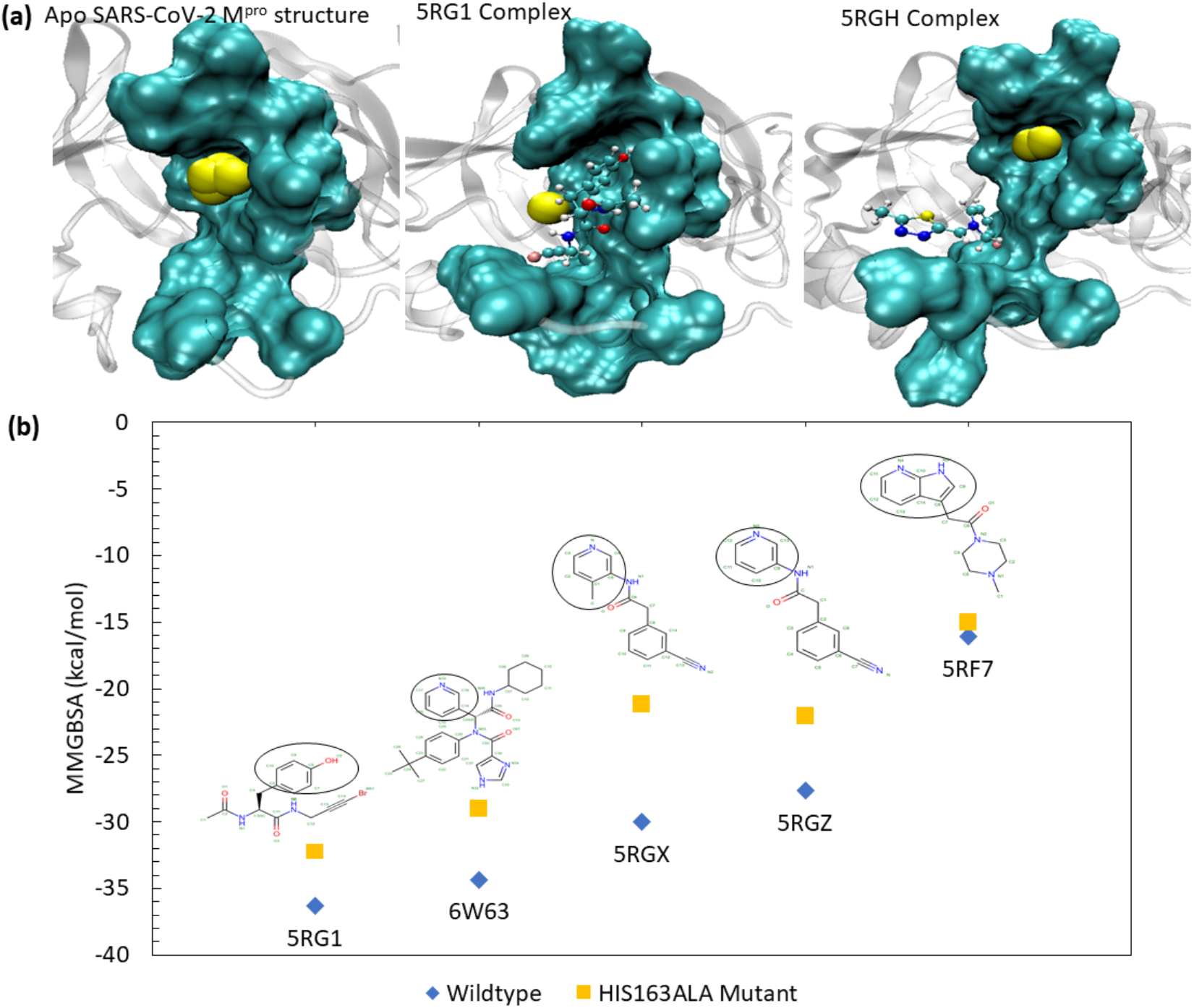
The identification of steric water sites in the apo and ligand-bound M^pro^ complexes (a) along with the comparison of the binding free energy of the wildtype (WT) and HIS163ALA mutant complexes of the select systems (b). (a) The presence of steric water molecules (shown as yellow spheres) are found within the lateral binding pocket in the apo protein structure. When a ligand is in position to occupy the lateral site, the steric water molecules are displaced to the edge of the pocket, allowing for more stable ligand binding. Nevertheless, when the ligand does not occupy the lateral pocket, the steric water molecules are again observed with this site. (b) Mutation of HIS163 to an alanine (ALA163) clearly lead to weak binding affinity between the selected ligand-M^pro^ complexes.

### 3.4. Steric water site analyses in the lateral pocket of M^pro^

Since our MD and MM-GBSA analyses clearly highlighted the role the lateral pocket in the ligand-Mpro affinity, we analyzed the steric water site in the apo M^pro^ structure. This lateral pocket is composed of residues ASN142, GLY143, SER144, CYS145, HIS163, HIS164, MET165, and GLU166. Our analyses revealed that this pocket, in the absence of a ligand, was occupied by steric water molecules (Fig. 9a). Next, we analysed the presence of this steric water in the lateral pocket when a ligand was bound in M^pro^. For example, in the PDB structure of 5RG1, the ligand occupied the lateral pocket and the steric water site was then shifted to the edge of the pocket (Fig. 9a). This displacement of water allowed the ligand to make stable interactions in the pocket and bind the target with enhanced binding affinity. In the case of ligands that did not fully occupy the lateral pocket, for example PDB 5RGH, the binding affinity was seen to be weaker and, now, the steric water molecule site was again found centered within the lateral pocket (Fig. 9a). This site likely favours the presence of an electronegative hydrogen bond forming moiety in whose absence the space is occupied by a water molecule that substitutes this role. This evidence supplements the significant role of the lateral pocket in ligand engagement and stronger binding energies.

### 3.5. Computational mutagenesis of HIS163 in the lateral pocket

Our MD, MM-GBSA and steric water site analyses clearly highlighted the importance of the lateral pocket in M^pro^, in particular the role of HIS163 in stabilizing the ligand-M^pro^ complexes. Therefore, in order to evaluate this finding, we mutated HIS163 with ALA163 (called the HIS163ALA) in the selected M^pro^ complexes and reperformed 30 ns long MD and MM-GBSA calculations as carried out earlier for the wild-type (WT) complexes. For this mutagenesis calculations, we selected five complexes (listed in the supplementary table, ST. 4) related to the PDB structures 5RGZ, 5RF7, 6W63, 5RG1, and 5RGX and mutated HIS163 with ALA163 in the protein structure of each of these complexes. These ligand structures and their wildtype and mutated MMGBSA binding affinities are shown in Fig. 9b. As can be seen in this figure and the supplementary table, ST. 4, the HIS163ALA mutation caused an MM-GBSA difference of up to −9 kcal/mol. For example, the PDB structure of 5RGX complex showed the greatest difference in binding affinity with a decrease of −8.868 kcal/mol. In this complex, a methyl-attached pyridine ring was occupying the lateral pocket. In the WT complex, shown in supplementary figure, SFig. 12, the nitrogen atom of the pyridine ring formed a strong hydrogen bond with HIS163. GLY143, SER144, and CYS145 all formed electrostatic interactions with the double-bonded oxygen atom of the ligand, and ASN142 and MET165 greatly contributed to the ligand’s binding affinity via Van der Waals interactions. In the HIS163ALA mutant complex, the nitrogen atom was no longer able to form a stable interaction with HIS163 (SFig. 12b), dropping the pairwise residue energy from −3.68 kcal/mol to −0.76 kcal/mol and enabling the pyridine ring to become much more flexible throughout the trajectory. In additional to the HIS163 interaction loss, the 5RGX mutant complex also lost its strong ASN142, GLY143, and SER144 interactions, instead formed transient contacts with CYS145 and GLU166. This drastic change in pairwise residue contribution accounted for the −8.868 kcal/mol loss in MM-GBSA binding affinity, as clearly shown in SFig. 12c.

The 5RGZ HIS163ALA mutant complex also showed a significant decrease in binding affinity. This complex had an initial MM-GBSA of −27.6558 kcal/mol which was decreased by −5.582 kcal/mol in the mutated form. The ligand structure in the 5RGZ complex is very similar to that of the 5RGX complex as previously described, with the absence of a methyl group attached to the pyridine ring positioned in the lateral pocket (Supplementary figure, SFig. 13). Although these ligands are very similar in their chemical structures, they behaved differently when subjected to the HIS163ALA mutation. The 5RGZ mutant complex initially adopted a pose much like its WT complex. During the first 15 ns of the MD trajectory, the pyridine ring was found well within the lateral pocket. After 15 ns of simulation, the 5RGZ complex adopted a new pose in which the pyridine ring moved significantly out of the lateral pocket to instead interact with LEU27, HIS41, SER46, and MET49 (SFig. 13).

Complexes 5RG1 and 6W63 (the top MM-GBSA scoring complexes in this study) were also mutated due to their initially strong binding affinities with MM-GBSA values of −36.3341 and - 34.3563, respectively. The 5RG1 complex contained a phenol group in the lateral binding pocket, where the oxygen atom formed a strong H-bond with HIS163. The 6W63 complex contained a very large ligand, with a pyridine ring moiety located in the lateral pocket, as discussed earlier. In addition to its strong interaction with HIS163, in the WT complex, the 5RG1 complex formed stabilizing electrostatic interactions with GLU166 and GLN189. When the histidine residue was mutated, the MM-GSBA binding affinity of the 5RG1 complex decreased by −4.172 kcal/mol. This is attributed to the loss of the HIS163 interaction with the phenol group and GLN189 interaction. In the mutated complex, GLU166 maintained its hydrogen bonds with the ligand, and formed an additional, but transient, electrostatic interaction with the mentioned phenol group. This additional interaction to stabilize the phenol group was seen in the pairwise decomposition graph in Supplementary figure, SFig. 14 and also reflected in the slightly lessened decrease in MM-GBSA binding affinity in comparison to other complexes such as 5RGX and 5RGZ. In the WT 6W63 complex, the pyridine ring moiety and nearby double-bonded oxygen atom formed strong, electrostatic interactions with HIS163, ASN142, and GLY143. When mutated, the double-bonded oxygen atom lost its stabilizing electrostatic interactions, and the HIS163 H-bond with nitrogen pyridine ring atom was replaced by a H-bond with CYS145, decreasing the binding affinity of the complex by −5.339 kcal/mol. This representation can be seen in Supplementary figure, SFig. 15. Since the 6W63 ligand is a bulkly compound, the complex can be stabilized in the binding site and lateral pocket by its other important electrostatic and Van der Waals interactions.

We also assessed the effects of mutation on the PDB complex of 5RF7 by building a HIS163ALA mutant model. This complex has a unique 7-azaindole, double-ringed moiety that fits in the lateral pocket. In theWT 5RF7 complex, this moiety formed stable H-bonds with both HIS163 and HIS164. The rest of the ligand was unable to form lasting interactions with the protein residues and oscillated between ASN142/GLY143 and GLU166. In the HIS163ALA mutant, although the HIS163 bond was lost, the strong bond with HIS164 was maintained. In addition, the ligand was positioned so that it interacted stronger with GLU166. The gain of this interaction is evident in the residue decomposition graph in Supplementary figure, SFig. 16. Since the mutated 5RF7 complex can maintain key, stabilizing interactions, specifically in the lateral pocket, its MMGBSA binding affinity was only decreased by −1.076 kcal/mol. Therefore, our computational mutagenesis analyses clearly demonstrate the importance of HIS163 and its role in ligand engagement in SARS-CoV-2 Mpro.

### 3.6. Effects of explicit water molecules in the binding affinity of ligand-M^pro^ complexes

Our MM-GBSA analyses revealed several key interactions that contribute to the binding affinity of the reversible ligands against Mpro. It is known that water molecules play an important role in the catalytic process of Mpro in SARS-CoV-2, as also confirmed by our steric water site analyses. Therefore, we analyzed the effects of explicit water molecules on the binding free energies of the M^pro^ complexes, in which the ligand was bound within the catalytic binding site (or pocket 1). For this purpose, we employed the NWAT-MMGBSA variant, as described earlier by Maffucci et al^65^ and explained in the methods section. We estimated the MM-GBSA energies with the presence of 0 to 6 explicit water molecules (noted as NWAT=0 to 6 in the text). The calculated NWAT-MM-GBSA values for the complexes studied are provided as a polar plot in Fig. 10a and the corresponding values are listed in the supplementary table, ST.5. In the polar chart (in Fig. 10a), angular axis (along the circle) presents the PDB codes of the complexes studied; and the radial axis (connecting the circle to the center of the plot) presents the binding affinity values in kcal/mol. The MM-GBSA energies with no explicit water (i.e., NWAT=0) for the systems is the outer polar line shown in blue. The MM-GBSA energies with the presence of (1 to 6) explicit water molecules are also shown as data points connected by polar lines in different colors. For any given system, if the presence of a water molecule enhances the affinity, then the polar data points for that system is expected to move inside towards the center. Whereas, if the water molecule did not make any impact, then the polar data points are expected to cluster at the same position. As can be seen in Fig. 8, we noticed that most of the systems had the impact of explicit water molecules, except in some cases. The complexes represented by the PDB structures 5RHD, 5R7Y, 5R81, 5RGZ, and 5RG1 clearly had a significant impact by the presence of explicit water molecules, as the data points for these systems moved inward with the increase of each water molecule. This suggests that the increase in binding affinity (lower MM-GBSA scores) with increasing numbers of waters. For example, the maximum impact of water molecules on binding affinity was seen for the 5RHD complex, where the MM-GBSA scores dropped by ~8 kcal/mol (from −13.63 for NWAT=0 to - 22.09 for NWAT=6). Analyzing the structures (in Fig. 10b), it can be noted that the water molecules bridged the bound ligand (1-[4-(methylsulfonyl)phenyl]piperazine) with CYS44 and HIS41 residues during MD. The other water molecules were generally found engaging with the sulfonyl and piperzine groups. Similarly, for the high affinity complexes such as 5RGZ and 5RG1, the inclusion of explicit water molecules further enhanced their affinity towards M^pro^ (Fig. 10a). As expected, the water molecules were found to stabilize the electronegative sites such as amide, nitriles, or the hydroxyl groups in the ligands co-crystallized with these complexes (Fig. 10c-d). In some other complexes such as these represented by the PDB codes 5RF7, 5RGK, 5RH8, 5RGV, adding one of two water molecules helped to enhance the ligand-enzyme affinity; however, addition of more water molecules did not make any more impact in the affinity. As a result, the MM-GBSA scores for these complexes with NWAT >2 all cluster at one position suggested the saturation in the number of water molecules that can engage inside the catalytic site of M^pro^. Thus our NWAT-MM-GBSA analyses suggest that upto 2 water molecules could play an important role in the interactions of ligand-M^pro^ in the complexes studied in this work. Nevertheless, we were also able to see that some complexes did not have any impact from the explicit water molecules such as the complexes of 5RGX and 5RE9, as the ligand bound in these complexes were already engaged in optimal interactions with the residues in the pocket, therefore, the water could not contribute to the ligand-protein interactions (supplementary figure, SFig. 17).

**Fig. 10.**
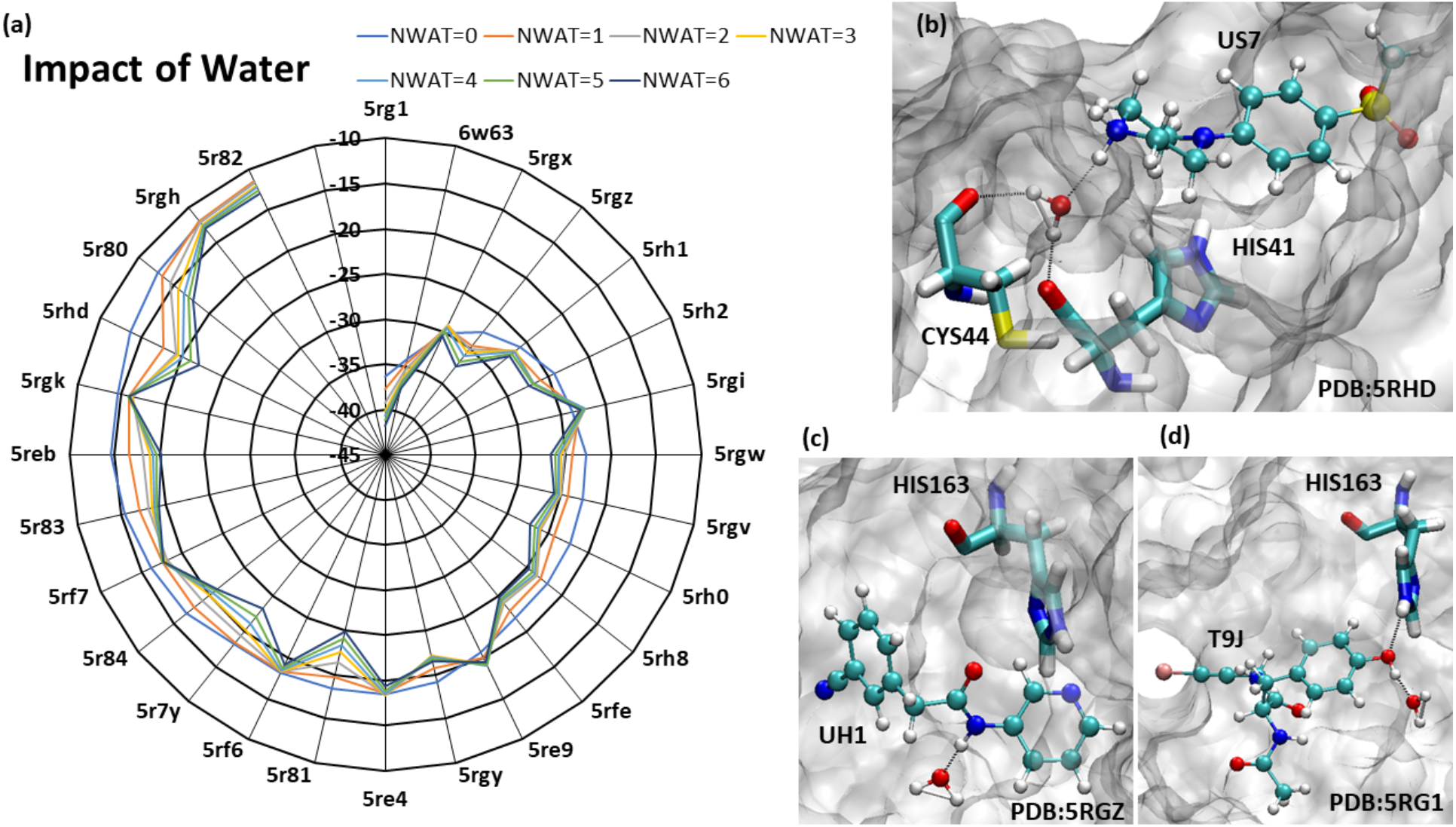
A polar plot describing change in energy affinity with the presence of varying numbers of explicit water molecules (NWAT=0 to 6) (a), and the different water interactions with the ligand (b-d). Water bridge in 5RHD, water serve as proton donor to M^pro^ and proton acceptor for the ligand (b). In 5RGZ and 5RG1, water forms 1 stable interaction with carboxamide and hydroxyl side chain of the ligand to keep ligand within lateral pocket (c-d).

### 3.7. Ligand interactions with a M^pro^ dimer

As we discussed above some of the ligands bound to the monomer form of M^pro^ did not exhibit stable interactions in MD and egressed the protein during the course of simulation. It is now known that M^pro^ is functional in a dimeric state, where the N-finger of the second monomer interacts with the first monomer chain to close its catalytic site. Therefore, the natural question is if the ligands egressed M^pro^ during our simulation where due to the absence of the involvement of the second chain of M^pro^. Therefore, in order to address this concern, we analysed the dynamic interactions of the select ligands in the dimeric state of M^pro^. These include the ligands that were originally bound (1) within the catalytic site of M^pro^ and unbound during MD, (2) at the dimer interface of M^pro^ (e.g., 5RGQ and 5RFA complexes), and (3) at the proximity to the dimer interface. Using these selection criteria, a total of 13 complexes were selected for evaluation in dimer system. Fig. 11 presents the positions of all the selected ligands within a dimer structure of M^pro^ and the PDB codes for the corresponding ligands are also marked in this figure. For each of the complexes selected, we modelled the dimer model by aligning a ligand-bound monomer against the PDB structure of a SARS-CoV-2 M^pro^ dimer (PDB 6WTT), and deleting an aligned chain of monomer. The modelled ligand-bound dimer complexes were subjected to 30 ns long MD simulation and their binding free energies were recalculated using the MM-GBSA method. The MM-GBSA scores of the select complexes both under monomer and dimer conditions are compared in the supplementary table, ST.6. We initially assessed the stability of the complexes, in which the ligands were bound at the proximity to the dimer interface. These include 5RF1, 5RE8, 5RF9, 5RE7, 5RGJ,5REZ (refer to ligand positions in Fig. 11a). As can be seen in the supplementary table, ST.6, the presence of a M^pro^ dimer does not increase the stability or affinity of these complexes when compared to their respective monomer M^pro^ complexes. These ligands still unbound from the dimer M^pro^ structures. This highlights the likely weak affinity of these ligands towards M^pro^. Subsequently, we analyzed the complexes, in which the ligands were originally bound to the catalytic site (pocket 1) and subsequently egressed the monomer Mpro during MD. This set of complexes includes the PDB structures of 5REH, 5RF2, 5RF3, 5RF8 and 5R7Z. Again, as can be seen in the supplementary table, ST. 5, the engagement of the second M^pro^ chain through its N-finger with that of the ligand-bound chain one M^pro^ did not help in improving the stability and affinity of the most of these ligands. For example, the presence of the second M^pro^ chain did not improve the affinity of the two smallest fragments bound at the binding site in the monomer one of M^pro^ such as 1-azanylpropylideneazanium (in 5RF2) and pyrimidin-5-amine (in 5RF3). This is not surprising given their small size. However, the only exception to this is the ligand 1-cyclohexyl-3-(2-pyridin-4-ylethyl) urea (PDB: 5REH) in Fig. 12a, whose affinity towards a M^pro^ dimer increased by ~-14 kcal/mol when compared to the monomer. The stability of the ligand with the dimer over the monomer is described by the ligand RMSD fluctuation in both the states (in Fig. 12b). Analyzing the MD trajectory of ligand-bound dimer complex, we noticed that the binding of N-finger of the second monomer against the monomer A of M^pro^ closed the binding site, thereby, pushing the ligand further into the binding site. In fact, the closure of the catalytic pocket by the N-finger from the second M^pro^ chain pushed the pyridine ring of the ligand further into the lateral pocket to establish the crucial H-bond interactions with HIS163 (in Fig. 12c). Further, the amine groups in the ligand made stronger H-bonds with GLU166 (Fig. 11c) that helped to improve the stability and affinity of this complex.

**Fig. 11.**
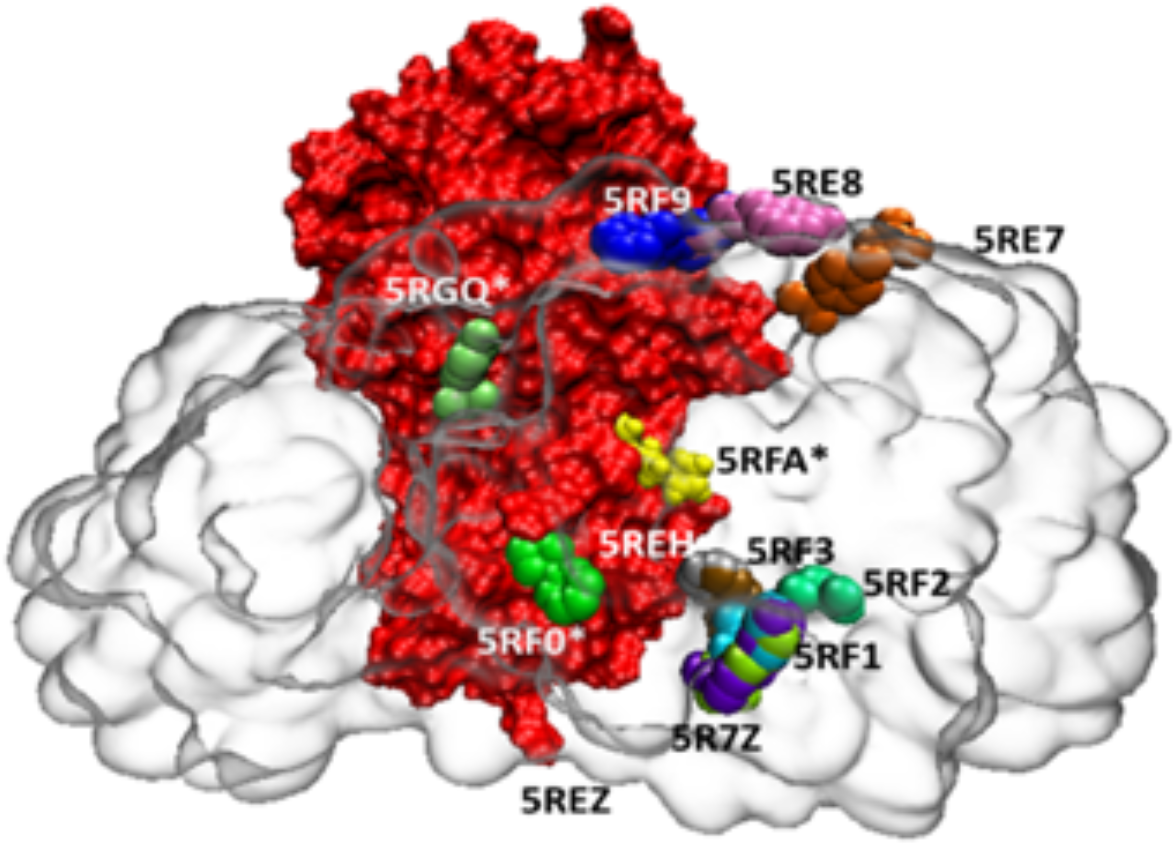
A surface representation of a dimer model of M^pro^ (red surface is monomer 1 and transparent surface is monomer 2) showing the interactions of ligands (shown as spheres) that are bound at different sites at proximity to the dimer interface. The PDB codes of the ligands are marked. The ligands binding at the dimer interface is marked by *.

**Fig. 12.**
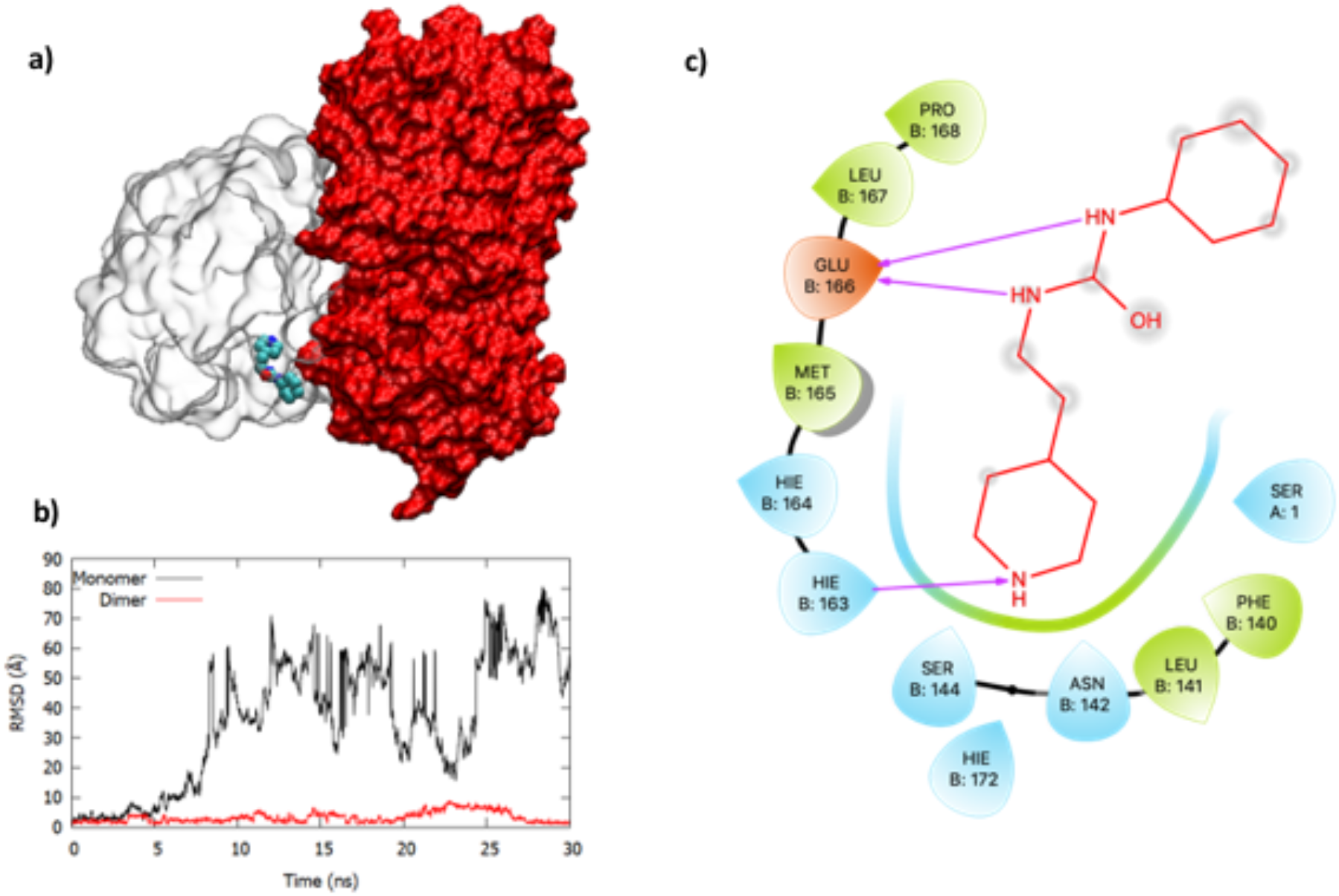
The binding of ligand from 5REH within a dimer model of M^pro^ (shown as surface representation) (a) shown, along with the fluctuation of the ligand when bound with the monomer and dimer (b) and the binding pose of the ligand within the dimer. The RMSD plots describe that the ligand was unstable when bound to a monomer of M^pro^; however, it was more stable within a dimer condition. This is likely due to its favourable binding in the active site containing pocket where the ligand made stable hydrogen bonds with HIS163 and GLU166 of monomer B thus occupying the lateral pocket that was extensively explored by other ligands part of our set (c). However, it is seen that the N terminal of the second monomer, specifically SER 1, interacts in a hydrophobic manner, which indeed helped in the stable ligand interactions with M^pro^.

Finally, the ligands that benefitted the most by the presence of the Mpro dimer where obviously the ligands that bound at the dimer interface such as 1-(4-fluoro-2-methylphenyl)methanesulfonamide (5RGQ), 1-methyl-N-{[(2S)-oxolan-2-yl]methyl}-1H-pyrazole-3-carboxamide (5RFA), and [1-(pyridin-2-yl)cyclopentyl]methanol] (5RF0). While the methanesulfonamide ligand (in 5RGQ) remained stable even within the monomer structure, this was not the case with the other two ligands. Nevertheless, the presence of a dimer environment assisted in the stability of these ligands and improved their binding affinity with M^pro^ significantly. For example, the carboxamide-based ligand (in 5RFA) was bound at the interface of two M^pro^ monomers (Fig. 13a) and remained stable throughout the course of MD simulation, which was not true when it was bound with a monomer enzyme, as described by the ligand RMSDs in Fig 13b. However, within the dimer form, the ligand underwent some structural changes around ~10 ns during MD and remained in a much stable pose that was supported by the interactions of the ligand with the key residues such as MET6, PHE8, ILE152, ASP298 from one M^pro^ monomer, and predominantly SER123 from the other monomer (Fig. 13c). This pose is described by the 2D interaction diagram shown in Fig. 13d. In a similar nature, the methanol-based ligand (in 5RF0) also displayed enhanced affinity with the dimer when compared to the monomer form (refer to supplementary table, ST. 5). The 3D structure of all the ligand-bound dimer complexes along with their RMSD fluctuations in monomer and dimer states of M^pro^ are provided in the supplementary figure, SFig. 18.

**Fig. 13:**
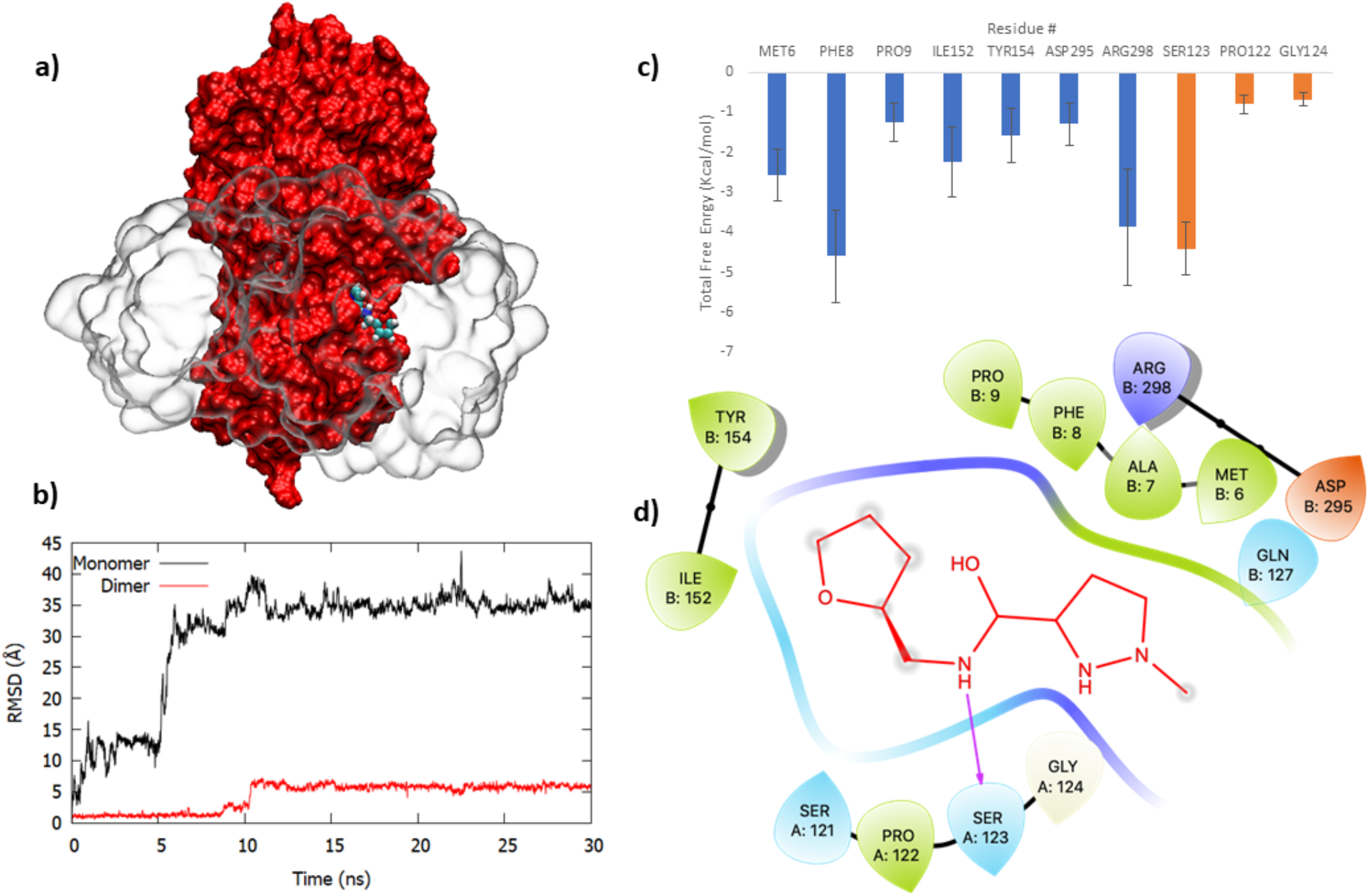
The binding of ligand from 5RFA within a dimer model of M^pro^ (shown as surface representation) (a) shown, along with the fluctuation of the ligand when bound with the monomer and dimer (b), the pairwise energy decomposition plot (c) and the binding pose of the ligand within the dimer (d). The RMSD plots describe that the ligand was unstable when bound to a monomer of M^pro^; however, it was more stable within a dimer condition although it unwent some conformational changes at ~10ns before reaching a pleateau indicating the stability of the ligand after this point (b). The energy decomposition analyses (c) identified a number of key residues from both the Mpro monomer chains to stabilze the ligand interactions, as described in the 2D interaction diagram (d).

Since dimerization of M^pro^ is a crucial mechanism for the activity of the enzyme and hence the replication of SARS-CoV-2 enzyme, there is an increasing interest towards disrupting the dimerization of Mpro. Therefore, the molecular level interactions of the ligands at a specific pocket present at the dimer junction identified through earlier X-ray crystal screening efforts^70^ and currently studied through our efforts using MD and MM-GBSA analyses could be useful for developing new compounds or identifying known drugs with physicochemical complementarity to this site. Identifying such compounds will allow us to explore the widely acknowledged novel approach of targeting the protease activity in SARS-CoV-2.

## Conclusion

The novel coronavirus pandemic caused by the SARS-CoV-2 virus has emerged as a huge challenge for the 21^st^ century. In less 10 months of outbreak, this infection has already caused over 46 million cases and 1.16 million deaths worldwide. Absence of a vaccine or a drug against this virus remains a serious concern. The Mpro enzyme in SARS-CoV-2 has been identified as a potent drug target as it plays a vital role in regulating the replication and transcription of the virus. With consistent scientific efforts, several high-resolution 3D structures of ligand-Mpro complexes have already been reported in the PDB and this number is rapidly growing. These structures offer useful insights about the ligand interactions with Mpro and it is important to understand the dynamic interactions of these complexes. In this work, we employed a wide array of computational methods including MD simulation, binding free energy calculation using implicit and explicit water molecules, pairwise decomposition of free energy, pocket analyses methods, steric water site analyses and computational mutagenesis to present a comprehensive picture about ligand-Mpro interactions at atomic level. We studied all 62 reversible ligand-M^pro^ complexes that were published in the PDB as of June 10^th^, 2020.

Initially, we performed rigours MD and cavity analyses of the apo Mpro structure and identified and characterized the conformational dynamic properties of 8 different pockets including the known catalytic pocket (pocket 1). Ligands in the PDB structures studied in this work bound to almost all of these sites, majority of the ligands were bound to the catalytic site though. Through MD and stability analyses, we revealed that the ligands interacting in the catalytic site of monomer Mpro in general remained stable, except for a few ligands that egressed the protein during MD. We also identified that some ligands bound in the pockets 4 and 3 that are located in the domain I and at the intersection of domain II and III, respectively, displayed some affinity and stability during MD simulations. Nevertheless, all the ligands bound to pockets 5 and 6 were not stable and egressed the binding site during MD simulation. This suggests that either the ligands do not have suitable physicochemical complementary against the pockets or the pockets are not suitable for ligand binding and modulation. Binding free energy analyses revealed that their affinities were always better than the complexes with ligands bound in other pockets.

Through our analyses, we highlight four key components in the catalytic site of Mpro and the binding of ligands with these elements determine their affinity to Mpro. In specific, we identified a lateral pocket, featuring HIS163 as a key residue, which played a critical role in enhancing the stability and the affinity between ligands and Mpro. We revealed that, in the absence of a bound ligand, the steric water molecules occupied the lateral pocket to engage with HIS163. Nevertheless, when the ligand binds at this site, the steric water moves away so as to facilitate the binding of ligands to HIS163, which eventually increased the affinity of ligands to Mpro. *In silico* mutation of this HIS163 to an alanine reduced the affinity of the ligand with Mpro, thereby clarifying the importance of this residue in efficient ligand recognition by Mpro. Further, using NWAT-MM-GBSA variant, we demonstrated that the presence of up to 2-3 explicit water molecules can improve the affinity between certain ligands and the catalytic site in Mpro. Finally, assessed the role of dimerization in Mpro for the binding and stability of some ligands. As widely acknowledged, presence of a dimer might be important for binding certain ligands in the catalytic site of Mpro, but not for all the ligands studied in this work. The ligands studied here were either already stable in the monomer state or unstable in both monomer and dimer complexes. However, a few set of ligands, particularly those bound at the dimer interface in Mpro, required the presence of both the monomer chains to display stability. We discussed several key molecular dynamic interactions of ligands binding at the dimer interface of Mpro.

In summary, our work has highlighted several key molecular determinants that are have critical for ligand binding in Mpro including different pocket characteristics, role of water molecules in the ligand-Mpro interactions, the importance of a lateral pocket and the key electrostatic interactions rendered by HIS163 in this site, and molecular level details about ligand interactions at the dimer interface in Mpro. These comprehensive molecular and mechanistic insights about the crucial Mpro target could be useful to facilitate the identification of known drugs or the rational design of compounds with specific characteristics to exploit all fingerprint hotspots in Mpro and effectively inhibit this enzyme.

## List of abbreviations

M^pro^: Main protease
MD: Molecular dynamics
PDB: Protein data bank
MM-GBSA: molecular mechanics generalized Born surface area
RMSD: Root mean square deviation
H-bond: Hydrogen bond.

## Acknowledgments

This research was undertaken thanks in part to funding from the Canada First Research Excellence Fund to AG and KS. AG acknowledges the support for YLW from BioTalent Canada through the student work placement program. This research was enabled by the support provided by Compute Canada through access to high performance GPU clusters of GRAHAM and Cedar.

## Availability of data and materials

Any data related to this work might be available from the corresponding author.

## Authors’ contributions

AG and KS contributed to the conception and design of the work. YLW, SRN, and ND contributed equally to carrying out all the modelling, simulation and preparation of figures. All authors contributed equally for the analyses of results, findings and writing the manuscript. All the authors read and approve the final manuscript.

## Ethics approval and consent to participate

Not applicable.

## Consent for publication

Not applicable.

