## Supplementary for "Molecular dynamics and in silico mutagenesis on the reversible inhibitor-bound SARS-CoV-2 Main Protease complexes reveal the role of a lateral pocket in enhancing the ligand affinity"

### Supplementary Information

Ying Li Weng[1$], Shiv Rakesh Naik[1$], Nadia Dingelstad[1$], Subha Kalyaanamoorthy[2*], Aravindhan Ganesan[1*]

[1]ArGan's Lab, School of Pharmacy, Faculty of Science, University of Waterloo, Ontario, Canada.
[2]Department of Chemistry, Faculty of Science, University of Waterloo, Ontario, Canada.
[$]Authors contributed equally.

**ST. 1:** Following contains the list of all the 62 reversible ligands that were simulated and analyzed in this study with assigned index numbers and an figure that follows the table showing the stuctures of each compound.

| Index | PDB | Index | PDB | Index | PDB | Index | PDB |
|---|---|---|---|---|---|---|---|
| **1** | 5R7Y | 16 | 5REC | 31 | 5RF4 | 46 | 5RGH |
| **2** | 5R7Z | 17 | 5RED | 32 | 5RF5 | 47 | 5RGI |
| **3** | 5R80 | 18 | 5REE | 33 | 5RF6 | 48 | 5RGJ |
| **4** | 5R81 | 19 | 5REF | 34 | 5RF7 | 49 | 5RGK |
| **5** | 5R82 | 20 | 5REG | 35 | 5RF8 | 50 | 5RGR C1 |
| **6** | 5R83 | 21 | 5REH | 36 | 5RF9 | 51 | 5RGR C2 |
| **7** | 5R84 | 22 | 5REI | 37 | 5RFA | 52 | 5RGS |
| **8** | 5RE4 | 23 | 5RGZ | 38 | 5RFB | 53 | 5RGY |
| **9** | 5RE5 | 24 | 5RH4 | 39 | 5RFC | 54 | 5RGX |
| **10** | 5RE6 | 25 | 5RHD | 40 | 5RFD | 55 | 5RH0 |
| **11** | 5RE7 | 26 | 5REZ | 41 | 5RFE | 56 | 5RH2 |
| **12** | 5RE8 | 27 | 5RF0 | 42 | 6W63 | 57 | 5RH1 |
| **13** | 5RE9 | 28 | 5RF1 | 43 | 5RGG | 58 | 6YVF |
| **14** | 5REA | 29 | 5RF2 | 44 | 5RGQ | 59 | 5RGV |
| **15** | 5REB | 30 | 5RF3 | 45 | 5RG1 | 60 | 5RGW |
| | | | | | | 61 | 5RH3 |
| | | | | | | 62 | 5RH8 |

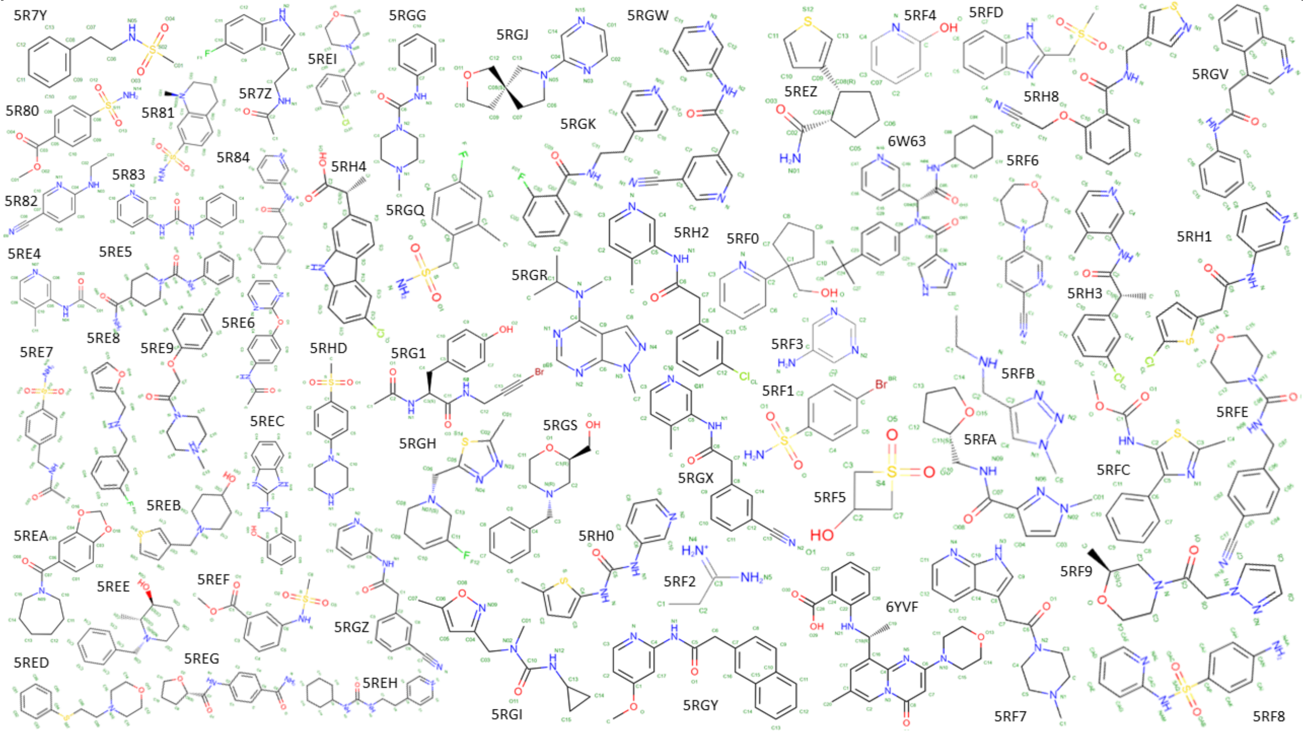

**ST. 2: Apo pocket characterizations.** More details on the pockets identified in the apo protein MD trajectory. Each were characterized by pocket name, residue composition, PDB ID's of ligands bound in their crystal structure positions, and pocket volume.

| Pocket Name | Residues | PDB ID's in crystal structure | Volume |
|---|---|---|---|
| 1 | PHE140 LEU141 ASN142 GLY143 SER144 CYS145 HIS163 HIS164 MET165 GLU166 HIS172 VAL186 ASP187 ARG188 GLN189 GLN192 | | 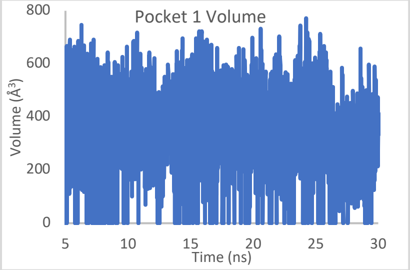 |
| 2 | PHE3 ARG4 LYS5 MET6 PHE8 THR111 GLN127 PHE291 THR292 PHE294 ASP295 VAL296 ARG298 GLN299 GLY302 THR304 | 5RFA<br>5RGQ | 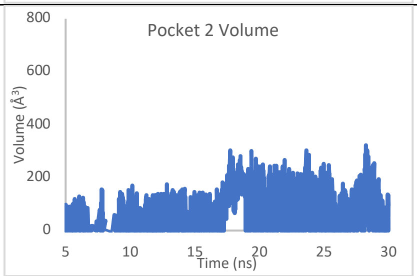 |
| 3 | THR198 MET235 ASN238 TYR239 GLU240 PRO241 | 5REC<br>5RGS<br>5REE | 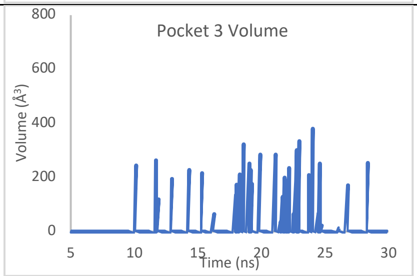 |
| 4 | ASP34 VAL35 VAL36 TYR37 GLY79 HIS80 SER81 LYS88 LEU89 LYS90 | 5RFC<br>5RH4<br>5RE6<br>5RE5<br>5RGG<br>5RFB<br>6YVF | 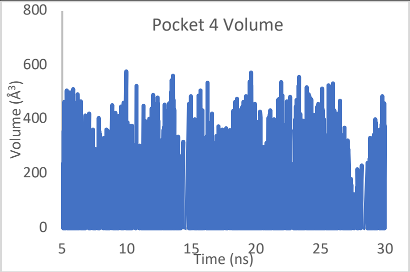 |
| 5 | PHE103 VAL104 ARG105 PHE159 CYS160 TYR161 THR175 ASN176 LEU177 GLU178 ASN180 TYR182 | 5REI<br>5RED<br>5RF5<br>5RGR | 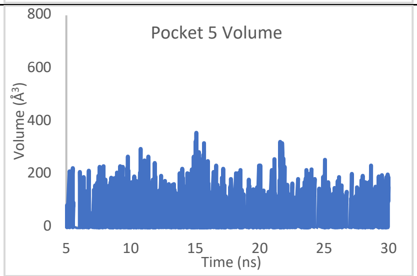 |

| | | | |
|---|---|---|---|
| 6 | PHE8 PRO9 SER10 GLY11 LYS12 VAL13 GLU14 GLY15 CYS16 MET17 LEU30 LEU32 ASP33 ASP34 ILE78 LYS90 VAL91 ASP92 THR93 ALA94 ASN95 PRO96 LYS97 THR98 PRO99 LYS100 TYR101 LEU115 VAL148 PHE150 ASN151 ILE152 ASP155 CYS156 VAL157 PHE159 | 5RF4<br>5RFD<br>5RE8<br>5RF9<br>5RGJ | 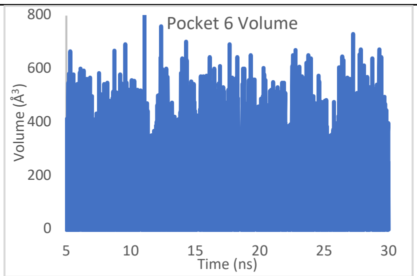 |
| 7 | ARG105 ILE106 GLN107 PRO108 MET130 PHE134 PHE181 TYR182 GLY183 PRO184 | 5REG | 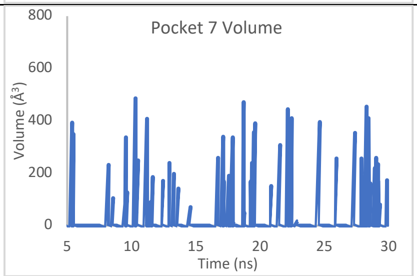 |
| 8* | LYS5 ILE106 GLN107 PRO108 GLY109 GLN110 THR111 PHE112 VAL114 LEU115 ALA116 TYR118 SER123 GLY124 VAL125 TYR126 GLN127 CYS128 ALA129 MET130 ARG131 PRO132 ASN133 PHE134 THR135 ILE136 LYS137 GLY138 SER139 PHE140 SER147 VAL148 GLY149 CYS160 TYR161 MET162 HIS163 LEU167 PRO168 THR169 GLY170 VAL171 HIS172 TYR182 ALA193 ALA194 GLY195 THR196 ASP197 THR198 THR199 ILE200 THR201 VAL202 ASN203 VAL204 LEU205 TYR237 ASN238 TYR239 GLU240 PRO241 LEU242 THR243 ASP245 HIS246 ILE249 LEU250 LEU287 GLU288 ASP289 GLU290 PHE291 THR292 PRO293 PHE294 ASP295 | 5RF0 | 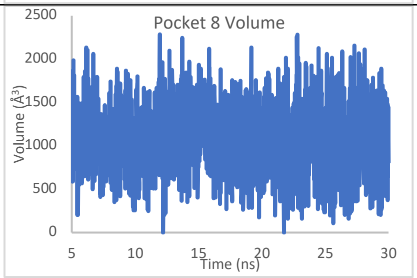 |

***The figure here depicts residues forming pocket 8 in yellow surface representation, red and blue represents monomer A and B respectively.**

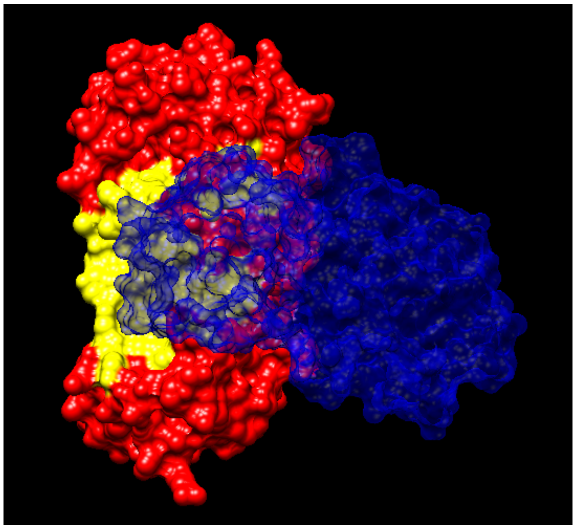

**ST. 3:** Below presented are all the 62 reversible ligands according to their crystal structure complexes, presented alongwith their MM-GBSA Binding Free Energy and averaged Protein Backbone RMSD and Unaligned Ligand RMSD. The first section of the table presents all the ligands in the active site/Pocket1 and the section thereafter the ligands are grouped according to their binding pocket.

| Index | PDB | MMGBSA (kcal/mol) | Protein Average RMSD(Å) | Standard Deviation | Ligand Average RMSD(Å) | Standard Deviation | Pocket |
|---|---|---|---|---|---|---|---|
| **1** | **5R7Y** | **-18.12** | **1.916665** | **0.338393** | **4.350324** | **0.871391** | **1** |
| **2** | 5R7Z | -7.5255 | 2.082408 | 0.380917 | 10.36614 | 6.734171 | 1 |
| **3** | 5R80 | -12.7838 | 1.776822 | 0.350001 | 4.838292 | 1.563662 | 1 |
| **4** | 5R81 | -18.3757 | 1.498825 | 0.190576 | 3.676136 | 1.165353 | 1 |
| **5** | 5R82 | -11.6298 | 1.657093 | 0.227184 | 5.620554 | 1.534471 | 1 |
| **6** | 5R83 | -15.3713 | 1.856718 | 0.369908 | 2.306814 | 0.735295 | 1 |
| **7** | 5R84 | -16.8059 | 1.955554 | 0.31836 | 2.53541 | 1.224635 | 1 |
| **8** | 5RE4 | -18.4114 | 1.617992 | 0.329689 | 3.142076 | 0.518411 | 1 |
| **13** | 5RE9 | -20.6948 | 1.693995 | 0.296101 | 1.855428 | 0.988273 | 1 |
| **15** | 5REB | -14.65 | 2.128578 | 0.415933 | 7.586501 | 3.174916 | 1 |
| **21** | 5REH | 0.0032 | 1.743523 | 0.327849 | 35.866791 | 21.567743 | 1 |
| **23** | 5RGZ | -27.6558 | 1.700701 | 0.237055 | 2.42632 | 0.186166 | 1 |
| **25** | 5RHD | -13.6088 | 2.026874 | 0.399533 | 3.206183 | 0.70888 | 1 |
| **26** | 5REZ | -11.56 | 1.419772 | 0.175804 | 5.63 | 3.322453 | 1 |
| **29** | 5RF2 | -0.0124 | 1.815365 | 0.261984 | 59.83 | 23.908938 | 1 |
| **30** | 5RF3 | -0.1652 | 1.848161 | 0.455702 | 20.52 | 18.616497 | 1 |
| **33** | 5RF6 | -18.2075 | 1.52232 | 0.205489 | 3.98 | 0.845343 | 1 |
| **34** | 5RF7 | -16.1116 | 1.680846 | 0.296371 | 4.08 | 1.04664 | 1 |
| **35** | 5RF8 | -4.4487 | 1.44285 | 0.205282 | 7.64 | 6.504823 | 1 |
| **41** | 5RFE | -21.9365 | 1.827949 | 0.336063 | 6.03 | 0.829553 | 1 |
| **42** | 6W63 | -34.3563 | 1.756177 | 0.371969 | 2.1 | 0.523618 | 1 |
| **45** | 5RG1 | -36.3341 | 1.506031 | 0.29765 | 1.2044 | 0.376415 | 1 |
| **46** | 5RGH | -12.1294 | 1.671961 | 0.194733 | 3.571218 | 1.199408 | 1 |
| **47** | 5RGI | -23.8108 | 1.78312 | 0.380094 | 1.507002 | 0.403063 | 1 |
| **49** | 5RGK | -14.5249 | 1.672967 | 0.287044 | 4.005973 | 2.589809 | 1 |
| **53** | 5RGY | -19.1657 | 1.80972 | 0.336406 | 3.248273 | 0.851531 | 1 |
| **54** | 5RGX | -30.0259 | 1.626892 | 0.190424 | 2.423812 | 0.779939 | 1 |
| **55** | 5RH0 | -22.2447 | 1.671874 | 0.268831 | 1.549979 | 0.529899 | 1 |
| **56** | 5RH2 | -24.2701 | 1.68253 | 0.401635 | 1.893947 | 0.660018 | 1 |
| **57** | 5RH1 | -25.9355 | 1.485169 | 0.257258 | 4.529707 | 2.265744 | 1 |
| **59** | 5RGV | -22.6488 | 1.797764 | 0.265732 | 2.113314 | 0.48039 | 1 |
| **60** | 5RGW | -22.7985 | 2.034154 | 0.422069 | 1.564342 | 0.555704 | 1 |
| **61** | 5RH3 | -24.488 | 1.556123 | 0.22126 | 1.26012 | 0.42493 | 1 |
| **62** | 5RH8 | -22.1462 | 1.545132 | 0.266675 | 3.984186 | 0.732088 | 1 |

| | | | | | | | |
|---|---|---|---|---|---|---|---|
| **Ligands initially bound in pockets other than pocket 1 are grouped below according to their initial site of binding.** | | | | | | | |
| **44** | 5RGQ | -18.3015 | 1.711714 | 0.369385 | 2.288655 | 0.46103 | 2 |
| **37** | 5RFA | -10.8007 | 1.413174 | 0.26024 | 30.38 | 9.114598 | 2 |
| **16** | 5REC | -17.5638 | 1.852156 | 0.277849 | 3.644229 | 0.731061 | 3 |
| **18** | 5REE | -2.0283 | 1.72434 | 0.283528 | 36.141055 | 12.469357 | 3 |
| **52** | 5RGS | -14.7914 | 1.887242 | 0.303038 | 5.919377 | 0.626652 | 3 |
| **9** | **5RE5** | **-16.2026** | 2.844302 | 0.405643 | 31.10255 | 18.156573 | 4 |
| **10** | **5RE6** | **-12.056** | 1.667771 | 0.215722 | 38.135293 | 24.649068 | 4 |
| **38** | **5RFB** | **-6.844** | 1.914196 | 0.595251 | 38.7 | 22.077026 | 4 |
| **39** | 5RFC | -10.6264 | 1.860463 | 0.280546 | 6.05 | 2.082824 | 4 |
| **43** | **5RGG** | **-10.0251** | 1.370923 | 0.153117 | 21.941146 | 16.319402 | 4 |
| **24** | **5RH4** | **-17.5026** | 1.920799 | 0.283871 | 3.48053 | 0.734264 | 4 |
| **58** | **6YVF** | **-15.3489** | 1.85554 | 0.213391 | 3.440421 | 1.159273 | 4 |
| **22** | 5REI | -7.7987 | 1.47849 | 0.213572 | 22.50945 | 13.849877 | 5 |
| **17** | 5RED | -1.4305 | 1.564717 | 0.249023 | 49.816839 | 16.664439 | 5 |
| **32** | 5RF5 | 0.3446 | 2.247144 | 0.435769 | 35.11 | 15.83506 | 5 |
| **50** | 5RGR | -3.0798 | 1.899464 | 0.338068 | 21.966997 | 15.395446 | 5 |
| **51** | 5RGR | -9.0979 | 1.800823 | 0.21708 | 24.34032 | 21.539441 | 5 |
| **58** | 6YVF | -15.3489 | 1.85554 | 0.213391 | 3.440421 | 1.159273 | 5 |
| **31** | 5RF4 | -3.6433 | 1.764083 | 0.333263 | 63.18 | 18.850572 | 6 |
| **12** | 5RE8 | -0.1094 | 1.672967 | 0.317862 | 47.158758 | 14.972364 | 6 |
| **36** | 5RF9 | -8.414 | 1.902701 | 0.369736 | 24.87 | 8.163983 | 6 |
| **40** | 5RFD | -1.5178 | 1.359467 | 0.24306 | 40.74 | 13.492713 | 6 |
| **48** | 5RGJ | -6.6131 | 1.812472 | 0.402485 | 45.739098 | 10.068594 | 6 |
| **20** | 5REG | -18.2217 | 2.866418 | 0.951508 | 2.931274 | 1.129699 | 7 |
| **27** | 5RF0 | -2.0509 | 1.616171 | 0.283994 | 31.212753 | 15.480229 | 8 |
| **11** | 5RE7 | -0.6158 | 2.147856 | 0.29577 | 40.999004 | 21.265048 | Unique pocket |
| **14** | 5REA | -16.0384 | 2.272991 | 0.701226 | 6.311012 | 2.028778 | Unique pocket |
| **28** | 5RF1 | -9.1914 | 1.933961 | 0.320806 | 3.52 | 1.27251 | Unique pocket |

**ST. 4: Mutation analysis of HIS163 to ALA163.** A mutation analysis was performed on five PDB complexes; 5RGZ, 5RF7, 6W63, 5RG1, and 5RGX. HIS163 was mutated to ALA163 in these complexes to demonstrate the importance of a key residue in the lateral pocket for ligand binding. Stable3 compares the initial wildtype (WT) MMGBSA vs. mutated MMGBSA, and HIS163 contribution (in kcal/mol) vs. ALA163 contribution (in kcal/mol). As seen in this table, PDB 5RGX demonstrated the highest MMGBSA difference of -8.868 kcal/mol and PDB 5RF7 demonstrated the least MMGBSA difference of -1.076 kcal/mol.

| **Index** | **PDB** | **MMGBSA (WT)** | **MMGBSA (Mutant)** | **HIS163 contribution** | **ΔΔG (WT-mutant)** | **HIS163 contribution** | **ALA163 contribution** |
|---|---|---|---|---|---|---|---|
| **45** | 5RG1 | -36.3341 | -32.1617 | -2.34 | 4.172 | -2.34 | -0.08 |
| 54 | 5RGX | -30.0259 | -21.1579 | -3.68 | 8.868 | -3.68 | -0.76 |
| 23 | 5RGZ | -27.6558 | -22.0734 | -3.47 | 5.582 | -3.47 | -0.07 |
| 34 | 5RF7 | -16.1116 | -15.0403 | -3.14 | 1.076 | -3.14 | 0.66 |
| 42 | 6W63 | -34.3563 | -29.0172 | -3.24 | 5.339 | -3.24 | 0.80 |

**ST. 5:** Summary of MMGBSA-NWAT results from last 30ns production simulation, NWAT=0 is the energy scores without explicit water molecules and NWAT=6 represents the energy released by complex when there are 6 explicit water molecules in the $M^{pro}$ receptor.

| Index | PDB | NWAT=0 | NWAT=1 | NWAT=2 | NWAT=3 | NWAT=4 | NWAT=5 | NWAT=6 | Energy change |
|---|---|---|---|---|---|---|---|---|---|
| **1** | 5r7y | -18.1287 | -18.3749 | -19.3506 | -20.1558 | -21.0057 | -22.0588 | -23.1672 | -5.0385 |
| **3** | 5r80 | -12.791 | -13.4153 | -14.545 | -15.7179 | -16.4716 | -17.0831 | -17.7926 | -5.0016 |
| **4** | 5r81 | -18.3853 | -19.7202 | -21.4202 | -22.5298 | -23.3569 | -24.1105 | -24.9361 | -6.5508 |
| **5** | 5r82 | -11.634 | -11.3951 | -11.6064 | -11.9264 | -12.1958 | -12.5321 | -12.9327 | -1.2987 |
| **6** | 5r83 | -15.3785 | -17.0998 | -18.0877 | -18.5331 | -18.7721 | -18.9785 | -19.2964 | -3.9179 |
| **7** | 5r84 | -16.8109 | -17.9342 | -19.0646 | -20.0172 | -20.5821 | -20.8528 | -21.1861 | -4.3752 |
| **8** | 5re4 | -18.4189 | -18.6189 | -18.4756 | -18.3869 | -18.4949 | -18.8364 | -19.2811 | -0.8622 |
| **13** | 5re9 | -20.7185 | -20.0649 | -19.5152 | -19.1598 | -19.0291 | -19.1813 | -19.5292 | 1.1893 |
| **15** | 5reb | -14.6536 | -16.5801 | -18.1079 | -18.9421 | -19.3415 | -19.6317 | -20.1031 | -5.4495 |
| **23** | 5rgz | -27.6665 | -29.5802 | -30.0991 | -30.553 | -31.1224 | -31.8197 | -32.5853 | -4.9188 |
| **25** | 5rhd | -13.6331 | -17.7615 | -19.2188 | -19.5682 | -20.1527 | -21.0507 | -22.0846 | -8.4515 |
| **33** | 5rf6 | -18.2132 | -18.3084 | -18.3689 | -18.3459 | -18.4431 | -18.7389 | -19.2025 | -0.9893 |
| **34** | 5rf7 | -16.124 | -17.5443 | -18.1157 | -17.8168 | -17.5766 | -17.5217 | -17.6513 | -1.5273 |
| **41** | 5rfe | -21.9428 | -22.9403 | -23.9376 | -24.2479 | -24.3416 | -24.5049 | -24.8203 | -2.8775 |
| **42** | 6w63 | -34.104 | -34.7496 | -35.5315 | -36.2082 | -36.7294 | -37.0549 | -37.418 | -3.314 |
| **45** | 5rg1 | -36.3457 | -37.7238 | -39.2366 | -40.1227 | -40.6586 | -41.207 | -41.8411 | -5.4954 |
| **46** | 5rgh | -12.1315 | -11.9458 | -12.1232 | -12.4275 | -12.5795 | -12.8112 | -12.9963 | -0.8648 |
| **47** | 5rgi | -23.8187 | -22.9437 | -22.5315 | -22.3715 | -22.36 | -22.5407 | -22.8702 | 0.9485 |
| **49** | 5rgk | -14.5308 | -15.9573 | -16.2735 | -16.1214 | -15.938 | -15.8067 | -15.8598 | -1.329 |
| **53** | 5rgy | -19.1731 | -20.7299 | -21.763 | -22.1552 | -22.0556 | -21.7481 | -21.4977 | -2.3246 |
| **54** | 5rgx | -30.0391 | -29.3442 | -29.0539 | -29.0731 | -29.3262 | -29.8207 | -30.4148 | -0.3757 |
| **55** | 5rh0 | -22.2513 | -24.2944 | -25.9261 | -26.1551 | -26.2571 | -26.632 | -27.203 | -4.9517 |
| **56** | 5rh2 | -24.2783 | -25.9492 | -26.7775 | -26.7428 | -26.7905 | -26.9646 | -27.3465 | -3.0682 |
| **57** | 5rh1 | -25.9447 | -26.8075 | -26.6082 | -26.5304 | -26.7123 | -27.041 | -27.4347 | -1.49 |
| **59** | 5rgv | -22.6573 | -24.2236 | -25.081 | -25.219 | -25.3196 | -25.5493 | -25.8739 | -3.2166 |
| **60** | 5rgw | -22.8069 | -24.3444 | -25.2402 | -25.5859 | -25.7932 | -26.1974 | -26.689 | -3.8821 |
| **62** | 5rh8 | -22.1523 | -22.9963 | -23.5743 | -23.6666 | -23.7845 | -24.1634 | -24.6588 | -2.5065 |

**ST. 6:** The following table summarizes all they ligands in PDB complexes that were run in both the monomeric and dimeric simulations alongwith their calculated MMGBSA binding affinity in both cases as well as the residues that had a significant impact on the binding energy in each case extracted from the pairwise decomposition calculations. Ligands in 5RFA,5RGQ and 5RF0 were present at the dimer interface and as indicated through the binding affinity changes, were stable already in the monomer(5RGQ) or became stable only when simulated in the dimer model (5RF0 and 5RFA). The rest of the compounds were found within proximity of the dimer interface and this includes the active site. While only 5REH(in the active site) was stabilised when simulated in the dimer, the rest of the ligands either had minimal impact from the dimer simulation or even a negative impact in the case of 5RE7.

| PDB ID | Average (Monomer) | Std. Dev. | Average (Dimer) | Std. Dev. | Decomposed residues(monomer) | Decomposed residues (Dimer) |
|---|---|---|---|---|---|---|
| 5rgq | -18.3015 | 4.23 | -21.786 | 3.15 | 4 6 8 295 298 299 | 308 310 429 593 397 600 601 |
| 5rfa | -10.8007 | 4.05 | -15.9308 | 2.49 | Weak binding | 123 310 454 456 |
| 5reh | 0.0032 | 0.02 | -14.4416 | 3.01 | Weak binding | 468 465 444 442 |
| 5rez | -11.56 | 5.3 | -11.8501 | 4.02 | Weak binding | Weak binding |
| 5rf0 | -2.0509 | 3.18 | -10.9406 | 4.26 | Weak binding | Stable |
| 5r7z | -7.5255 | 6.07 | -7.8501 | 4.51 | Weak binding | Weak binding |
| 5re7 | -0.6158 | 1.84 | -7.5187 | 3.46 | Weak binding | 153 294 |
| 5rf9 | -8.414 | 2.37 | -7.3238 | 5.46 | Weak binding | Weak binding |
| 5rgj | -6.6131 | 2.96 | -6.1587 | 4.54 | Weak binding | Weak binding |
| 5re8 | -0.1094 | 0.58 | -3.5218 | 3.04 | Weak binding | Weak binding |
| 5rf1 | -9.1914 | 2.39 | -3.4376 | 2.91 | Weak binding | Weak binding |
| 5rf3 | -0.1652 | 0.87 | -2.5144 | 2.49 | Weak binding | Weak binding |
| 5rf2 | -0.0124 | 0.78 | -0.1581 | 1.03 | Weak binding | Weak binding |

**SFig. 1 –** RMSD for 6M2Q apo structure, organized in domains.

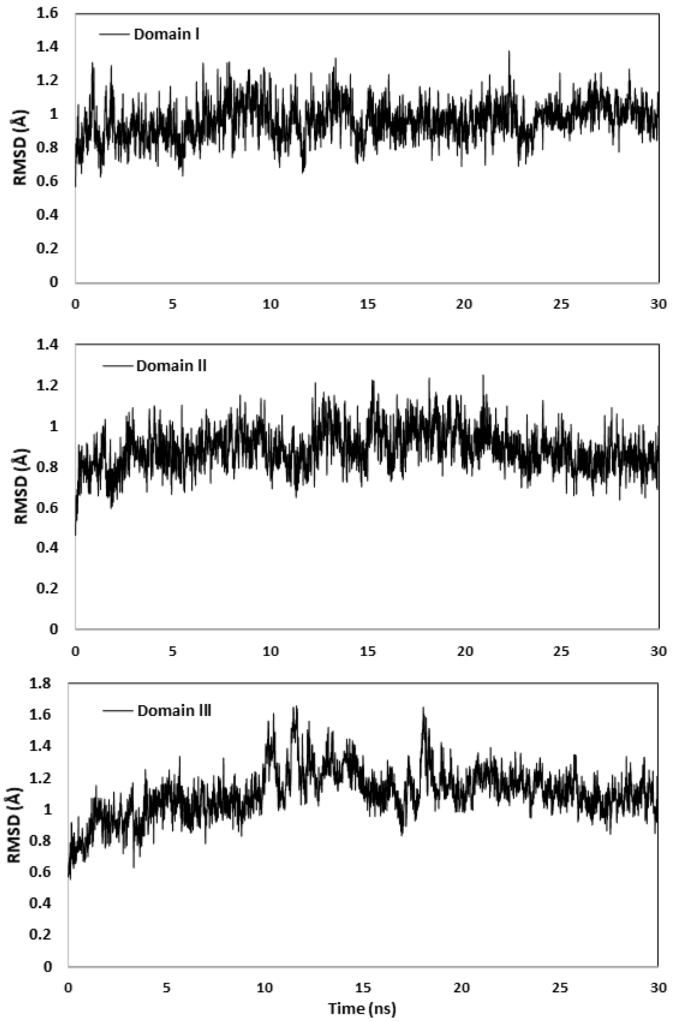

**SFig. 2** Different interactions like hydrogen bonds within protein residues (*), salt bridges (**) and water molecules (***) are examined. Details for each interaction are shown in SFig. 3.

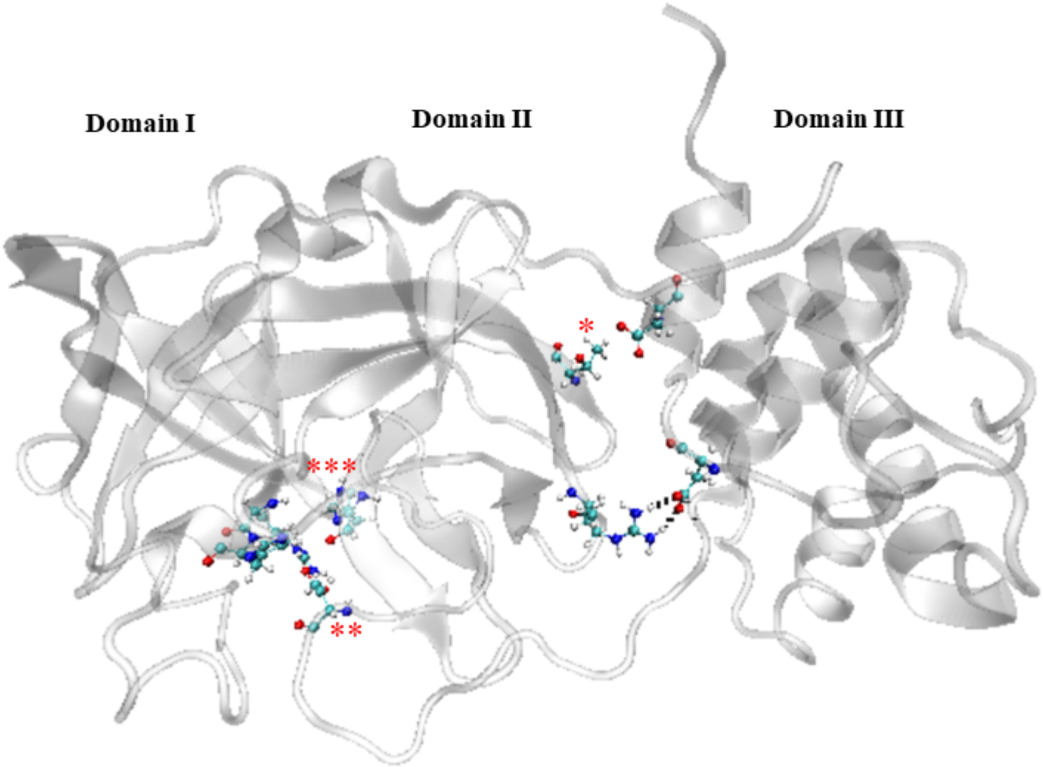

**SFig. 3** – Interactions in apo 6M2Q, hydrogen bonds between THR111, ASP295 (a), and salt bridges between ARG131, ASP189(b), and between ARG40, ASP289 (c).

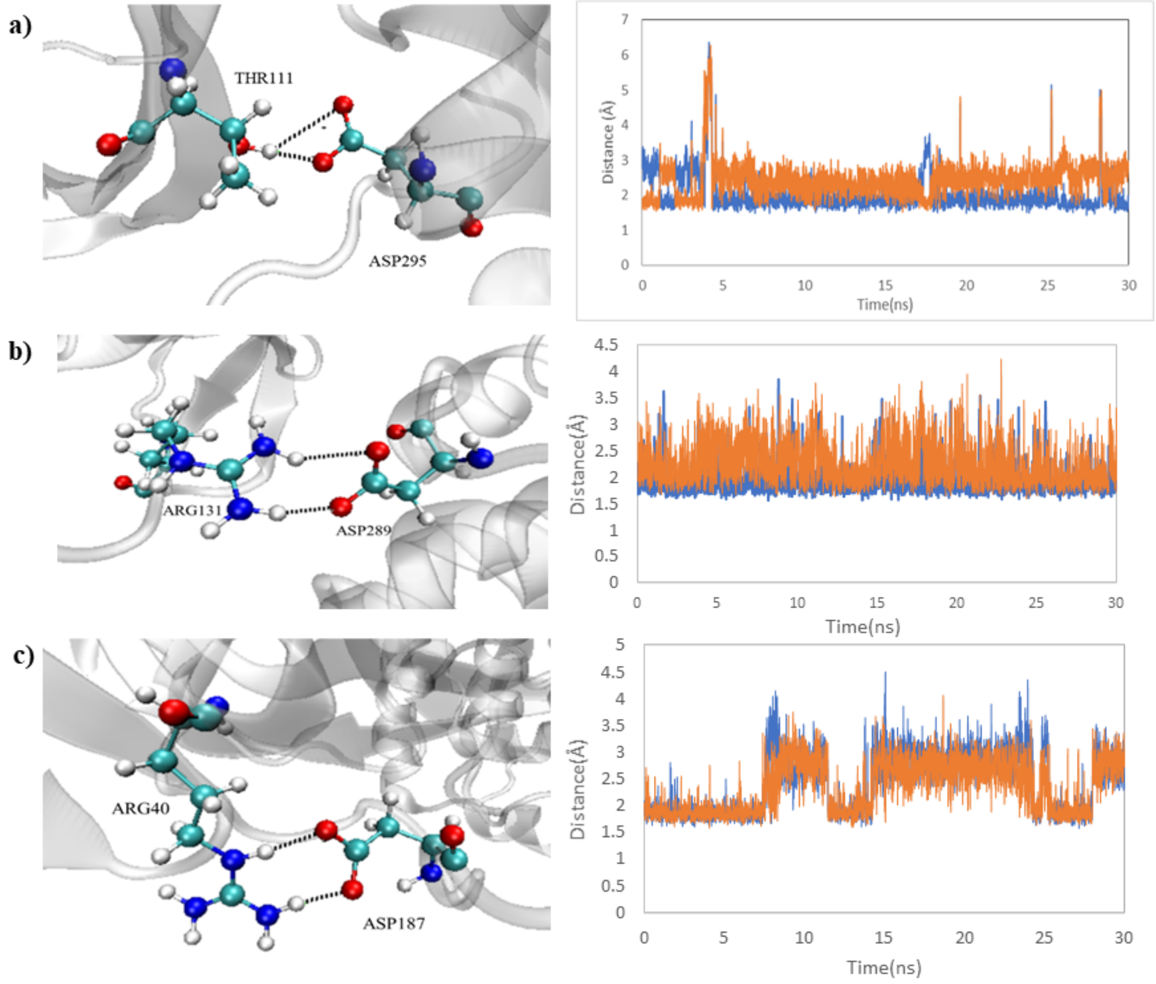

**SFig. 4 Ligand (US7) pose of PDB ID 5RHD (a) and its RMSD plots (b).** US7 represented in red and blue to show ligand position in beginning and end of production simulation (a). RMSD plots were provided to show stable protein backbone and blue ligand represents favorable pose throughout MD, as RMSD fluctuates around 3.5 Å for about 27 ns (b).

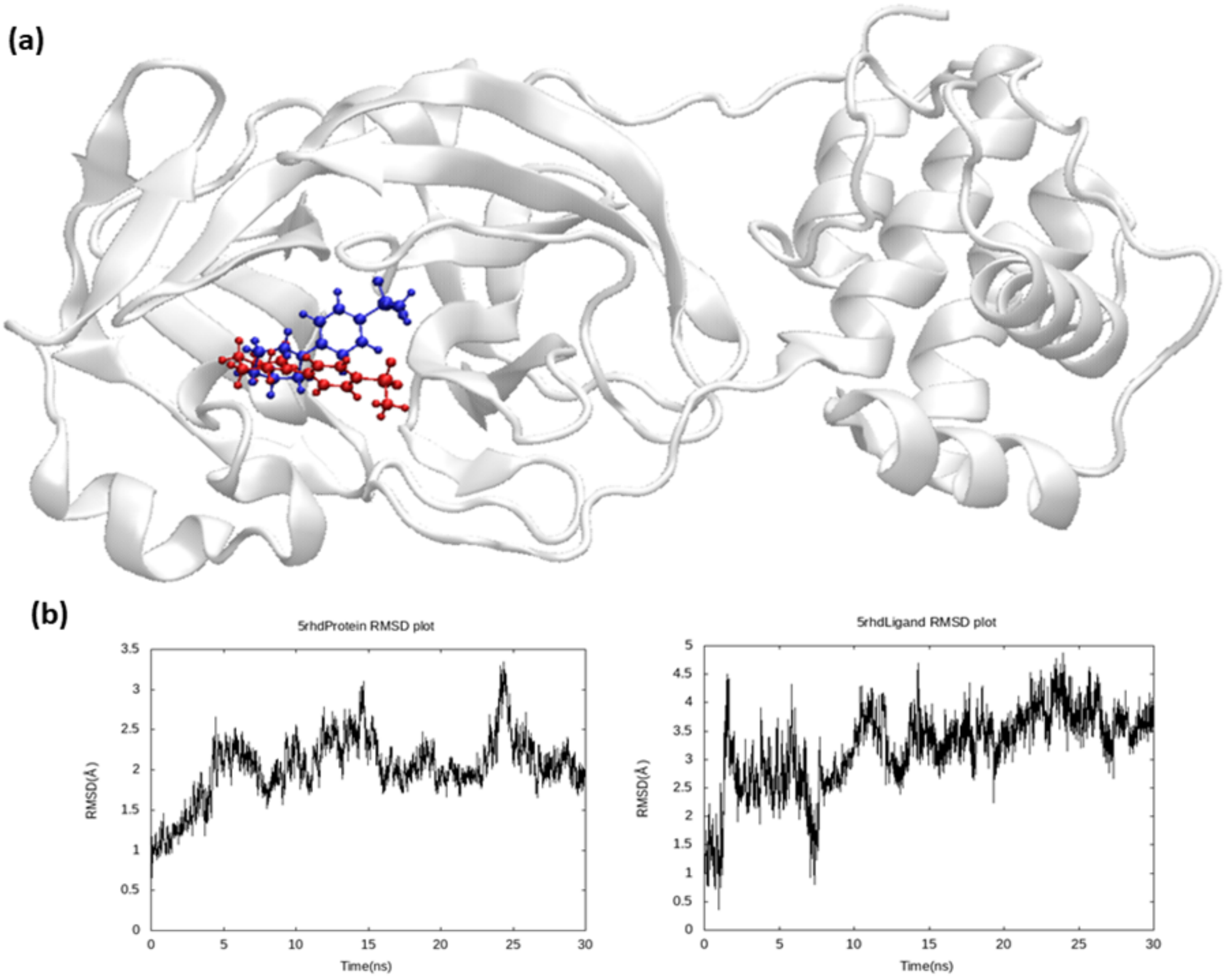

**SFig. 5** The following graphs represent the average ligand RMSD of the compounds in Pockets 2-6 as observed in the 30ns simulation. Details of ligands in pockets 7,8 and unique pockets are outline in Stable 3 above.

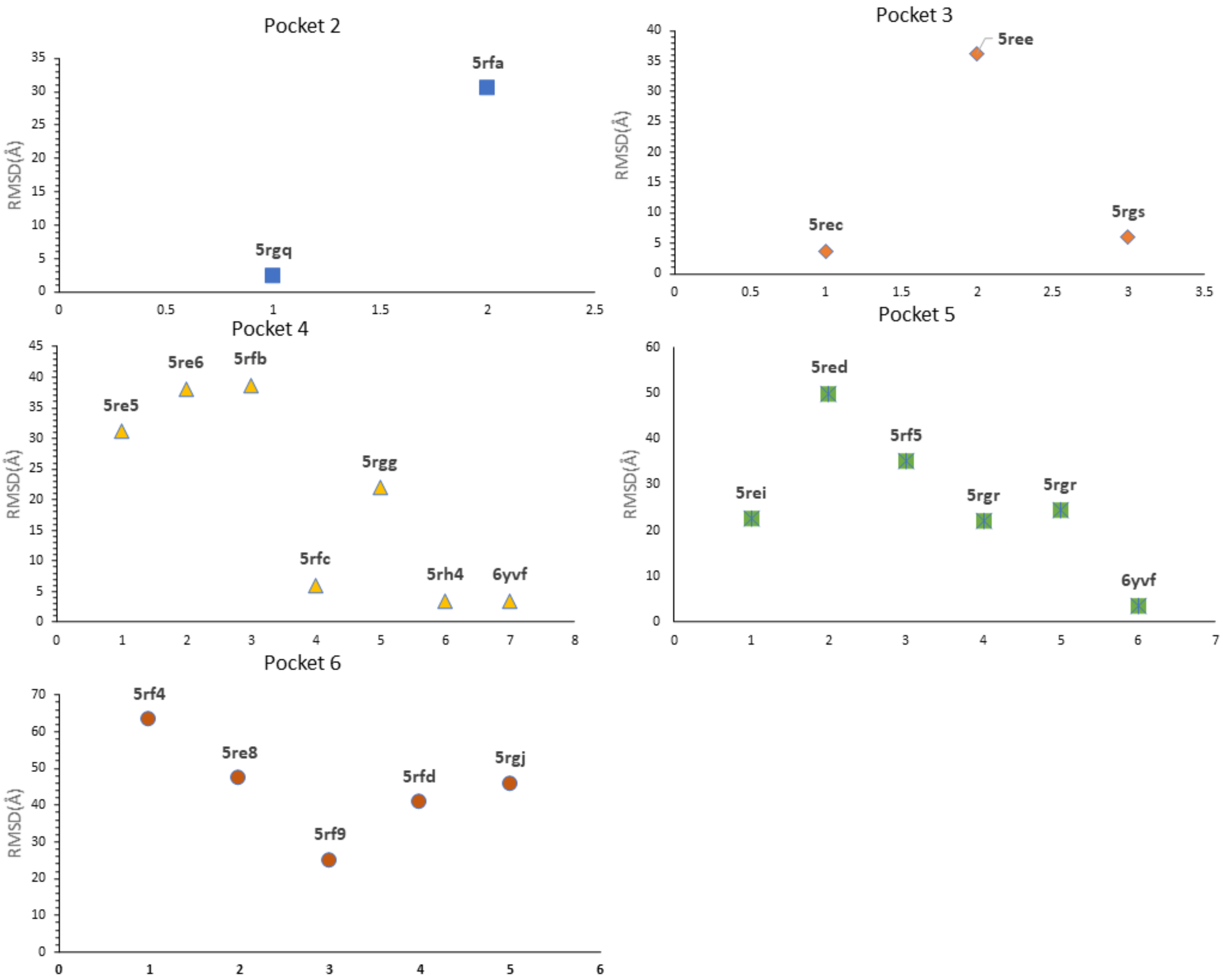

**SFig. 6.** Graph representing the MMGBSA binding affinities calculated in terms of free energy (kcal/mol)for 34 non-covalently bound ligans that are bound the the Main protease in an unstable manner and ligands bound to other sites other than the orthosteric site.

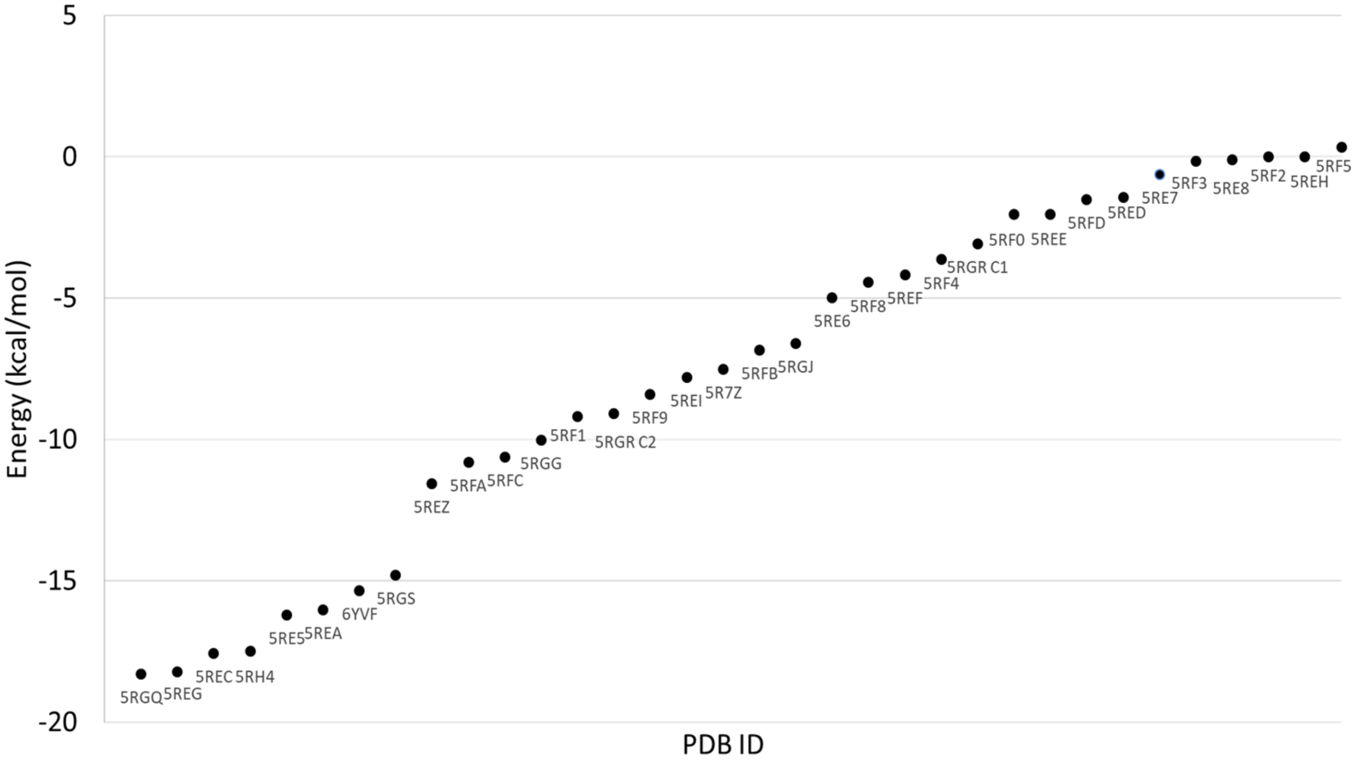

**SFig. 7. Diagrams representing the MD behavior of ligand in complex 5RGH(b) (Binding affinity -12.1 kcal/mol) and some of the interacting residues (a).** The different confirmations of the ligand shown as a function of time in (b) range from 0ns (red) to 29ns(blue) with white and pink showing intermediate poses, clearly highlighting the exhaustive motion around the active site while being anchored near the 180's loop through fluoride interactions with surrounding residues.

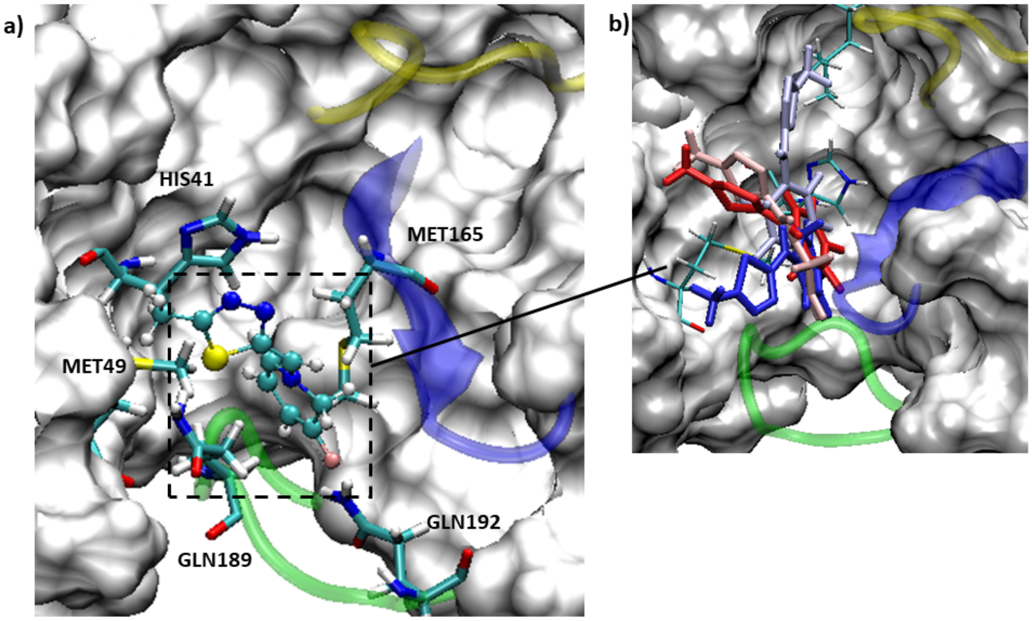


**SFig. 8.** Diagram showing the ligand in complex 5R82 with a binding affinity of -11.6 kcal/mol changing confirmations frequently in the binding site which are represented in the same diagram and coloured according to the timescale from 0ns (red) to 29ns (blue) and the colours in between. This fluctuating behavior explains why the ligand in 5R82 has relatively poor binding affinity.

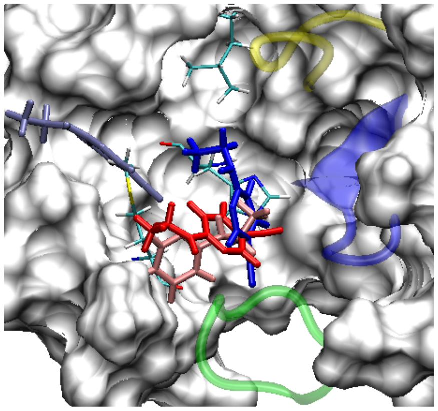

**SFig. 9. Another high scoring ligand, in complex 5RH3 with a binding affinity of -22kcal/mol. The interaction diagram in (a) shows the key residues in stick representation which impact the overall binding affinity of the ligand as seen in the pairwise decomposition graph in (b). Stable hydrogen bonds with GLU166 and HIS163 shown in (c,d).** These hydrogen bonds are clearly stable and therfore instrumental in stabilising this ligand in addition to the Van der Waals interactions with MET49, GLN189, MET165 and ASN 142 seen through the stick representations in (a) and through their energy impacts in(b).

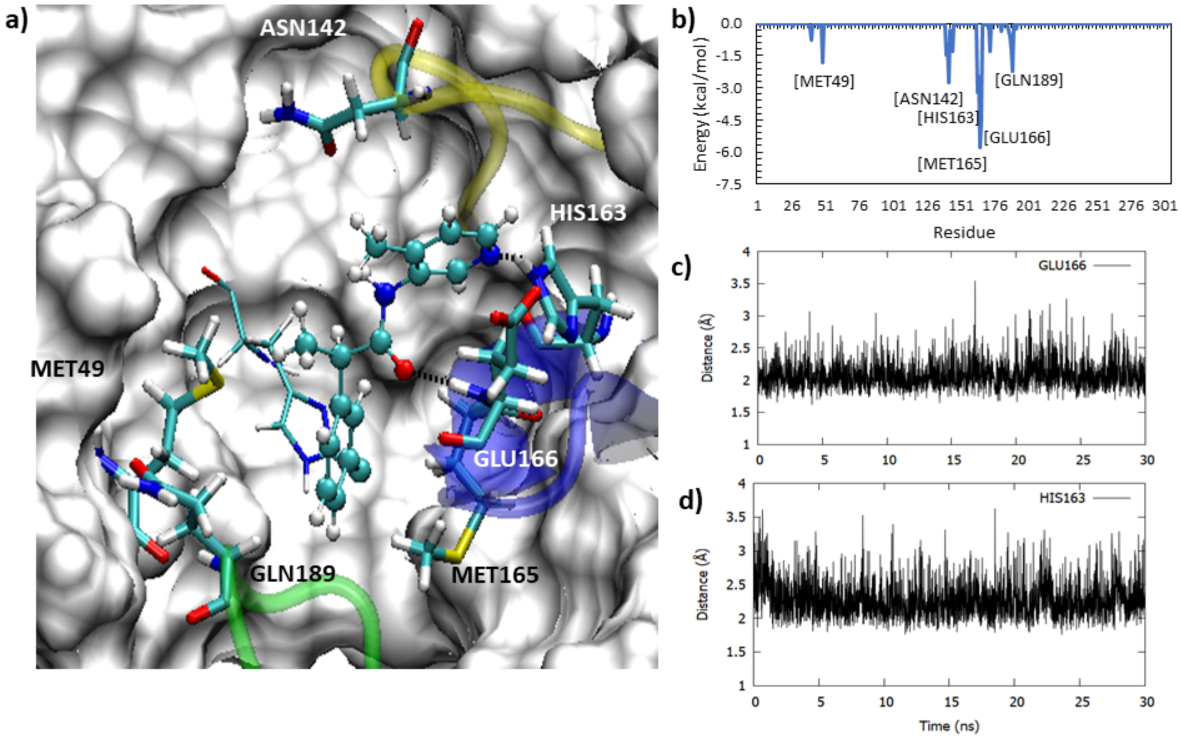

**SFig. 10. Diagram representing the predominant binding pose of the ligand in 5R81(a) with a binding affinity of -18.35 kcal/mol and associated energy impacts from key residues in the decomposition graph (b) as well as hydrogen bond evolution plot with GLN192(c).** The ligand forms Van der Waals interactions with three of the four main regions of the active site as seen in the interaction diagram and in the energy, peaks corresponding to these residues in (b) namely HIS41, MET49, MET165, GLU166, GLN189. The oxygens of the sulfonamide group form a combined stable hydrogen bond with the amino group of GLN192 while the terminal amino moiety forms hydrogen bonds with THR190 in the latter half of the simulation.

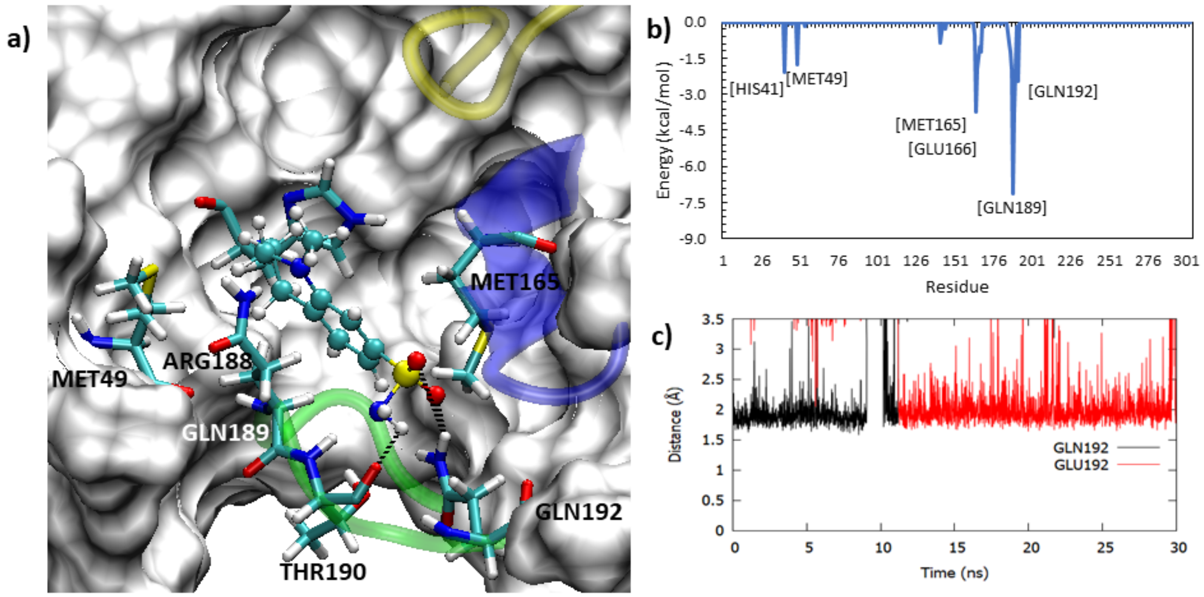

**SFig. 11. Interaction diagram of ligand in complex 5RF7 (a) alongwith its pairwise decomposition graph in (b) and impactful hydrogen bonding evolution plots in (c) and (d).** The ligand has a binding affinity of and is clearly occupying the lateral pocket and forming a stable hydrogen bond with HIS163 (a,c) but the rest of the ligand is not stabilised evidenced by fluctuating hydrogen bonds with GLY143, ASN142 and GLU166 as well as the lack of interactions with other main regions of the active site.

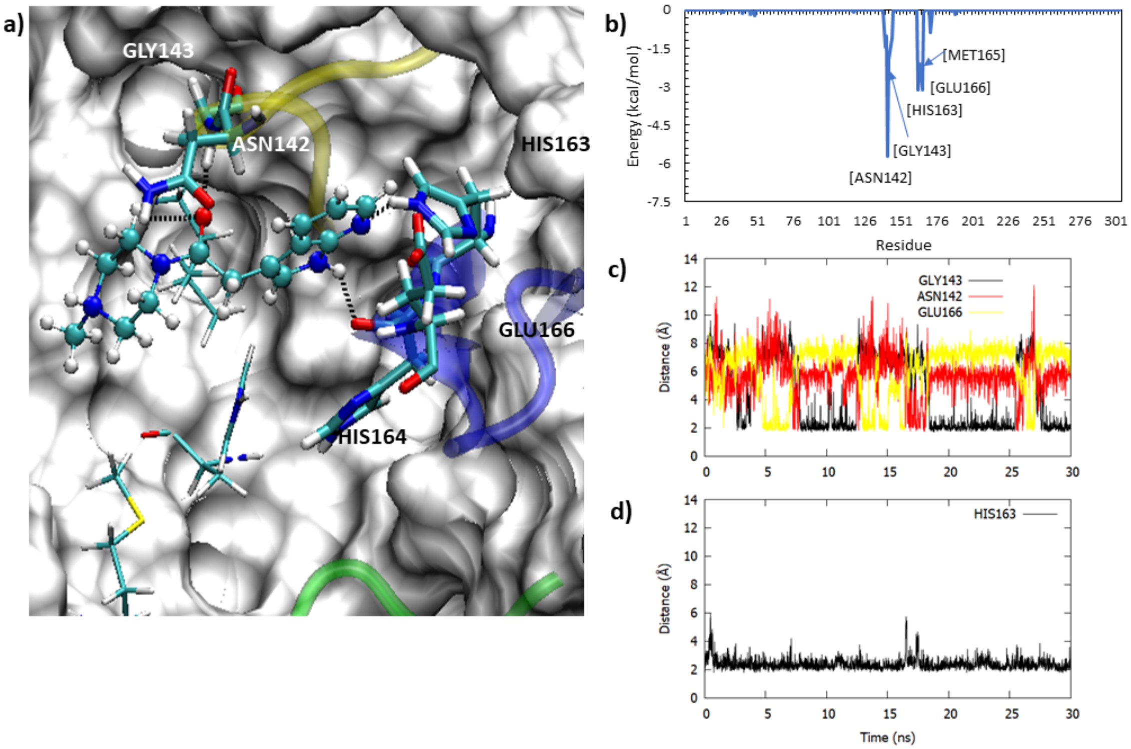

**SFig. 12. Wildtype 5RGX complex interactions (a), 5RGX HIS163ALA mutant complex interactions (b), and pairwise residue decomposition graph depicting residue contribution in 5RGX wildtype and mutant complexes (c).** Representative snapshots of key interactions near the pyridine ring are shown in Fig.X(a) and (b). The wildtype complex forms strong electrostatic interactions with HIS163, GLY143, SER144, and CYS145. The mutant complex loses electrostatic interactions with the pyridine ring and the double bonded oxygen atom is shifted to form transient interactions with CYS145 and GLU166. The difference in residue contribution to overall MMGBSA binding affinity is shown via the wildtype and mutant pairwise decomposition graphs where the residue denoted by an asterisk (*) represents HIS163ALA.

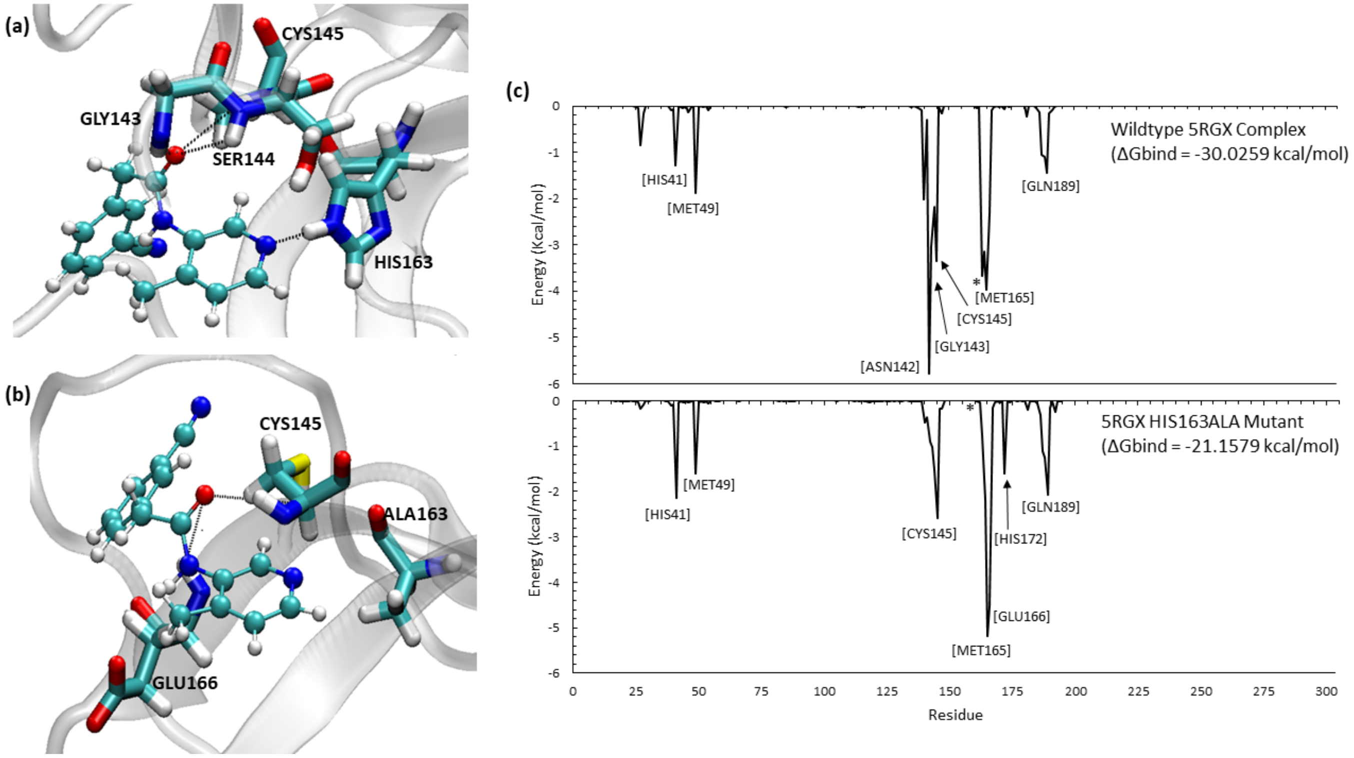

**SFig. 13. Representative snapshot of the 5RGZ HIS163ALA mutant complex at 0 ns and 15ns (a), and pairwise residue decomposition graph depicting residue contribution in 5RGZ wildtype and mutant complexes (b).** The initial binding poses of the 5RGZ HIS163ALA mutant complex (red) reveals the pyridine ring located close to ALA163 in the lateral pocket. After 15 ns, the pyridine ring rotates out of the binding pocket (blue), greatly decreasing its MMGBSA binding affinity. The difference in residue contribution to overall MMGBSA binding affinity is shown via the wildtype and mutant pairwise decomposition graphs where the residue denoted by an asterisk (*) represents HIS163ALA.

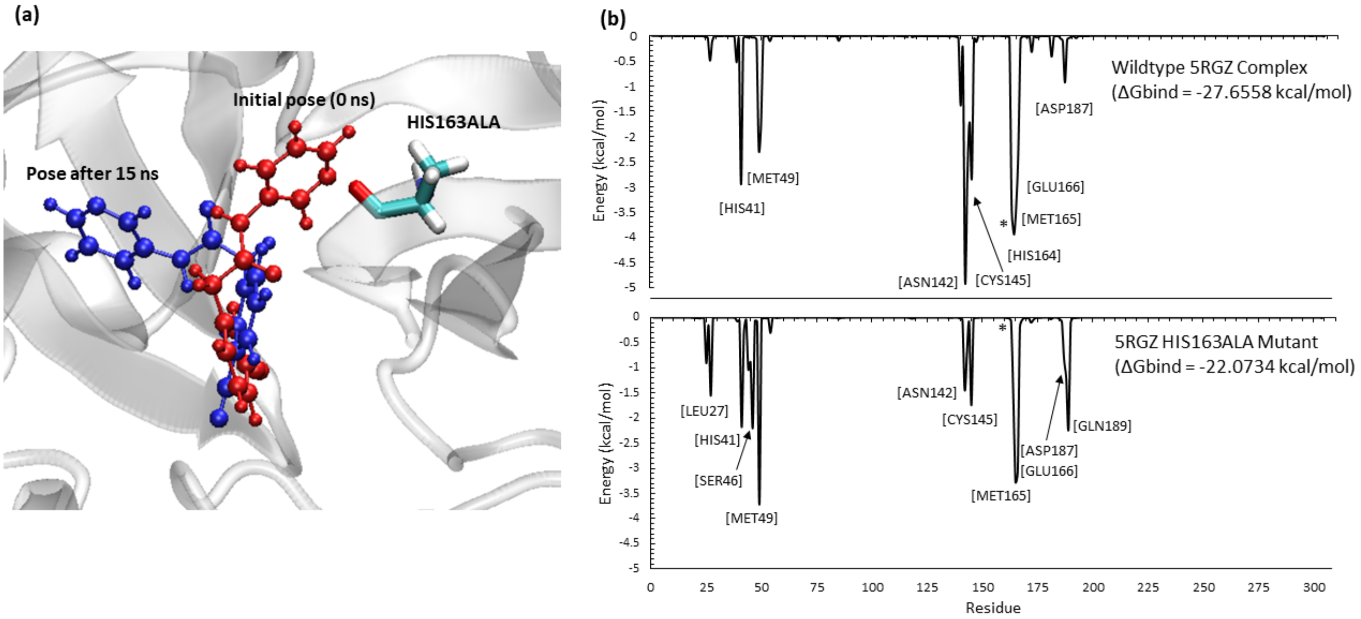

**SFig. 14. Wildtype 5RG1 complex interactions (a), 5RG1 HIS163ALA mutant complex interactions (b), and pairwise residue decomposition graph depicting residue contribution in 5RG1 wildtype and mutant complexes (c).** The wildtype 5RG1 complex shows stable interactions with key residues HIS163, GLU166, and GLN189. In the HIS163ALA 5RG1 mutant complex, the HIS163 strong, electrostatic interaction is replaced by an additional, but transient, electrostatic interaction with GLU166. The alanine mutation also causes the GLN189 to be lost. The difference in residue contribution to overall MMGBSA binding affinity is shown via the wildtype and mutant pairwise decomposition graphs where the residue denoted by an asterisk (*) represents HIS163ALA.

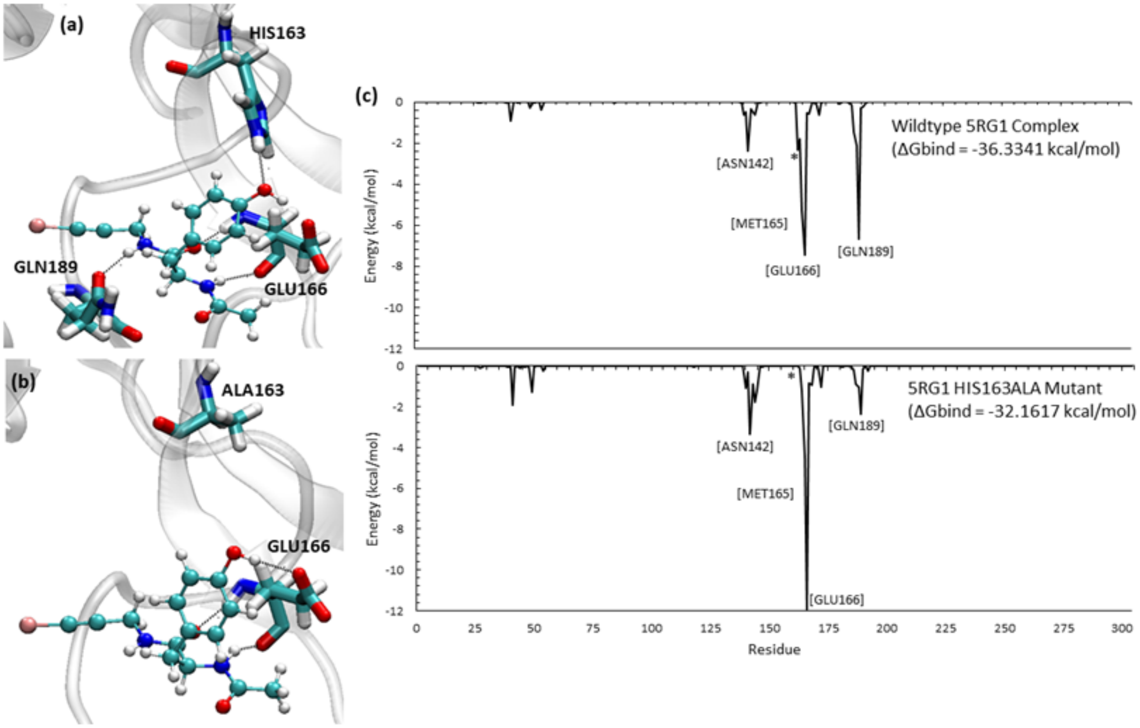

**SFig.15. Two representations of key wildtype 6W63 complex interactions (a), 6W63 HIS163ALA mutant complex interactions near pyridine ring (b), and pairwise residue decomposition graph depicting residue contribution in 6W63 wildtype and mutant complexes (c).** The wildtype 6W63 complex contains a large ligand that forms many interactions with different residues to maintain its position in the binding site. Some of these key residues include HIS41, ASN142, GLY143, HIS163, and GLU166. Due to its large ligand size, the HIS163ALA 6W63 mutant complex is still relatively stable, but the pyridine ring in the lateral pocket and nearby double-bonded oxygen atoms do undergo some changes in the interactions they form with the protein. For example, the nitrogen atom of the pyridine ring loses its stable interaction with HIS163 to instead form an electrostatic interaction with CYS145. The difference in residue contribution to overall MMGBSA binding affinity is shown via the wildtype and mutant pairwise decomposition graphs where the residue denoted by an asterisk (*) represents HIS163ALA.

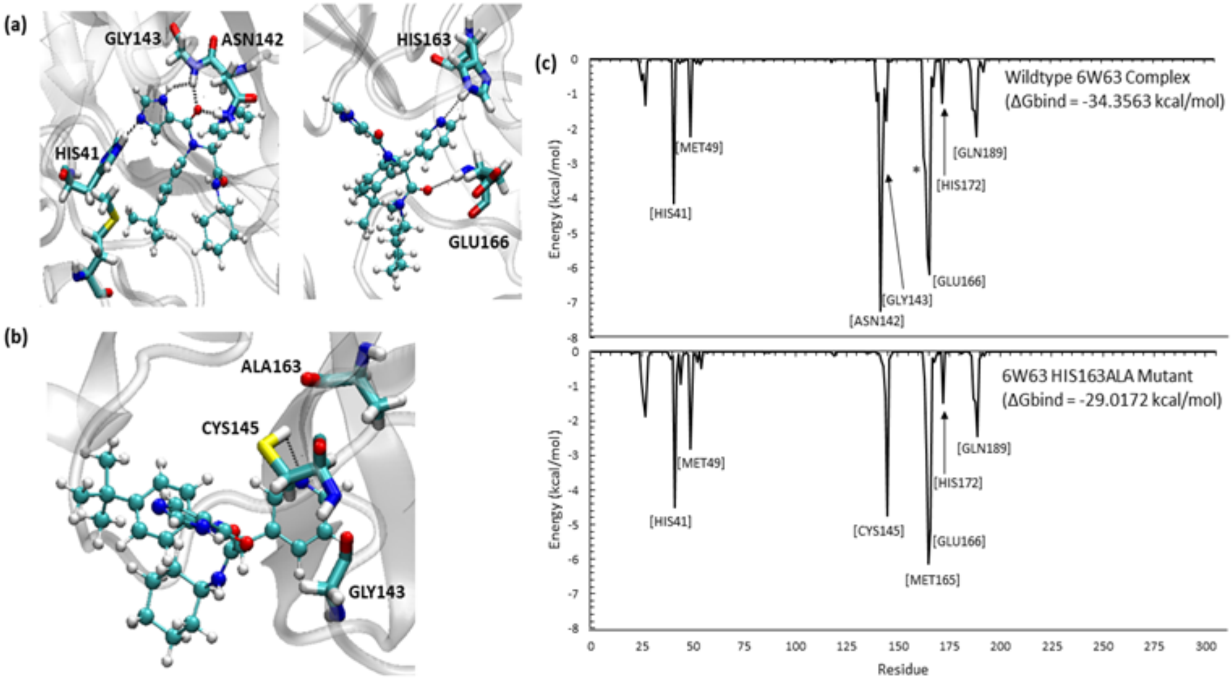

**SFig. 16. Wildtype 5RF7 complex interactions (a), 5RF7 HIS163ALA mutant complex interactions (b), and pairwise residue decomposition graph depicting residue contribution in 5Rf7 wildtype and mutant complexes (c).** The 5RF7 complex contains a unique 7-azaindole double-ring structure positioned in the lateral pocket. This enables two strong hydrogen bonds with HIS163 and HIS164 to be present in the wildtype 5RF7 complex, shown in Fig.Z.a. When the complex is mutated from a histidine to alanine at reside 163, the hydrogen bond with the 6-membered ring of the 7-azaindole group is lost, but the hydrogen bond between HIS164 and the 5-membered ring is maintained. This interaction stabilizes the HIS163ALA mutant complex in the lateral binding pocket and reduces the MMGSBA difference between the wildtype and mutated complexes. The difference in residue contribution to overall MMGBSA binding affinity is shown via the wildtype and mutant pairwise decomposition graphs where the residue denoted by an asterisk (*) represents HIS163ALA.

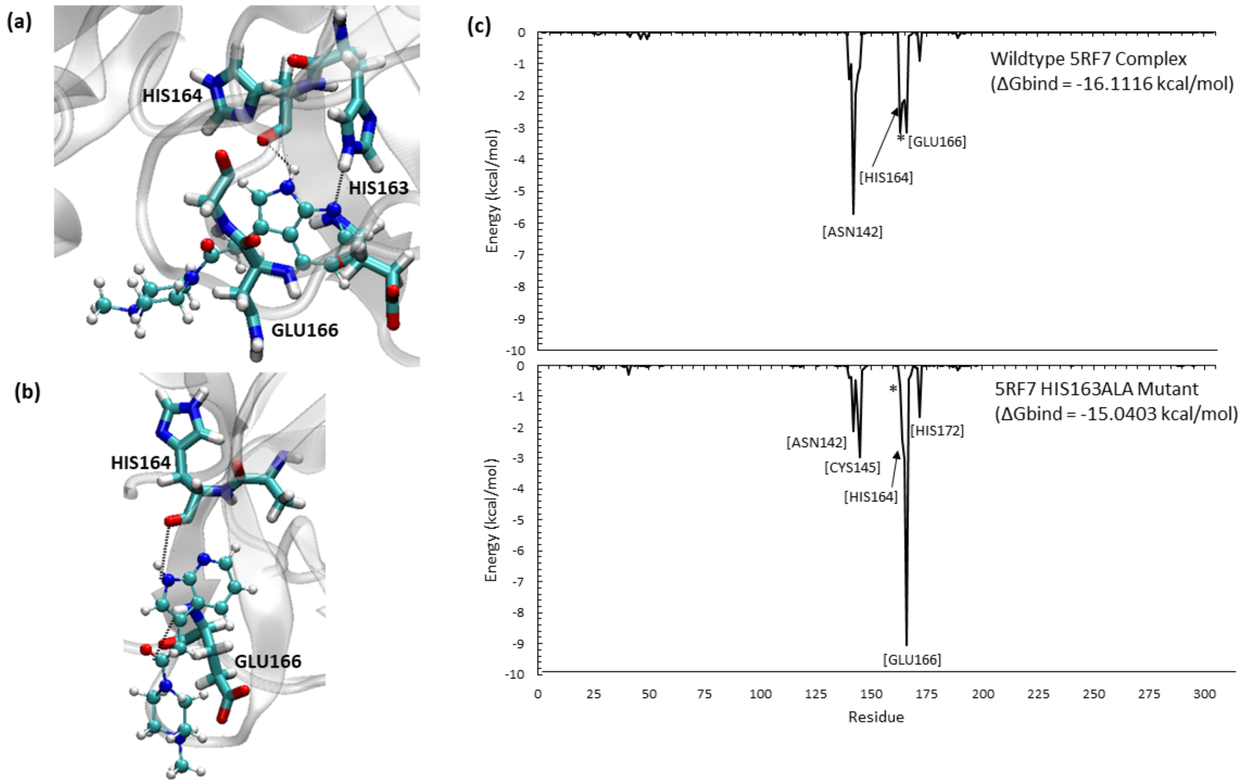

**SFig. 17. Some unfavourable NWATER interactions in complexes showing a negative energy impact of water.** 2-3 water forms unfavorable interaction with benzene ring of 5RGX and 5RE9 at polar region of $M^{pro}$ (e-f).

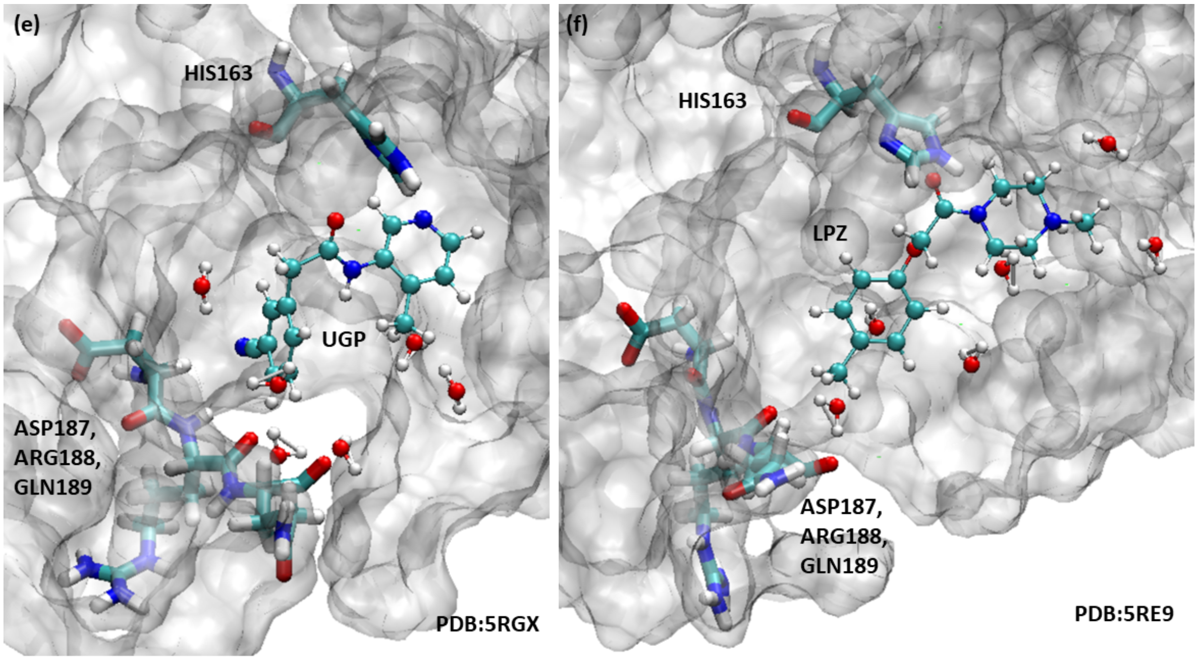

**SFig. 18. Binding Site representations and RMSD for 11 of the 13 ligands simulated in the M$^{pro}$ dimer model.** The following 11 figures are titled with the original crystal structure complex and they showcase the initial binding site of each compound in the dimer in (a) of each figure and the unaligned ligand RMSD throughout the 30ns timescale for both the monomer and dimer simulations are compared in the graphs shown in (b). It can be clearly seen that some ligands definitely benefit from the dimer model while most ligands in this list have not had a significant impact and have egressed from the surface of M$^{pro}$ in both the models.

5R7Z Complex

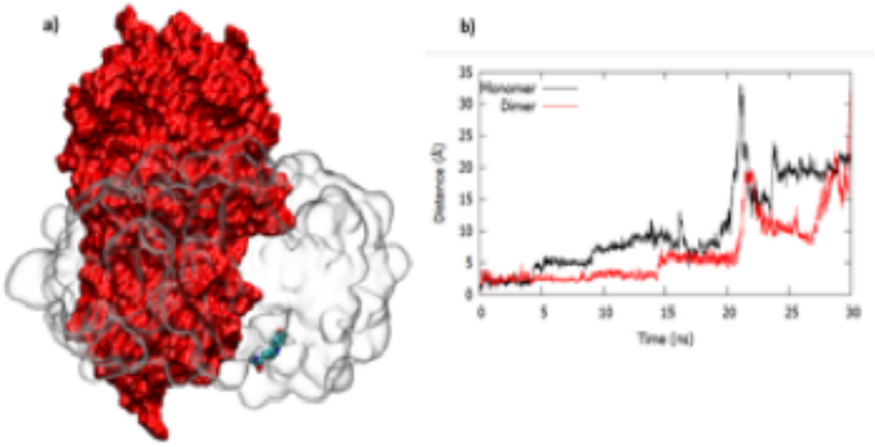


5RF0 Complex

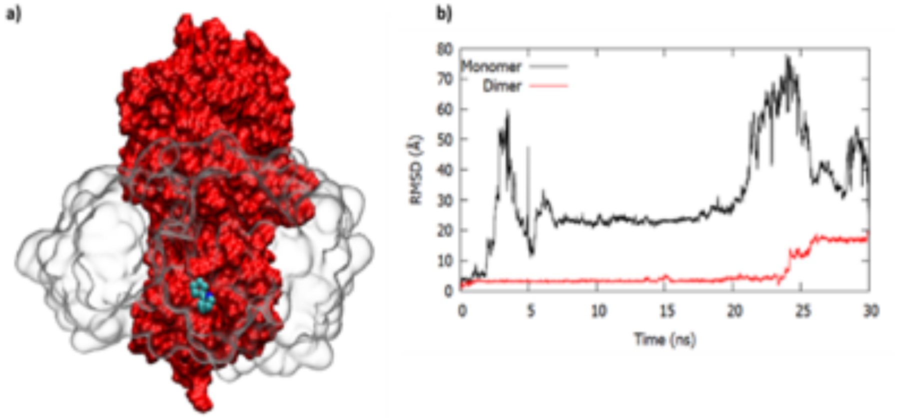

5RE8 Complex

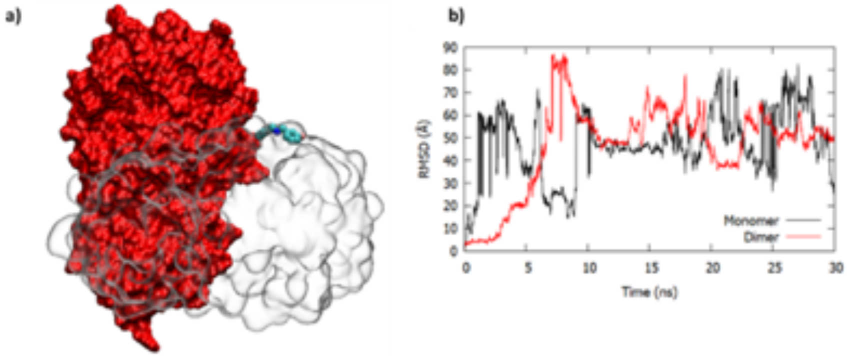


5RE7 Complex

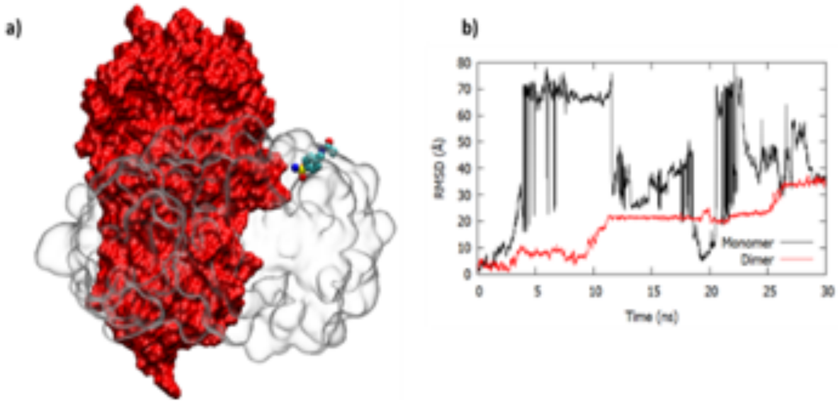


5REZ Complex

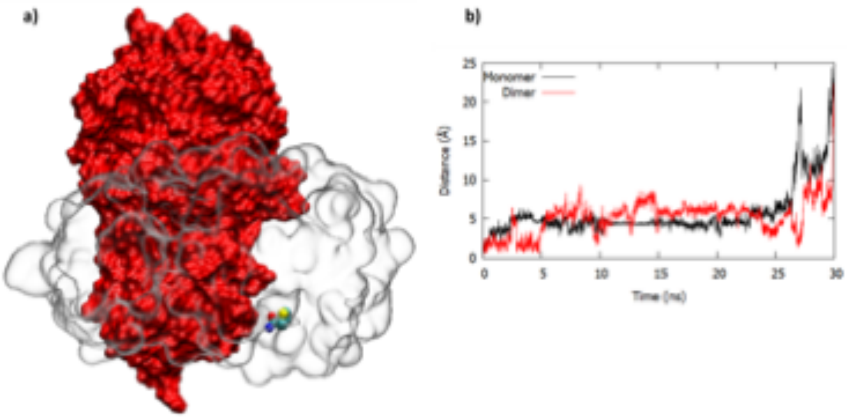

#### 5RF1 Complex

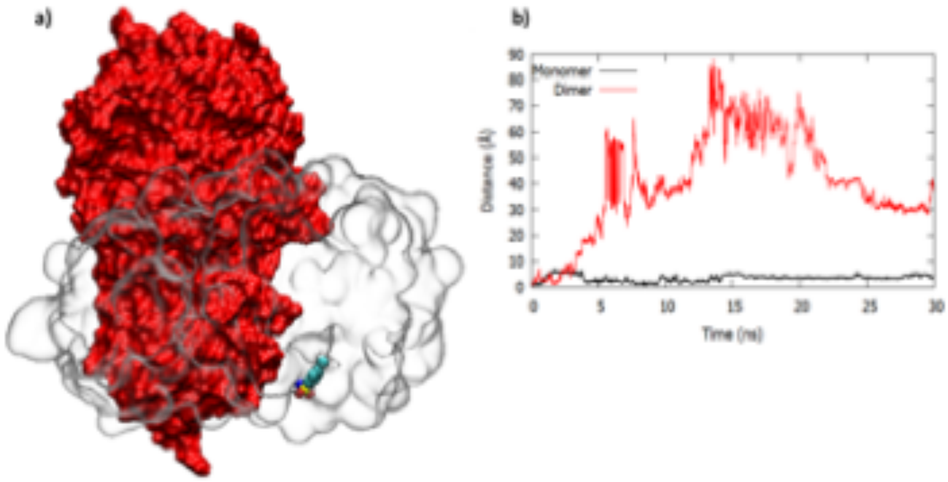


#### 5RF2 Complex

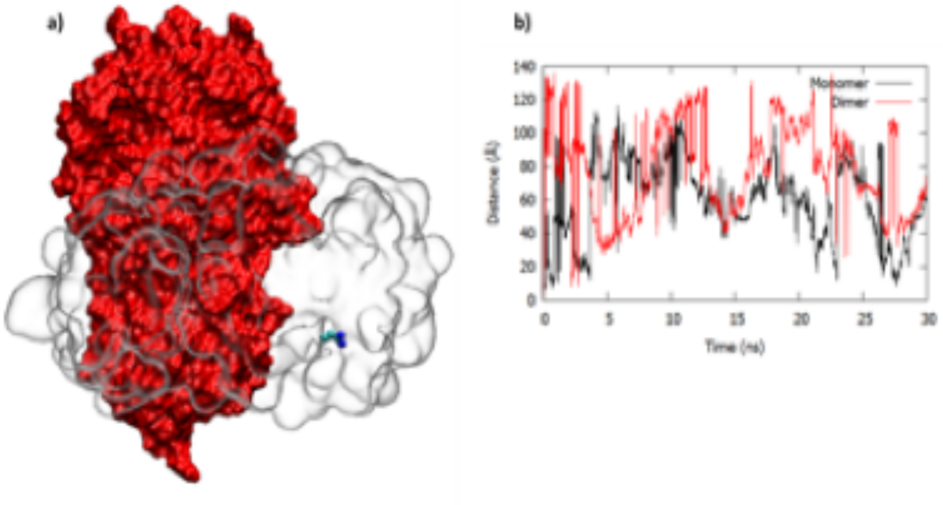

#### 5RF3 Complex

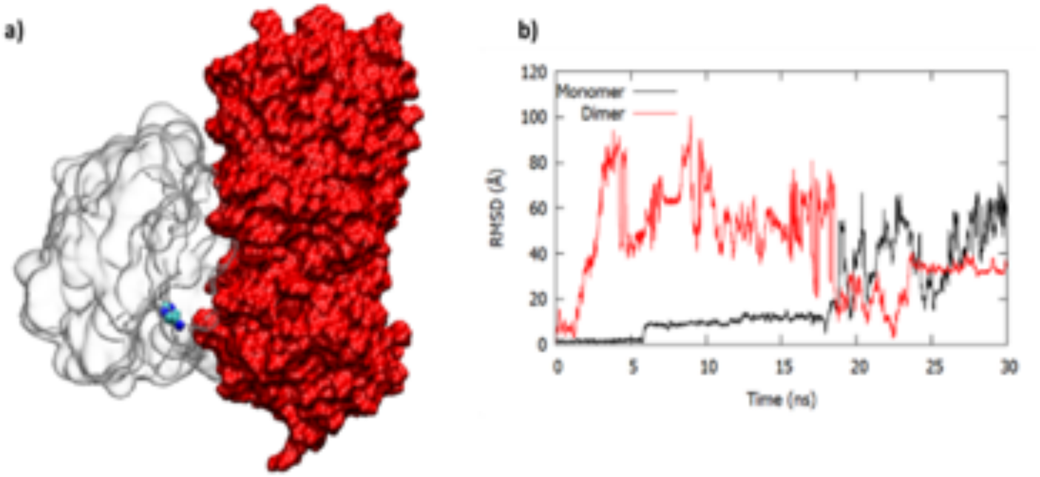


#### 5RF9 Complex

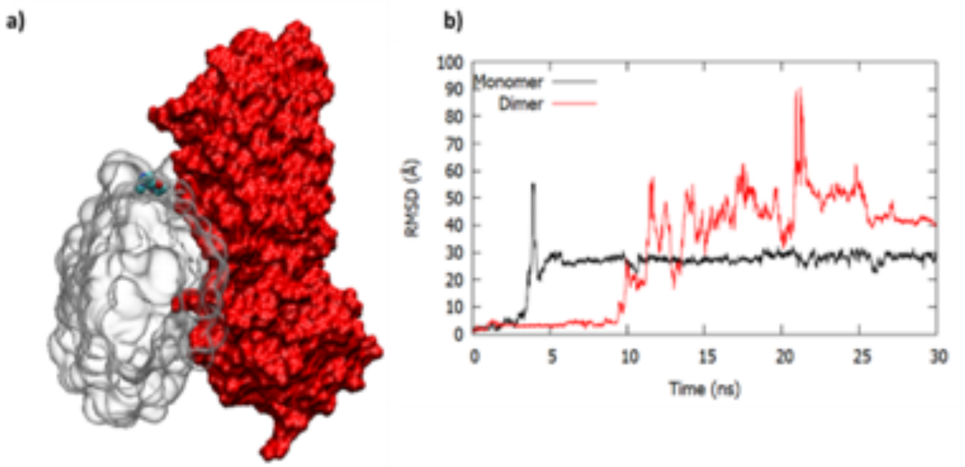

5RGJ Complex

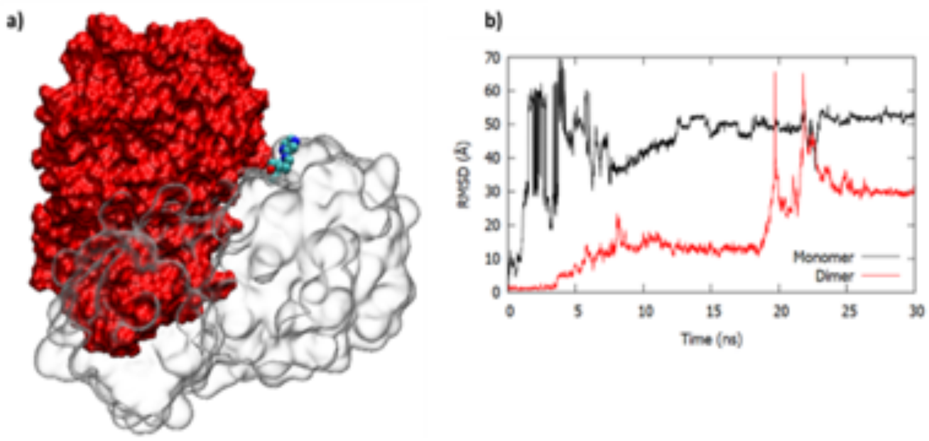


5RGQ Complex

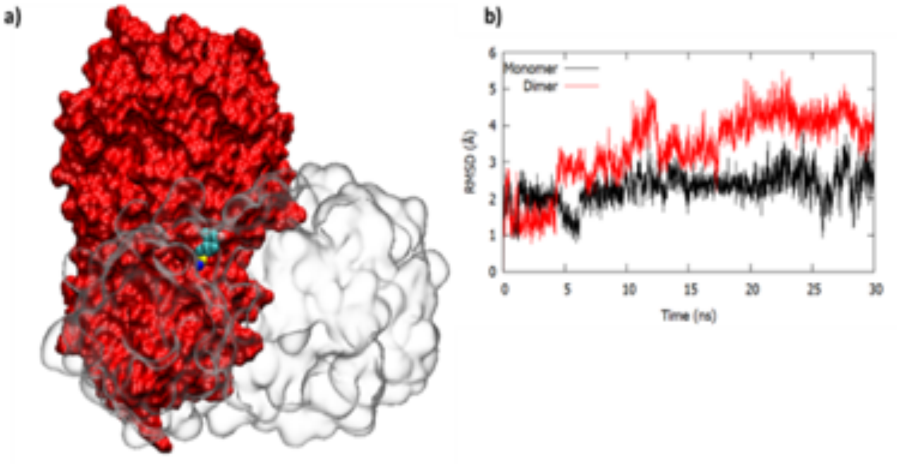
